# Distinct cortical profiles underlie the common reportability of thought-free experiences

**DOI:** 10.1101/2025.10.14.681984

**Authors:** Paradeisios Alexandros Boulakis, Anikó Kusztor, Naotsugu Tsuchiya, Thomas Andrillon, Athena Demertzi

## Abstract

Mind blanking (MB) refers to moments of seemingly no reportable thought content, challenging the view of a constant thought-full mind. What remains unclear is which brain modes can lead to MB and why. Here, we aimed at delineating the nuances of MB by combining EEG–fMRI with experience-sampling of thought content and self-reported vigilance. Using a dynamic framework, we show that MB is linked to a “rich” fMRI connectivity pattern of regional anticorrelations, while low vigilance is linked to a “simple” pattern of overall positive connectivity. Put together, an interaction appears: when people are alert, connectivity around MB resembles the hyperconnected pattern, indicating that the neural correlates of MB depend on vigilance levels. This hyperconnected pattern further correlated with EEG slow-wave-like activity, tying MB’s topology to sleep-like electrophysiology. We conclude that MB’s neurophysiological correlates vary with vigilance, and a more refined characterisation of the neural correlates of thought-free states exists.

## INTRODUCTION

Wakeful life is experienced as being full of thoughts. The content of these thoughts can reflect either external demands and goal-oriented tasks, such as problem-solving or responding to the environment, or can arise spontaneously in the absence of external cues, as seen in daydreaming, mind wandering, or internally generated thoughts ^1–3^. Considering this affluence in mental content people often assume that their mind is perpetually “thought-full”, continuously alternating between externally and internally generated contents. Yet, this view is challenged by occasional reports, such as “I have no thoughts”, “I am not sure what I was thinking about”, and “I forgot what I had in mind”, among others. These types of episodes are known as mind blanking (MB) ^4–6^. More formally, MB is defined as the absence of conscious awareness and a feeling of an empty mind ^7^. Operationally, an MB occurs when participants have no reportable content or are unable to report any content ^5^. This definition is in line with our previous work but contrasts with other approaches of MB, including those focusing more specifically on the absence of thoughts or on voluntary MB (see Andrillon et al. ^4^ for a review).

Our understanding of how MB episodes emerge has only now started to get illuminated ^4^. So far, experimental work indicates that MB stands in behavioural and neural contrast to mental states associated with reportable thought content ^7–11^. During wakefulness, MB reports occur infrequently (∼5-10% of the time), both in resting conditions ^9,10^ and during task-engagement ^7,12,13^. At the same time, the frequency of MB reports also seems to depend on arousal levels, as they increase after sleep deprivation ^10^ and during subjectively felt lower vigilance ^7,14^.

The indication that low arousal is linked to the absence of reportable thought content is further supported by neuroimaging and electrophysiological findings. Using fMRI, we previously examined how functional connectivity organises around content-full mental state reports and MB ^9^. We found that, under task-free conditions, MB was associated with a connectivity pattern characterised by a global inter-areal synchronisation. Considering that similar increases in whole-cortex functional connectivity have also been reported during NREM sleep in humans ^15^ and under isoflurane anaesthesia in animals ^16^, this overall positive connectivity has been interpreted as an indication that low cortical arousal underlie MB reports. The low arousal stance was further supported by the finding that the amplitude of the fMRI global signal (GS) was higher around the time of MB reports ^9^. The GS refers to the average signal intensity across all voxels in the grey matter ^17^. As the amplitude of the GS was shown to positively correlate with delta oscillations in the EEG frequency band and to decrease after ingestion of caffeine ^18^, it was considered a proxy of cortical arousal. Therefore, with regard to MB, the co-occurrence of fMRI hyperconnectivity and increased GS amplitude points to the possibility of low neurophysiological arousal during MB reports. A direct correlation between fMRI and electrophysiology could strengthen this interpretation.

Indeed, what we know so far about MB’s EEG substrate comes from a separate piece of work, where we have shown that MB reports are preceded by the presence of slow-wave-like (SW) activity during wakefulness, known as “local sleeps” ^7,19–21^. Combining EEG with experience sampling during a sustained attention to response task (SART), we found that attentional lapses were associated with the presence of transient sleep-like SWs, and the properties of these SWs differentiated the two mental reports: steeper SW over posterior electrodes was associated with MB, larger and steeper SW in frontal electrodes was associated with mind wandering reports ^7^. A replication and re-analysis of this paradigm revealed that MB was further associated with higher power in the EEG delta and alpha bands, lower power in beta and gamma and reduced parietal complexity, all indicative of a reduced cortical arousal mode ^22^. It should be noted, though, that the low spatial resolution of EEG limits the identification of the precise networks involved in sleep-like SW and MB.

Overall, MB posits a frontier in the study of spontaneous thinking and consciousness, challenging the long-held assumption of a constantly content-full mind ^4^. In turn, the neural correlates across EEG and fMRI suggest that the wakeful brain may transition to sleep-like states, with a stark impact on conscious experience ^9,22^. In the present study, we combined simultaneous fMRI-EEG recordings with experience sampling to determine the spatio-temporal profile of MB, as measured by multimodal brain activity. We hypothesised that MB-related fMRI connectivity would be organised in an overall positive inter-regional pattern and that this hyperconnected pattern would correlate with EEG SW-like activity. By taking reported vigilance levels into account, we were also open to the possibility that distinct MB fingerprints would emerge, in line with our recent theory that different forms of empty-mindedness may exist ^4^.

## RESULTS

### The frequency of MB reports increases along with reports about sleepiness

Participants completed a sustained attention task (SART; see Methods) with embedded experience-sampling probes assessing attentional states and experienced vigilance (Figure 1). Following our previous analysis^7^, we aggregated Blank and No Recall reports, as well as reports about sleepiness (Sleepy and Very Sleepy) and vigilance (Alert and Very Alert). Frequencies of mental states, vigilance reports and mental states stratified by vigilance level are provided in Supplementary Table S1. For transparency, the pre-aggregation descriptive statistics for all vigilance reports and mental states are also provided (Supplementary Table S2).

**Figure 1.**
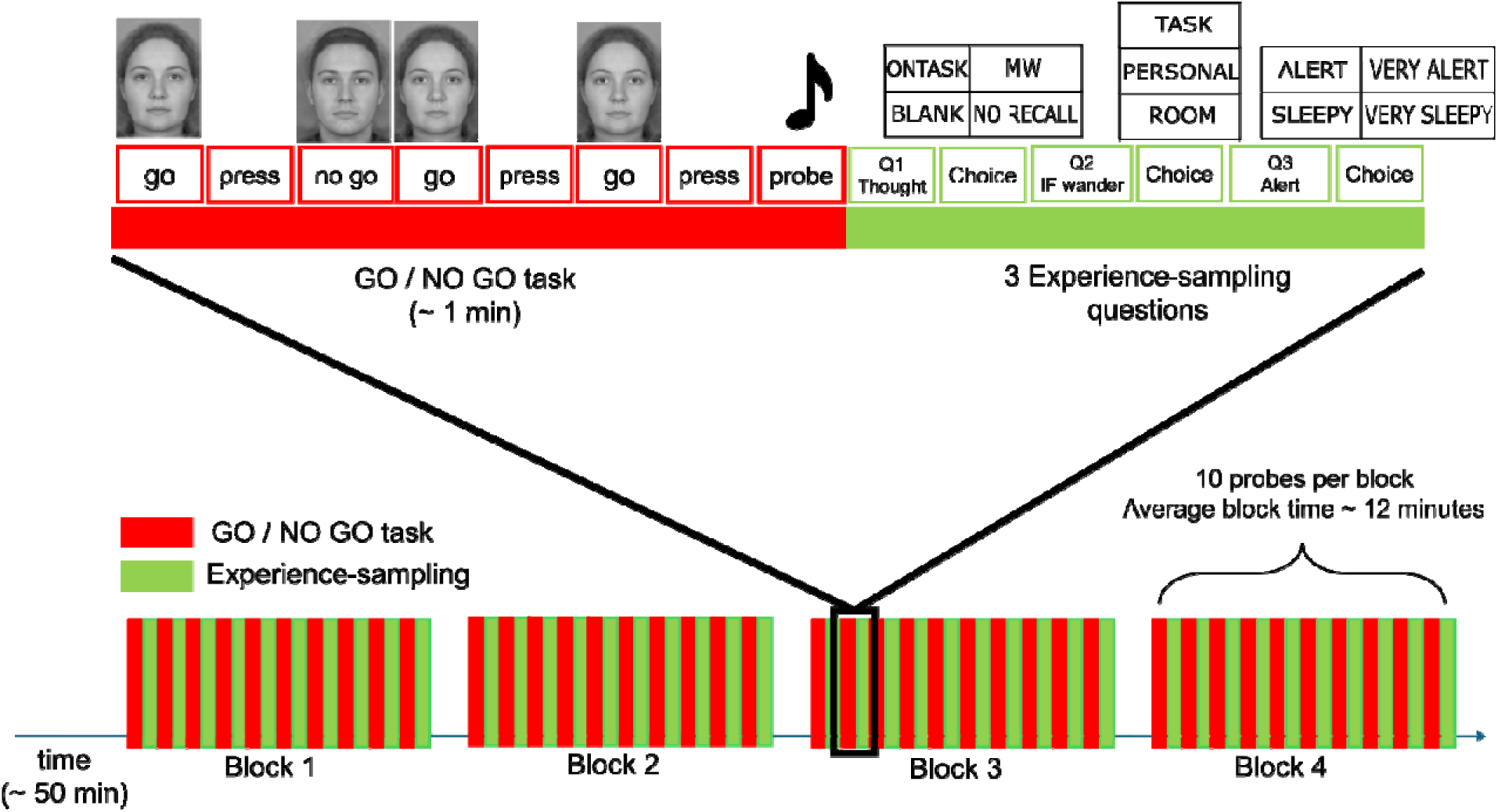
Experimental design of the sustained attention to report task (SART) combined with experience-sampling probes for reporting mental states and vigilance levels. Across four experimental blocks (∼12 min each), participants were presented with a sequence of female or male faces. Participants were instructed to press a button when a female face was presented (GO stimulus) and to avoid pressing a button when male faces were presented (NOGO stimulus). At random intervals, participants were presented with probes that invited them to report their mental content and their level of vigilance. First, participants indicated their thoughts by choosing amongst a) ONTASK thoughts, b) MW thoughts, c) BLANK, or d) I cannot recall my thoughts. If participants selected the MW thought option, they were further probed to report whether this related to the task, the experimental room or personal content. Finally, participants indicated their vigilance levels by choosing amongst a) Very Alert, b) Alert, c) Sleepy or d) Very Sleepy. In every block, 10 probes were presented, leading to a total of 40 probes per participant.

Overall, participants were engaged with the task (Mean proportions of reports ±SD: On-Task=.49±.25, MW=.39±.21, MB=.13±.14) and reported an alert state of vigilance (Mean proportions ±SD: Alert=.57±.3, Sleepy=.43±.3). Participants reported MB (Kruskal chi=43.81, df=2, p=3.1e-10, eta squared=.38, C.I.=[0.28,1]) and sleepiness (Kruskal chi=4.1, df=1, p=4.2e-2, eta squared=.04, C.I.=[0,1]) significantly less frequently (Figure 2a-b; Supplementary Table S3 for all pairwise contrasts across reports).

**Figure 2.**
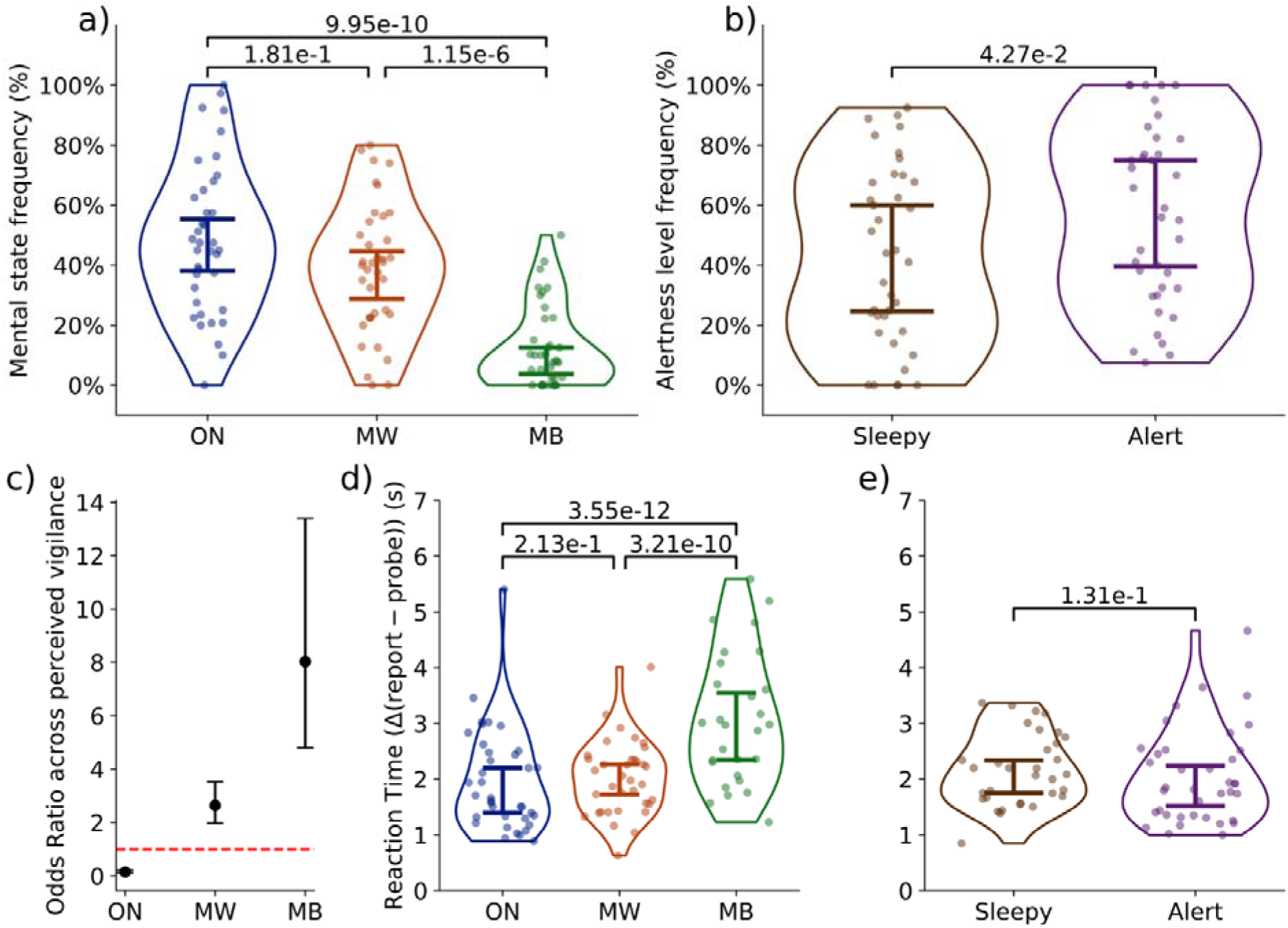
Characteristics of experience-sampling reports during the SART. a) Across the whole task, participants reported MB statistically less frequently compared to On-Task and MW. There was no significant difference between On-Task and MW reports. b) Across the whole task, participants tended to report being Alert more often than Sleepy. c) Perceived vigilance levels affected how frequently people experienced different mental states. When participants reported sleepiness, they were more likely to report MW and MB. When participants reported feeling alert, they were more likely to report On-Task thoughts. Black dots indicate logistic regression odds ratio beta estimates. Error bars indicate the confidence limits around the odds ratio estimates. The red, dotted line indicates equal probability between sleepiness and alertness (no effect). d) When probed to report their thoughts, MB was reported the slowest. e) There was no statistically significant difference in report times across vigilance levels. **Notes**: Violin plots indicate data dispersion. Point plots indicate single-subject mean estimates. Whisker plots indicate 95% confidence intervals around the median estimate. All contrasts were FDR-corrected for multiple comparisons. Mixed-effects models included subject ID as a random effects intercept. ON = On-Task thoughts, MW = mind wandering thoughts, MB = mind blanking.

To examine whether mental state reports were associated with vigilance, we fitted a binomial mixed-effects model for each mental state, with vigilance levels as the predictor. Our results showed that On-Task reports increased when participants reported being generally alert (very alert + alert), while MW and MB reports increased when participants reported being generally sleepy (very sleepy + sleepy; Figure 2c; Supplementary Table S4 for detailed pairwise contrasts).

Finally, we examined whether mental states and vigilance levels affected reaction times to probes (Chi squared=50.47, df=5, p=1.11e-09, BF10=1245.62). A contrast analysis of the main effects of mental states showed that MB tended to be reported slower compared to the other mental states. Yet, there were no statistically significant differences across vigilance levels. (Figure 2d-e; Supplementary Table S5 for descriptive statistics of reaction times across mental states and alertness levels; Supplementary Table S6 for all main and interaction pairwise contrasts).

A control analysis examining the potential influence of sex on thought proportions did not reveal any statistically significant effects across mental report categories (On-Task: t(19.81) = -1.02 [−0.29, 0.10], p = 3.20e-01, d = -0.38 [−1.04, 0.29]; MW: t(18.60) = 0.92 [−0.10, 0.25], p = 3.69e-01, d = 0.35 [−0.31, 1.01]; MB: t(30.99) = 0.44 [−0.07, 0.11], p = 6.62e-01, d = 0.14 [−0.52, 0.80]).

To determine whether MB reflects micro-sleep events, we quantified the mean duration of sequential “miss” errors preceding MB reports. If MB were preceded by multiple sequential misses, this would indicate prolonged disengagement and support the presence of micro-sleeps ^23^. Otherwise, it would suggest that MB reports occurred during wakefulness. We found that no MB report was preceded by sequential miss errors (more than 1 miss in a row) within the 10 s before the report, suggesting that the period of behavioural disengagement was very brief (<3 s). MB and sleepiness reports are associated with higher GS amplitude

We estimated the global signal amplitude as the real part of the Hilbert transform of the BOLD global signal. A gamma mixed-effects model using global signal amplitude as the dependent variable and containing both main effects (mental states, vigilance levels) and their interactions showed greater evidence compared to a null model of random intercepts (Chi squared=106.32, df=5, p=2.45e-21, BF10=1.85e13): Both MB reports and sleepiness were associated with increased global signal amplitude (Figure 3; Supplementary Table S7 for descriptive statistics of global signal amplitude across mental states and vigilance levels; Supplementary Table S8 for all main and interaction pairwise contrasts). When participants reported feeling alert, both MB and MW showed an increased global signal amplitude compared to On-Task thoughts; at the same time, during sleepiness, MB scans showed higher global signal amplitude only compared to MW thoughts (Figure 3). These results were replicated across various analysis lags (Supplementary Figure S2). When we regressed out task effects, the relationship between MB and global signal amplitude did not reach statistical significance (Supplementary Figure S3). For the sake of comparability with our past works ^9^, we also performed the analysis by regressing out the GS. We found that after GS regression, the relationship between sleepiness and global signal amplitude did not reach statistical significance, but MB was associated with significantly lower global signal amplitude, inverting the relationship (Supplementary Figure S4-5).

**Figure 3.**
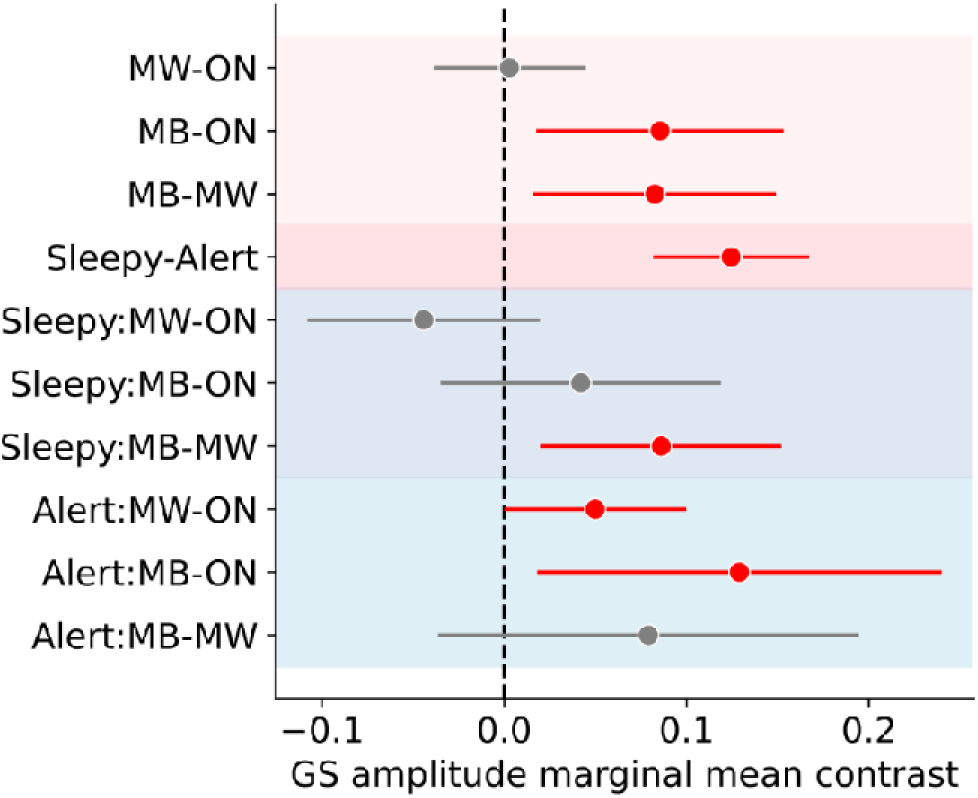
MB and sleepiness are accompanied by increased global signal (GS) amplitude. (main effects-red shades and interactions -blue shades). MB was characterised by higher GS amplitude relative to ON and MW (light red shade), and sleepiness was associated with increased GS amplitude compared to alertness (dark red shade). Simple-effect analyses indicated that during sleepiness, MB showed higher GS amplitude than MW (dark blue shade). In contrast, during alertness, both MB and MW were accompanied by increased GS amplitude compared to ON reports (light blue shade). **Notes**: Red points and error bars indicate statistically significant contrasts. Point plots and error bars indicate contrast means and 95% confidence limits. The x-axis represents marginal mean contrast estimates of global signal amplitude. All contrasts were FDR-corrected for multiple comparisons. Mixed-effects models included subject ID as a random effects intercept. All statistical models were compared with null models containing only random intercepts. ON = on-task thoughts, MW = mind wandering thoughts, MB = mind blanking. For the complete list of descriptive statistics and model parameters, see Supplementary Tables S7 and S8.

### Time-varying functional connectivity self-organises in distinct and replicable brain patterns

Our primary goal was to replicate the previously established relationship between fMRI connectivity patterns and MB, and to test whether the fMRI hyperconnectivity is linked to EEG SW-like activity during wakefulness. Therefore, we first asked whether time-varying FC during task engagement would organise in the same way as during rest. Establishing that these FC patterns reliably reappear during task performance was a necessary first step before testing their relationship to MB and SW-like activity.

We estimated how functional connectivity self-organises into distinct connectivity patterns during task engagement. Specifically, we estimated time-varying phase-based connectivity matrices using coherence as the connectivity measure. We then aggregated the matrices at every TR and every subject and applied K-means clustering (Manhattan distance) to generate representative patterns. The number of centroids varied from 3 to 8. Here, we focus on = 5, as it showed the largest interpattern covariance. Effectively, this suggests that at lower *k*, clusters were not that well defined, while at higher *k*, clusters contained redundant information (Supplementary Figure S6).

Overall, we found that functional connectivity during task decomposed into a) a pattern of complex inter-areal interactions, containing positive and negative phase coherence values, with higher coherence between the DMN and the limbic network (P1), b) a pattern of anticorrelations primarily between the visual network and the other networks (P2), c) a pattern of complex inter-areal interactions, containing positive and negative phase coherence values, with higher coherence between the DMN and the visual, control and limbic networks (P3), d) a pattern of primarily low inter-areal coherence, but between the visual and the somatosensory networks (P4), and e) a pattern of overall high inter-areal coherence (P5) (Figure 4a). Critically, patterns P1, P2 and P5 were reproduced irrespective of clustering algorithm (Euclidean or cosine distance) or denoising parameters (task regression, global signal regression), while P3 and P4 showed variations across clustering schemes (Supplementary Figures S6-9 for Manhattan distance; S10-13 for cosine distance; S14-17 for Euclidean distance).

**Figure 4.**
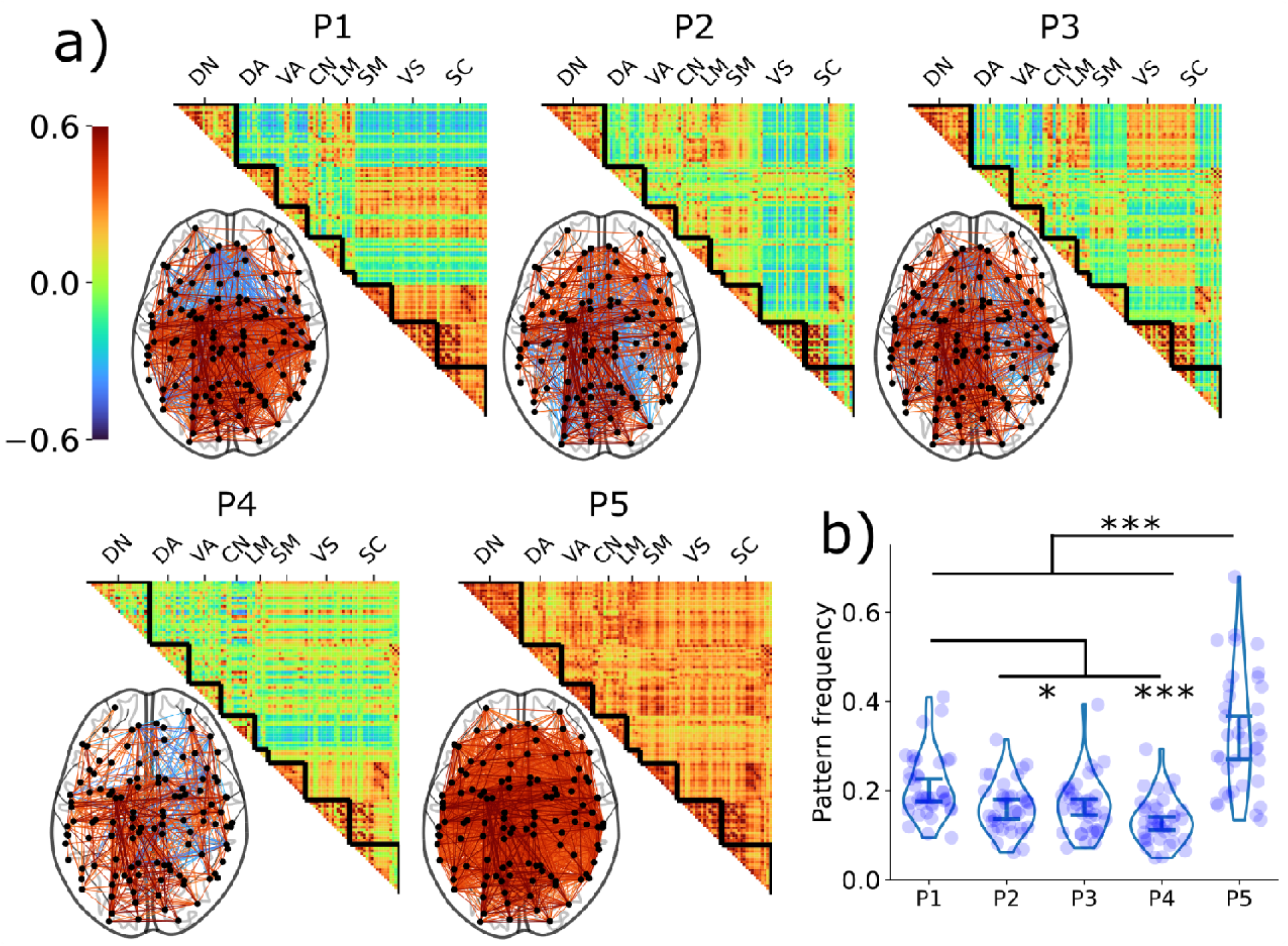
Time-varying functional connectivity during task engagement self-organises into five connectivity patterns. a) Data-driven clustering identified a pattern dominated by anticorrelations, with higher coherence between the DMN and the limbic network (P1), visual network anticorrelations pattern (P2), a pattern dominated by anticorrelations, with higher coherence between the DMN and the visual, control, limbic networks (P3), a lower overall coherence pattern (P4), and a cortex-wide high coherence pattern (P5). b) Patterns occurred at different frequencyrates, with P5 being the most frequent, followed by P1. The remaining patterns did not show a statistically significant difference in frequencies. **Notes**: Patterns are ordered based on decreasing standard deviation of the connectivity within a connectivity pattern. Colour bars and brain renders indicate the strength of coherence. Positive values indicate higher phase synchronisation. Values approaching zero indicate the absence of a systematic phase relationship. To facilitate visualisation onto brain renders, a 0.3 coherence edge threshold was applied. DN = default network, DA = dorsal attentional network, VA = ventral attentional network, CN = control network, LM = limbic network, SM = somatosensory network, VS = visual network, SC = subcortical network, *p<.05, **p<.01, ***p<.001.

The examination of pattern occurrence frequency suggested that P5 was the most frequent, followed by P1 (linear mixed effects model: Figure 4b; Supplementary Table S9 for descriptive statistics of connectivity pattern occurrence frequencies; Supplementary Table S10 for pairwise contrasts). Importantly, regressing out the global signal led P5 to be one of the least frequently occurring patterns (Supplementary Figure S8-9). This trend was also present in the clusters generated by cosine distance (Supplementary Figure S10-13). However, using Euclidean distance, a low-connectivity pattern tended to dominate the frequency space, instead of the high-connectivity one (Supplementary Figure S14-17). This potentially indicates algorithm-specific effects on the identified clusters. Finally, it should be noted that for Euclidean distance at *k* = 7, the algorithm did not assign any time points to one connectivity pattern when both global signal and task information were regressed out.

### Neurobehavioral coupling associates MB and vigilance with distinct brain configurations

Overall, mental states and vigilance levels were associated with different connectivity patterns. P1 had a higher proximity (lower distance values) to MB reports as well as reports about being alert (Figure 5; Manhattan distance from P1). Simple effects analysis further indicated that the link between P1 and MB was present only when participants reported being alert. Proximity to P2 differentiated both attentional lapses (MW thoughts and MB reports) from On-Task reports (Supplementary Figure S18); MW thoughts further differed from On-Task only during sleepiness, yet MB differed from both On-Task and MW thoughts during alertness. Proximity to P3 was associated with MB reports (Supplementary Figure S18), and this was consistent across vigilance levels. P4 correlated with sleepiness (Supplementary Figure S18), with no statistically significant relationship between mental states, either in terms of main effects or the simple effects.

**Figure 5.**
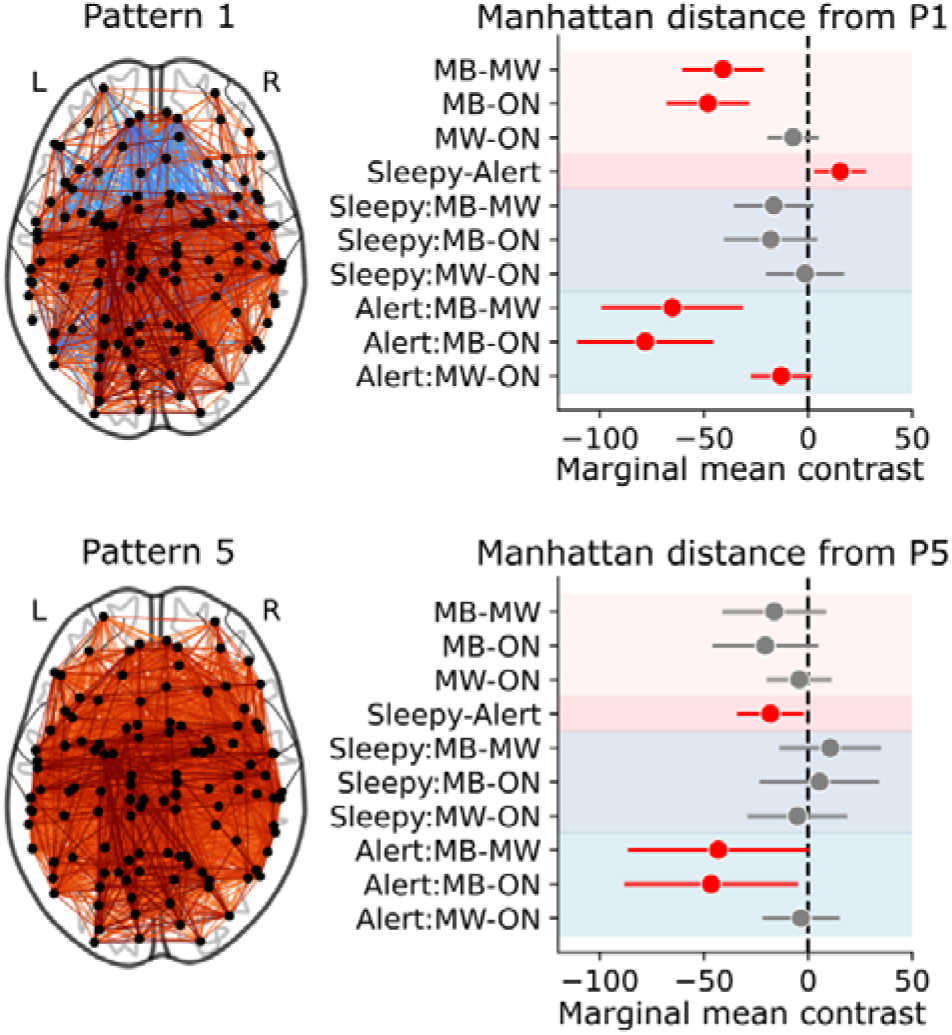
The link between brain patterns around thought reports and MB varies based on the levels of self-reported vigilance. (main effects-red shades and interactions -blue shades). P1 had higher proximity (lower distance values) to MB reports and to reports about being alert. Simple effects analysis further indicated that the relationship between P1 and MB was present only when participants reported being alert. Consistent with our hypothesis, proximity to P5 was higher during self-reports of sleepiness; importantly, when we stratified mental states across vigilance levels, we observed that connectomes around MB reports were closer to P5 during alertness compared to On-Task and MW thoughts, suggestive of the presence of two distinct modes of MB. **Notes**: Areas shaded in light red indicate the main effects of mental states. Areas shaded in dark red indicate the main effects of self-reported vigilance. Areas shaded in dark blue indicate simple effects when people reported being alert. Areas shaded in light blue indicate simple effects when people reported being sleepy. The x-axis represents marginal mean contrast estimates of distance to the accompanying pattern. Point plots in bold and error bars indicate contrast means and 95% confidence limits. Red points indicate statistical significance. All contrasts were FDR-corrected for multiple comparisons. Mixed-effects models included subject ID as a random effects intercept. All statistical models were compared with null models containing only random intercepts. ON = on-task thoughts, MW = mind wandering thoughts, MB = mind blanking. For the full list of descriptive statistics and model parameters, see Supplementary Tables S11 and S12.

Importantly, there was a higher proximity of the hyperconnected P5 to sleepiness reports (Figure 5; Manhattan distance from P5). However, no mental state showed a statistically significant relationship with P5. By stratifying mental states by vigilance levels, though, we showed that, during alertness, MB reports were indeed associated with higher proximity to P5, compared to On-Task and MW thoughts. No significant relationship was observed for self-reported sleepiness (See Supplementary Table S11 for descriptive statistics of the relationship between mental states, vigilance, and connectivity patterns, and Supplementary Table S12 for pairwise contrasts). Taken together, our initial hypothesis that the overconnected pattern would be linked to MB holds true, but only when participants report being alert.

To validate our analysis, we systematically varied the number of *k* clusters, GS regression, task regression, clustering algorithm and the lag of the BOLD signal (216 distinct permutations of analysis parameters) across all *k* clusters (Supplementary Figures S18-21 for Manhattan distance, *k* = 5; Dataverse: https://doi.org/10.58119/ULG/4LEK9M for all results dataframes across parameterisation). Overall, we demonstrate that our analysis is replicated across task regression and different BOLD lags. Regarding GS regression, we found that it abolished or inverted the relationship of vigilance reports across P1 and P5. Finally, different clustering algorithms produced similar main effects, with MB being related to the anticorrelations pattern (94/108 parameter permutations when excluding global signal regression), while sleepiness was associated with hyperconnectivity (106/108 parameter permutations when excluding global signal regression). Critically, the relationship between MB and hyperconnectivity during alertness was statistically significant only when using Manhattan distance as the clustering algorithm. For cosine and Euclidean distance, we note that MB’s proximity to P5 was consistently but non-significantly lower compared to on-task and MW thoughts (54/72 parameter permutations when excluding global signal regression and results from Manhattan distance).

### EEG SW-like activity and fMRI hyperconnectivity are linked to one another

Having validated that the hyperconnected pattern is related to decreased vigilance and MB reports during alertness, we investigated whether we could link the fMRI brain patterns and EEG SW-like activity, by means of Canonical Correlation Analysis (CCA; see Methods for details, Figure 6a,b).

**Figure 6.**
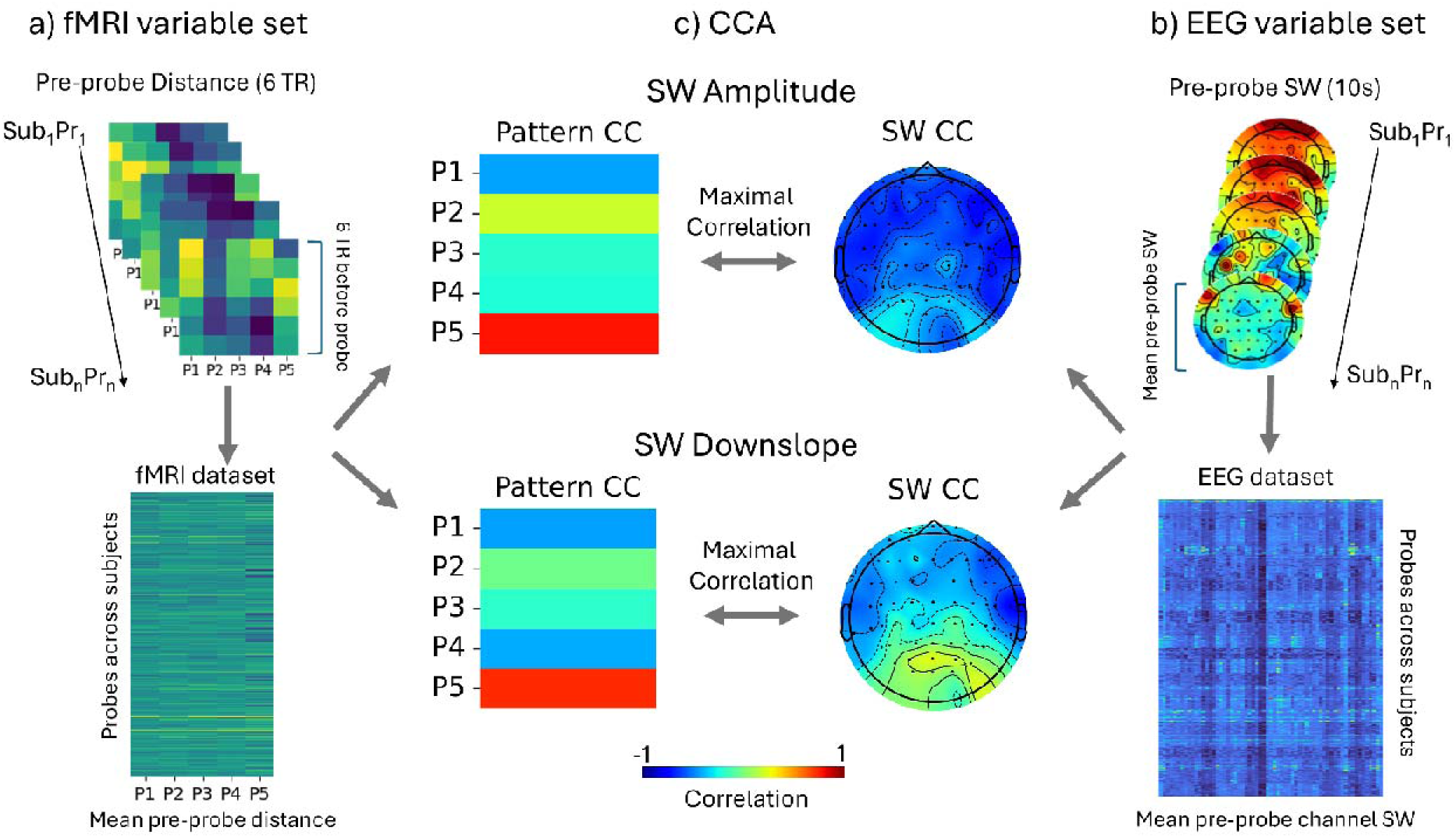
FMRI hyperconnectivity (P5) is linked to slow-wave-like (SW) activity. Canonical Correlation Analysis (CCA) is a multivariate statistical technique designed to identify latent relationships between two sets of multivariate variables. We here used CCA to uncover shared variance between time-varying fMRI connectivity patterns and EEG-derived slow-wave-like (SW) markers. **a)** For each probe, we first extracted the functional connectivity profile from the six TRs (∼10 sec) immediately preceding the probe. Then, we computed the Manhattan distance of functional connectivity at each TR relative to the five canonical brain connectivity patterns identified through clustering. These pre-probe distances were then averaged to yield a single distance measure per connectivity pattern, for each probe. By concatenating these measures across all subjects and probes, we constructed an fMRI dataset of size (N_subj_*N_probes_) x 5, where each row represents the pre-probe connectivity profile expressed in terms of similarity to the five patterns. **b)** In parallel, SW activity was quantified for the same pre-probe period. From the 10 sec of EEG preceding each probe, we extracted markers of SW activity. These features were likewise concatenated across all subjects and probes, yielding an EEG dataset of size (N_subj_*N_probes_) x 63. CCA was then applied to these paired datasets to identify canonical components (linear combinations of EEG and fMRI variables) that maximally correlate across the two modalities**. c)** SW Amplitude: The distance to brain patterns P1,3,4 showed a negative correlation with the first fMRI canonical component, while the distance to brain pattern P5 showed a positive correlation to the component, indicating that fMRI hyperconnectivity is linked to EEG slow-wave like activity. SW Downslope: The first canonical component was correlated with time-varying functional connectivity that was more similar to brain pattern P5 and less similar to P1, 3, and 4. The SW downslope component further showed a fronto-centro-temporal orientation, suggesting that a stronger SW downslope is linked to fMRI hyperconnectivity. **Notes:** Heatmaps illustrate the correlation between the fMRI canonical components and the Manhattan distance of the connectivity matrices from brain patterns (P1-5). EEG topomaps depict the correlation between the EEG canonical components and SW activity in each brain channel. Colour bars represent correlation values ([-1,1]) for both the fMRI and the EEG canonical components. Statistical significance of each canonical correlation was evaluated by estimating an empirical null distribution of canonical correlations using permutations of one of the datasets. CCA = canonical correlation analysis, CC = canonical component, Corr = correlation, SW = slow-wave-like.

Overall, we observed that proximity to the hyperconnected pattern correlated with the amplitude and the downslope of the SWs in a subset of 25 participants containing complete EEG and fMRI sessions. For the SW amplitude, the fMRI component positively correlated with distance from hyperconnectivity, while the EEG component negatively correlated with SW amplitude. These results suggest that, as SW amplitude decreases, functional connectivity moves further away from the high-coherence pattern (P5) (Figure 6c; SW Amplitude). Regarding the SW downslope, a frontal-temporal-central SW component showed anticorrelation with proximity to P5. Similarly to SW amplitude, as the downslope of the SW increased, functional connectivity organised in a manner more similar to the fMRI hyperconnectivity (Figure 6c; SW Downslope). No statistically significant relationship was observed between SW density and upslope and the brain patterns.

A replication of our analysis by varying task regression, GS regression, and clustering algorithm showed that frontal-central-temporal SW amplitude, upslope and downslope were consistently related to the hyperconnectivity brain pattern. Visual inspection showed similar canonical components across clustering algorithms (Supplementary Figures S22-25 for Manhattan distance; Dataverse: https://doi.org/10.58119/ULG/4LEK9M for all canonical correlations across parametrisation).

### Control analysis of differences between Blank and No Recall reports

In order to assess whether Blank and No Recall reports are associated with different neural effects, we redid the analyses by dividing the MB category further into these two categories. It should be noted that in the present dataset, only 17 participants reported at least one “No Recall”, and only 8 had more than one such report, hence limiting the inferential capacity for this response option. Regardless, we replicated the association between higher GS and Blank responses. In contrast, the relationship between No Recall and GS did not reach statistical significance (Supplementary Table S13). Similarly, regarding the neurobehavioral coupling, all previously observed relationships between Blank reports and connectivity patterns were replicated, except for the interaction between MB and alertness during hyperconnectivity. In contrast, results for No Recall responses did not reach statistical significance (Supplementary Table S14).

## DISCUSSION

We combined simultaneous fMRI-EEG during experience sampling to uncover nuanced modes of MB reports. We specifically sought to understand how MB relates to levels of perceived vigilance and how multi-modal imaging helps unravel associated neural profiles. The underlying hypothesis was that MB’s overall positive fMRI functional connectivity would be mediated by EEG slow-wave (SW) like activity, hence shedding light on the role of reduced arousal in the reportability of seemingly content-less experiences.

Behaviourally, we showed a link between MB reports and experienced low arousal. First, MB was reported less frequently compared to mental states with thought content (i.e., On-Task, MW), in accordance with past findings about the sparse prevalence of MB reports ^7,9^. At the same time, MB report frequency increased as participants became sleepy. These results replicate previous work showing that lower vigilance levels accompany MB ^7,14^ and align with recent findings showing more frequent MB reports at lower arousal states, such as after sleep deprivation ^10^. The additional finding that MB had the slowest report time further corroborates the reduced vigilance scope ^24,25^. It should be noted that we here operationalised vigilance levels (sleepiness and alertness) based on participants’ self-reports to experience-sampling probes. Self-rated vigilance has shown a consistent relationship to MB, where lower levels of experienced alertness predict more frequent MB reports ^7,12,14,26^. Physiologically, experienced sleepiness has been systematically linked to lower cortical and autonomic arousal ^27,28^, thereby mapping the experience of sleepiness to discrete cortical correlates.

At the brain level, we replicated that MB is differentiated from content-oriented mental states based on a higher amplitude of the fMRI GS ^9^. At the same time, we show, for the first time, that GS amplitude increased as participants reported lower levels of experienced vigilance. Although the interpretation of the GS and what it represents remains an open question ^29^, it is generally assumed that the variance it introduces can be accounted for by artifacts, such as motion and physiology, such as heart rate and respiration ^30,31^. However, GS was also shown to covary with electrophysiological^32^ and metabolic markers ^33^ of ongoing neural activity, implying that it reflects neurophysiological activity, as well. Pertinently to this study, arousal fluctuations during wakefulness ^18,30,34^ and sleep ^30,35^ were shown to modulate the amplitude of the GS, leading to the formulation that arousal propagation across the cortex elicits cortical synchronisation and subsequent GS amplitude increases. Our results build upon this literature and show that fluctuations in the GS amplitude not only track physiological arousal and sleepiness but could also carry information about experiential mental states.

Having shown that the amplitude of the GS is linked to MB and subjective levels of sleepiness, we then set out to examine how ongoing mental states and vigilance reports related to whole-brain activity. As thinking is inherently a dynamic process, an approach that describes how thoughts evolve might be better situated when considering their underlying brain dynamics ^1^. The biological and theoretical motivation behind such a dynamics approach is that functional connectivity varies over time, organising itself across consistent and robust connectivity patterns that show generalizability across species ^36–38^, suggesting that they represent a fundamental mode of cortical organisation ^9,39,40^. Pertinently to ongoing thinking, the repertoire of dynamic brain states appears to vary with consciousness level ^39,40^ and task engagement ^41^, providing a time-resolved approach of mapping brain states to subjective phenomenology ^9,42^. Following this rationale, we examined how time-varying functional connectivity organises itself during task engagement. We found that functional connectivity can be optimally summarised into five patterns, ranging from complex inter-areal anticorrelations to globally synchronised modes. The patterns elicited during the task bear resemblance to those previously reported during rest ^9,43^ and at different levels of consciousness ^40^, suggesting that across various conscious states and cognitive demands, the brain explores similar topologies.

We consistently replicated three previously observed patterns ^9,40–43^, namely the complex anticorrelations pattern (P1), the pattern driven by the visual network (P2), and the pattern with an overall high positive coherence (P5). The overall positive coherence pattern was further found to be the most frequent, followed by the complex anticorrelations pattern. A significant deviation from previous works is that the low-coherence pattern, marked by reduced functional connectivity, was here clearly defined only at higher *k* clusters (>6). During rest, such low-coherence pattern appears to dominate the clustering space ^40^. We believe these results can be accounted for by the fact that we directly estimated connectivity patterns from task time-varying FC. Indeed, while rest and task connectivity produce similar whole-brain network structure ^44^, the functional properties of rest and task connectivity differ, with task producing weaker connectivity and less variable neural dynamics ^45,46^. Despite similar network organisation, task connectivity, compared to rest, also shows altered within-network ^47^ and between-network interactions ^48,49^, and increased connectivity across task-relevant cortical sites ^48,49^. Overall, our results support the idea that, while the brain explores similar connectivity repertoires during rest and task engagement, the dynamics of this exploration are determined by the task demands.

To relate the brain patterns with the ongoing vigilance and mental states, we estimated the distance of all connectivity matrices to each of the brain patterns produced, as in our previous work ^9^. Overall, different patterns captured nuanced interactions with mental states. Analysing the complex anticorrelations patterns P1 and P3, we found that they bore a greater relationship to how the connectome around MB reports was organised. These results were contrary to our original hypothesis, i.e. that MB would be linked to fMRI hyperconnectivity. Similar anti-correlated patterns were previously associated with enhanced stimulus perception at threshold levels ^43^. We recently proposed that MB might represent moments of transition (successful or unsuccessful) between different contents. In other words, when probing during transitions, the absence of a clearly formulated content phenomenologically could translate into “no thought”, leading to MB reports ^5^. As such, a potential interpretation is that the predominantly anti-correlated brain pattern might reflect moments where reportable content is not yet established. This interpretation is supported by the fact that spontaneous thought initiation is accompanied by increased connectivity between the default mode network, the limbic and the control network, regions with positive correlations in both P1 and P3 ^50–52^. However, since the connectivity patterns in the study here were generated from task-BOLD measurements, pre-probe periods inherently contain a mixture of overlapping cognitive processes associated with the task, such as perceptual discrimination, inhibition and motor processing. Therefore, this interpretation about thought transitions remains to be further explored.

Following our main line of inquiry, we set out to examine how the hyperconnected P5 relates to different electrophysiological and experiential events. Although the main effects did not provide evidence for a relationship between MB and the hyperconnected pattern, an interesting interaction emerged when vigilance levels were taken into account: by comparison to other mental states, MB-related connectivity was more similar to P5 when participants reported being alert. This finding offers partial support for our hypothesis that MB is linked to the hyperconnected brain pattern as previously shown during a task-free fMRI investigation of MB ^9^. To gain some insights about this interaction, we can turn to our recently proposed physio-cognitive model of thought reportability, which hypothesises that mental content results from the interplay between cognitive mechanisms and physiological arousal ^4^. In that model, arousal changes shape cortical activity patterns, which in turn influence how MB can be reported. In other words, we hypothesised that arousal functions as the regulator of a neural background ^53,54^. By modulating the strength of dynamic neural assemblies, it alters our capacity to introspect and report our thoughts ^4,5^. This account permits different types of MB to emerge, associated with discrete functions and neural correlates. Indeed, the idea of a unified blank has been criticised ^8,55^. As proximity to patterns is not necessarily mutually exclusive, every connectome retains a modicum of similarity to every identified connectivity pattern. Therefore, a potential reconciliatory approach would be that, across vigilance levels, MB approaches P1, as it potentially reflects cortical mechanisms associated with thought transition. However, during alertness, this might also be accompanied by overall changes to cortical arousal (as indicated by the proximity to P5).

When looking into the properties of the hyperconnected P5, we identified a correlation with increased amplitude in the EEG in the form of SWs. Particularly, SW markers during wakefulness (amplitude, downslope) correlated with functional connectomes more proximal to the hyperconnected P5 and were more distant to the anticorrelated patterns P1-4. The notion about the link between the fMRI hyperconnectivity and sleep is consistent with previous works ^15,56^. We recently hypothesised that such globally high synchronisation may also result from the presence of SW-like activity during wakefulness ^4,9^. Indeed, SW activity originates from alternating patterns of neural firing (up states) and neural silencing (down states) at a slow scale (1-4Hz) in both sleep ^57^ and wakefulness ^58^. While SWs represent localised cortical events, they are not static and propagate across the cortex as a travelling wave ^59,60^. Subsequently, propagation may elicit long and short-range neural synchrony ^61^, taking the form of cortex-wide increases in functional connectivity ^16,62^. Importantly, GS regression abolished or even inverted the relationship between the hyperconnected pattern and sleepiness. We previously showed that GS regression abolished hyperconnectivity in the clustered patterns and removed their subsequent relationship with MB ^9^. Therefore, we had hypothesised that GS is a core component of hyperconnectivity, though it remains ambiguous whether they reflect underlying neural event viewed through different analytical lenses (i.e., global BOLD fluctuations versus network-wide functional connectivity). However, because the GS tightly covaries with cortical and autonomic arousal ^30^, and given that GS regression abolishes the P5 state, we propose that P5 represents a global background state of altered arousal, rather than a neural substrate uniquely related to the phenomenology of MB. In turn, we consider that this altered arousal profile impairs general processes associated with wakefulness, such as maintaining mental content and metacognitive monitoring of content, which would explain why MB is more prevalent during lower experienced vigilance.

The most prominent finding of our analysis concerns the interaction between experienced vigilance and MB reports, which we suggest points to distinct cortical profiles underlying the common reportability of thought-free experiences. So far, MB reports have typically been interpreted as indicating a broadly content-free phenomenology that arises spontaneously during rest or task engagement. This interpretation assumes that instances in which the mind is described as “empty” are neurocognitively similar, since they share comparable phenomenological reports. We recently argued, though, that this might not be the case ^4^. Other content-free or content-reduced states, such as meditation and white dreams, can differ in their underlying mechanisms and phenomenology, including their levels of arousal, attention, perceptual richness, and temporal awareness. Here, we provide empirical evidence that the phenomenology of MB is supported by distinct neural profiles. Although protocols directly targeting the cognitive and microphenomenological constituents of MB are expected to determine such nuances, our results emphasise that shared reportable phenomenology does not necessarily translate to a shared neurocognitive state, and in fact, ostensibly similar reports can be supported by different cortical profiles based on levels of alertness.

Overall, our results have various implications. At the theoretical level, theories of consciousness that aim to explain the neural mechanisms underlying conscious content face the challenge of accounting for the nuanced aspects of MB, which, to our knowledge, have not yet been adequately incorporated. For example, the global neuronal workspace theory (GNWT ^63^) posits that conscious content manifests as the result of broadcasting specific contents to a global workspace consisting of specialised cognitive systems (such as attention, working memory, language, decision-making) that support the manipulation of the content and its reportability ^64^. Accordingly, ignition, the widespread and self-sustaining activation of distributed brain networks, occurs at content onset and offset, demarcating the presence of conscious content ^65^. In this sense, MB could be more likely to occur during moments between two ignition events and/or following a failed ignition. In these moments, the hyperconnected brain profile, marked by the absence of functional segregation, could impede the broadcast of specific content to the global workspace. On the other hand, information integration theory (IIT) starts by identifying properties assumed to characterise all conscious experiences, including MB, and postulates physiological mechanisms that can possibly actualise these axioms. According to IIT, a system is conscious if it can exert cause-and-effect power itself, with its level determined by the amount of irreducible integrated information it supports ^66^. From IIT’s perspective, global integration does not necessarily imply a high level of consciousness, as it can accompany loss of differentiation, leading to reduced integrated information and collapsed cause-effect structures, hypothesised to correlate with those found in generalised epilepsy ^67^. While there are many other theories of consciousness ^68,69^, an explanation of MB, as an intermediate, seemingly content-less state between wakefulness and low arousal, can be challenging to some if not all theories. As such, establishing MB as an explanandum for theories of consciousness would provide a meaningful test for such theories and contribute to the advancement of the field.

At the clinical level, the phenomenal profile of MB can assist the characterisation of specific clinical pathologies, such as ADHD ^70–73^, depression ^74,75^ and anxiety ^76^, where an increase in the frequency of empty thoughts is observed. Reports of “absence of thoughts”, “inability to recall immediate past events” and “gaps in experience” indeed do manifest across various clinical conditions ^4^. In this sense, the degree of content degradation or absence may serve as a meaningful experiential marker for characterising the symptomatology of different clinical conditions. In turn, investigating this phenomenology may guide the identification of the underlying neurobiological mechanisms. For example, the presence of SW activity, here shown to directly relate to the hyperconnected pattern typically associated with MB, can account for ADHD symptoms ^72^. Prospectively, we can leverage this experience of “absence of thoughts” to improve the diagnostic capacity of screening and neuropsychological testing by incorporating phenomenal reports.

The close association between MB and both subjective sleepiness and sleep-like brain states raises the question of how MB relates to microsleeps. Microsleeps are typically defined as brief (3 to 15 s) episodes characterised by slow eyelid closure and head nodding, accompanied by a global shift in neural activity from wakefulness toward NREM sleep and a failure to respond to the task at hand ^77,78^. Microsleeps and MB share commonalities, such as increased report frequency after sleep deprivation, or being marked by increased power in lower EEG frequencies. Yet, they are distinguished in at least two ways. First, the EEG neural correlates of MB and microsleep are distinct since MB has been characterised by the presence of local SW, while the shift in brain activity associated with microsleeps is defined at the scalp level ^19^. Second, when these are offered as separate categories during experience sampling, participants consistently distinguish periods of sleep from MB ^10,79^. In our data, we observed that the unresponsiveness associated with MB, in the form of miss errors, was very brief and always limited to isolated trials. As this corresponds to a behavioural disengagement of 1.5 seconds, it is likely that MB reports genuinely reflected blank instances during wakefulness rather than micro sleeps.

Our study is not without limitations. To account for the rarity of “No Recall” reports and in line with previous work ^7,70^, we aggregated blank reports (Blank and No Recall), as they both point to the phenomenology of “no immediate past thought content”. However, it should be noted that these two categories can reflect different cognitive mechanisms (e.g., metacognitive awareness of empty moments vs memory failure) ^4^. Therefore, in order to assess whether Blank and No Recall reports are also associated with different neural effects, we further split the MB category into Blank and No Recall reports and repeated the analysis. We found that, while we replicate the findings for the Blank category, no results from the No Recall reports reached significance, likely due to the very low number of these reports across a few participants. Another limitation is that functional connectivity cluster estimates rely on a series of methodological choices, including preprocessing parameters, the selection of connectivity metrics, and the details of algorithmic implementation. To address the combinatorial issue of our different decision-making processes, we opted for a systematic characterisation of our clustering approach across global signal regression, task regression, clustering algorithm, analysis lag, and clustering k. While this aims to ameliorate the issue of multiple potential pipelines, we noted some discrepancies in pattern frequency across the clustering metric. When Euclidean distance was used, the low-connectivity pattern emerged as the most frequent one. Therefore, our analysis could also capture effects elicited by clustering parameters, rather than properties of time-varying cortical organisation. Given the low number of subjects, in the CCA analysis, we did not group probes by subject, following previous MB explorations ^10^. While this approach suggests a stable relationship in our data, further work is required to demonstrate the generalizability of this finding. A final issue concerns the generalizability of our analysis to clustering distance. We based our primary analysis on the most established clustering metric for fMRI connectivity, the Manhattan distance, yet examined its invariance under Euclidean and cosine distances. Our results were significant only when using the Manhattan distance. This can potentially be attributed to how the distance is estimated. Manhattan distance sums absolute differences across dimensions, therefore making it more sensitive to distributed, coordinate-wise changes. However, the squared term in Euclidean distance disproportionately weights large, noisy fluctuations and downplays weaker connections. Finally, cosine distance removes overall magnitude and looks only at the relative pattern of connections, and, therefore, may not capture differences that depend on the overall strength of the connections. Here, all distances generated a hyperconnected pattern. However, only Manhattan-derived hyperconnectivity was significantly linked to MB, with the remaining distance metrics showing non-significant, yet similar results in directionality. Together, these findings suggest that our results may not fully generalise across clustering metrics, and that the association with MB may be most robustly captured by distance measures sensitive to distributed, small-magnitude changes across connections.

In conclusion, we show that the previously reported hyperconnected brain pattern and sleep-like intrusions that correlate with MB reports might represent two sides of the same coin. We show that, while MB and sleepiness both relate to fMRI hyperconnectivity, their interaction uncovers dissociable cortical correlates of MB based on how participants evaluate their vigilance level. Our work builds on the quest to bridge mental content and its absence with measurable brain activity and provides insights into how ongoing thinking is maintained during wakefulness. Our findings elaborate on how experienced and cortical arousal lead to subjective mental content, providing a more refined characterisation of the neural correlates of seemingly “thought-less” mental states.

## METHODS

### Participants

Thirty-nine participants were recruited for the study (mean age = 26.4, sd = 4.5, range = [18-32]; females = 24). All participants reported normal or corrected-to-normal vision and no neurological or psychiatric disorders. One participant was excluded due to subsequent changes in the experimental paradigm, resulting in a total sample size of n = 38. Participants received monetary compensation. The project was approved by the Monash University Human Resources Ethics Committee (Project ID 17674).

### Experimental Design and Stimuli

During a SART ^80^, participants were asked to pay attention to a series of pictures of human faces. In every trial, face stimuli (one male face, or eight different female faces, extracted from the Radboud Face Database ^81^) were presented for 500 ms, followed by a blank screen. The trial duration was between 1000–1500 ms (random uniform jitter). Participants were instructed to indicate the presence of a female face (GO trials) by pressing a button on a keypad placed under their dominant hand and to withhold button presses (NOGO trials) for the male face. The order of the face stimuli was randomised before the onset of every session. Participants completed four blocks of the task, with self-paced breaks in between, each block lasting approximately 12-15 minutes and totalling approximately 50 minutes.

Intermittently during the SART, we utilised experience sampling to probe reports of participants’ mental states and vigilance levels throughout the session. The mental state options were explained to participants both orally and in writing before the experience-sampling session. Specifically, every 30-70s (uniformly jittered), we interrupted the SART and displayed the word “STOP” on the screen. Following the probe, participants were asked: “*Just before the interruption*, *where was your attention focused?”* Possible responses were a) On-Task (defined as entirely focused on the task), b) Off-Task (defined as thinking about something other than the task), Blank (defined as not thinking of anything), and d) I cannot recall. If participants selected the Off-Task option, they were additionally presented with the following question: *“Just before the interruption, what distracted your attention from the task?”* Available options were: b1) Something in the room, b2) Something personal, and b3) Something about the task. In the present work, we operationally defined MB as any Blank or No Recall report. Apart from gaining statistical power, this definition allowed us to include different moments when thought content was absent, in line with previous examinations of MB ^7,70^. After reporting their mental states, participants were asked: *“Over the past few trials: How alert have you been?”* Available levels were: a) Extremely alert, b) Alert, c) Sleepy, and d) Extremely Sleepy. Each block included 10 sets of probes. Each participant responded to 40 probes across four blocks.

### Behavioural Analysis

Reaction times were defined as the difference between the onset of the probe screen and the button press, and they were contrasted across mental states and vigilance levels. Reaction times <.25s or >8s were discarded following our previous analysis protocol ^10^. Overall, we removed ∼8% of the total probes (118/1515).

### EEG

#### EEG Acquisition Parameters

EEG data were acquired using the BrainAmp MR plus system (Brain Products, Munich, Germany), equipped with a 64-channel EEG Cap (64Ch Standard BrainCap-MR with multitrodes). Electrode placement followed the international 10-20 system. The ground electrode was positioned in the location of AFz, and the reference was in the location of FCz. The electrodes were prepared with conductive gel to reduce impedance levels to <5kΩ. The data were recorded with the BrainAmp MR plus DC and bipolar BrainAmp ExG MR amplifiers and the BrainVision Recorder (Version 2.2) software with a resolution of 0.5μV/bit at 5000 Hz and with a hardware bandpass filter (0.01-250Hz). The EEG and fMRI data acquisition were synchronised using the Brain Products SyncBox.

#### EEG Preprocessing

Gradient artifacts were removed via subtraction of a mean template in each channel, according to the average artifact subtraction (AAS) methodology, utilising the software BrainVision Analyzer 2 (Version 2.2, Brain Products GmbH, Gilching, Germany). A sliding moving-average window implementation of 21 epochs of a full gradient artifact (1.56-sec time window) was used for the construction of the average artifact template. The reason for using 21 epochs was based upon the default settings of the BrainVision Analyzer. After this step, the data were downsampled to 500Hz. Cardioballistic artifacts were also removed using the AAS method: R-peaks were identified via a semi-automated pipeline, then they were visually inspected and corrected when necessary. Further preprocessing was done using the EEGLAB MATLAB toolbox (v2021.1). Data were split into 25s pre-probe epochs and filtered using a 100Hz high-cut-off Hamming filter. Next, the data were visually inspected for bad electrodes and epochs. Eye-and cardiac-related artifacts were detected and removed using ICA. At that stage, as we noticed that residual cardioballistic artifacts were still present in the frontal and temporal channels, we applied denoising source separation (DSS) on heartbeat-evoked potentials and removed the time-locked activity from every channel ^82^.

#### SW Extraction

We followed a previously validated method to detect SW activity ^7^. First, data were re-referenced to the average of mastoid electrodes and band-pass filtered within a [1,10] Hz range with a Chebyshev filter. Then, slow waves were defined in each electrode separately by identifying two consecutive zero-crossings (start and end of the slow waves). For each wave, we extracted its peak-to-peak amplitude, that is, the amplitude between the positive and negative peaks of the slow wave. Waves with extreme amplitudes (> 75µV positive peak, and > 150µV peak-to-peak di erences) and with duration corresponding to larger than 7Hz activity were excluded. Finally, we selected the top 10% of the largest waves in each electrode and extracted their timings, as well as their downward and upward slopes. Slow wave markers were eventually extracted from a 10s window before each probe.

### fMRI

#### fMRI Acquisition Parameters

Participants were instructed to lie still in the MRI scanner. Data were collected using a Siemens Skyra 3T MRI system, equipped with a Siemens 20-channel Head/Neck coil, allowing for simultaneous EEG and ECG recordings. The participants’ heads were stabilised with foam pillows placed between the cap and the MR coil. Anatomical images were acquired using a T1-weighted MPRAGE sequence with a voxel size of 1.00 mm³, TR=2.3s, TE=2.07s, and a slice thickness of 1.00 mm. Functional images were obtained using an Echo Planar Imaging (EPI) sequence with the following parameters: TR=1560 ms, TE=30ms, voxel size=3 mm isotropic; no interslice gap, FoV=192 mm, slices=42, matrix size: 94 x 94, slice orientation: T > C-7.7 > S0.9, phase encoding direction: A >> P, in-plane Acceleration (iPAT): 2, slice acceleration (multiband): 2, flip angle=70°.

#### fMRI Preprocessing

Before preprocessing, fMRI data were converted to BIDS ^83^. Preprocessing was performed using fMRIPrep 23.2.1 (Esteban et al., ^84^, RRID:SCR 016216), based on Nipype 1.8.6 (Gorgolewski et al. ^85,86^, RRID:SCR 002502 and the AFNI software 24.3.00 ^87^. We are reporting the preprocessing output text of the fMRIPrep software as provided.

#### Anatomical data preprocessing

A total of 1 T1-weighted (T1w) images were found within the input BIDS dataset. The T1w image was corrected for intensity non-uniformity (INU) with N4BiasFieldCorrection ^88^, distributed with ANTs 2.5.0 0 (Avants et al. ^89^, RRID:SCR 004757), and used as T1w-reference throughout the workflow. The T1w-reference was then skull-stripped with the antsBrainExtraction.sh workflow (from ANTs), using OASIS30ANTs as target template. Brain tissue segmentation of cerebrospinal fluid (CSF), white-matter (WM) and gray-matter (GM) was performed on the brain-extracted T1w using fast (FSL (6.0.7.14), RRID:SCR 002823, Zhang et al., ^90^). Volume-based spatial normalisation to one standard space (MNI152NLin2009cAsym) was performed through nonlinear registration with antsRegistration (ANTs 2.5.0), using brain-extracted versions of both T1w reference and the T1w template. The following template was selected for spatial normalisation and accessed with TemplateFlow (23.1.0, Ciric et al. ^91^ : ICBM 152 Nonlinear Asymmetrical template version 2009c [Fonov et al. ^92^, RRID:SCR 008796; TemplateFlowID: NI152NLin2009cAsym].

#### Functional data preprocessing

For each of the 4 BOLD runs per participant (across all tasks and sessions), the following preprocessing was performed. First, a reference volume was generated, using a custom methodology of fMRIPrep, for use in head motion correction. Head-motion parameters with respect to the BOLD reference (transformation matrices, and six corresponding rotation and translation parameters) are estimated before any spatiotemporal filtering using mcflirt (FSL, Jenkinson et al., ^93^). The BOLD reference was then co-registered to the T1w reference using mri coreg (FreeSurfer) followed by flirt (FSL, ^93^) with the boundary-based registration ^94^ cost-function. Co-registration was done using conFig.d with six degrees of freedom. Several confounding time series were calculated based on the preprocessed BOLD: framewise displacement (FD), DVARS and three region-wise global signals. FD was computed using two formulations following Power (absolute sum of relative motions, Power et al., ^95^) and Jenkinson (relative root mean square displacement between affines, Jenkinson et al., ^93^). FD and DVARS are calculated for each functional run, both using their implementations in Nipype (following the definitions by Power et al., ^95^). The three global signals are extracted within the CSF, the WM, and the whole-brain masks. Additionally, a set of physiological regressors were extracted to allow for component-based noise correction (CompCor, Behzadi et al., ^96^). Principal components are estimated after high-pass filtering the preprocessed BOLD time series (using a discrete cosine filter with 128s cut-off) for the two CompCor variants: temporal (tCompCor) and anatomical (aCompCor). tCompCor components are then calculated from the top 2% variable voxels within the brain mask. For aCompCor, three probabilistic masks (CSF, WM and combined CSF+WM) are generated in anatomical space. The implementation differs from that of Behzadi et al. in that instead of eroding the masks by 2 pixels on BOLD space, a mask of pixels that likely contain a volume fraction of GM is subtracted from the aCompCor masks. This mask is obtained by thresholding the corresponding partial volume map at 0.05, and it ensures components are not extracted from voxels containing a minimal fraction of GM. Finally, these masks are resampled into BOLD space and binarised by thresholding at 0.99 (as in the original implementation). Components are also calculated separately within the WM and CSF masks. For each CompCor decomposition, the k components with the largest singular values are retained, such that the retained components’ time series are sufficient to explain 50 percent of variance across the nuisance mask (CSF, WM, combined, or temporal). The remaining components are dropped from consideration. The head-motion estimates calculated in the correction step were also placed within the corresponding confounds file. The confound time series derived from head motion estimates and global signals were expanded with the inclusion of temporal derivatives and quadratic terms for each ^97^. Frames that exceeded a threshold of 0.5 mm FD or 1.5 standardised DVARS were annotated as motion outliers. Additional nuisance time series are calculated by means of principal components analysis of the signal found within a thin band (crown) of voxels around the edge of the brain, as proposed by Patriat et al., ^98^. All resamplings can be performed with a single interpolation step by composing all the pertinent transformations (i.e. head-motion transform matrices, susceptibility distortion correction when available, and coregistrations to anatomical and output spaces). Gridded (volumetric) resamplings were performed using nitransforms, conFig.d with cubic B-spline interpolation.

Apart from the fMRIPrep pipeline, we additionally applied a Gaussian kernel of 6-mm full width at half-maximum using the 3dmerge function from AFNI. Following smoothing, denoising was performed using a locally developed Nipype pipeline. In this pipeline, we fitted a GLM to the fMRI BOLD voxel time series. The GLM modelled a) the effect of six movement parameters (translation in x, y, and z directions, and rotation in yaw, roll, and pitch directions) and their first derivative, b) constant and linear trends using zero-th-order and first-order Legendre polynomials, c) principal components of signals in the WM and CSF masks, and d) motion outlier data points due to FD > 0.5 or DVARS > 1.5. Then, a bandpass filter in the range of 0.008 -0.09 Hz was applied on the residuals of the model to extract low-frequency fluctuations of the BOLD signal. Finally, we utilised the Schafer 100 ROI parcellation (Schaefer et al., 2018) and 19 subcortical ROIs to extract the averaged BOLD signals inside each ROI (Supplementary Figure S1a, b). To ensure generalizability and robustness of the pipeline, we varied the regression of the global signal and the regression of task events (Supplementary Material). For the task regression, we included task-related regressors in the GLM by convolving a boxcar representation of experimental events with the hemodynamic response function. The experimental events modelled were the go/no-go trials, participants’ button responses to go/no-go trials, the pre-probe blank screen, the question screens related to the probes, and participants’ button presses for each probe. The preprocessing steps described above were selected to replicate previous explorations of time-varying functional connectivity and its relationship to MB ^9,40^.

#### Global Signal Amplitude Analysis

We extracted the instantaneous global signal amplitude by applying the Hilbert transformation at each ROI timepoint as:

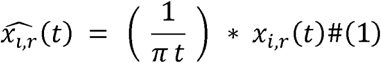

where * indicates a convolution operator. Using this transformation, we produced an analytical signal for each regional time series as:

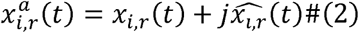

where 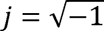. For each mental report and vigilance level, we extracted the last 6 TR (∼ 10s) preceding the probes of the amplitude of the analytical signal. To account for the potential lag of the BOLD response, we also reanalysed the shifted global signal amplitude time series by 1 and 2 TR (Supplementary Material).

#### Time-varying Functional Connectivity Estimation

We used phase-based coherence analysis to extract ROI-ROI functional connectivity (FC) at each TR of the scanning session. For every participant *i*, we z-scored each of the ROI *r* time series and applied a band-pass filter of 0.01-0.04 Hz to isolate the low-oscillatory BOLD signal of interest. Then, we calculated the Hilbert transformation. From the Hilbert analytical signal, the instantaneous phase of each time series can be estimated as:

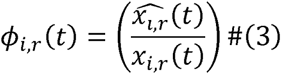

To ensure that the phase time series were accurately represented and free from artefacts, we applied Empirical Mode Decomposition (EMD) to each voxel phase time series. EMD is a signal processing technique that decomposes a signal into its underlying frequency components, known as intrinsic mode functions (IMFs). At each iteration of the algorithm, EMD identifies the local extrema of the original signal, from which upper and lower envelopes are constructed using spline interpolation. The mean of these envelopes is calculated and subtracted from the original signal to extract the first intrinsic mode function (IMF). This procedure is iterated until the extracted signal meets the criteria for being an IMF. Therefore, at each iteration, the algorithm produces an IMF and the residual trend of the signal after the IMF is subtracted. As such, we kept the first oscillation and discarded the remaining IMFs.

Then, we warped the denoised phase signal *ϕ_i,r_*(*t*), to the [-*π*, *π*] interval and, naming the obtained signal as *θ_i,r_*(*t*), we calculated a connectivity measure for each pair of regions as the cosine of their phase difference. For example, the connectivity measure between regions *r* and *s* in subject *i* was defined as:

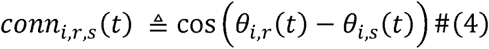

By this definition, completely synchronised time series lead to a connectivity value of 1, completely desynchronised time series produce a connectivity value of zero, and anticorrelated time series produce a connectivity measure of -1. Using this approach, we created a connectivity matrix of 119 x 119 (Supplementary Figure S1c), representing the sum of cortical and subcortical parcels, at each time point *t* for each subject *i* that we called *C_i_*(*t*):

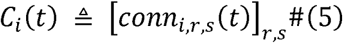

#### Connectivity Pattern Extraction (K-Means Clustering)

We applied clustering to identify representative connectivity patterns that summarise the dominant modes of variation in time-varying functional connectivity. First, given that the connectivity matrices described above are symmetrical, we extracted the upper triangular part of the matrix for every participant at every TR. By aggregating all connectivity vectors, we created a new matrix where each row represents the phase-based connectivity at a single time point (TR) for a participant, and each column represents the phase-based connectivity values between different ROIs (Supplementary Figure S1d). We applied k-means clustering to the resulting connectivity matrix, using the Manhattan distance as the distance metric (Supplementary Figure S1e). Manhattan distance has been repeatedly utilised to investigate how the functional connectome organises over time during rest ^40,99^ and has successfully linked behaviour to connectivity patterns ^43^. To ensure methodological coherence in neurobehavioral coupling, we opted for the same distance metric for both clustering the functional connectome and estimating its distance to each cluster. This approach ensures that clustering and subsequent characterisation of the connectome are based on consistent criteria. To our knowledge, previous studies that have both clustered and linked clusters to the connectome using the same metric have utilised the Manhattan distance. To assess the generalizability of the connectivity patterns to the distance choice, we also clustered the data using cosine and Euclidean distance (Supplementary Material). Clustering produced *k* distinct centroids that represented brain connectivity matrices, along with the occurrence frequency of each configuration for every participant. To ensure the generalizability of the centroids, we varied the number of centroids from *k* = 3 to *k* = 8. We initialised clusters using the “K-means++” method and applied clustering 200 times ^100^. Finally, we ordered the patterns by their standard deviation and estimated the frequency of each centroid by counting the number of matrices assigned to each centroid for each participant.

Then, we estimated the optimal number of centroids by calculating the inter-pattern correlation variance across all *k*. We further assessed centroid similarity across *k* by correlating all patterns with the first and last centroid obtained at *k* = 5. If the centroids were stable when ordered, the first centroid at *k* = 5 is expected to exhibit the strongest correlation with the first centroid across *k* = 3–8. Similarly, the last centroid at *k* = 5 should have shown the strongest correlation with the last centroid across the same range of *k*.

#### Coupling Between Connectivity Patterns and Experience Sampling

To assess the relationship between each of the connectivity patterns created by our clustering approach and the experience-sampling reports, we extracted the upper triangular parts of the connectivity matrices 6 TR (∼10s) preceding each report. As the goal of this study is to eventually link electrophysiology with time-varying connectivity, the choice of the window (10s) was motivated by previous results showing that mental states correlated with distinct neural and electrophysiological activity from that window ^7–9^. For every distance metric used during k-means clustering (Euclidean, Manhattan or cosine), we estimated the distance of the extracted connectivity vectors with each of the *k* centroids (Supplementary Figure S1f-g). Higher values suggest that, during that TR, the connectome is organised into a configuration *dissimilar* (or more distant) to the examined connectivity pattern. Inversely, lower values indicate larger *similarity* (or proximity) to the examined connectivity pattern. In the main text, we present results originating from a *k* = 5, as that number of clusters showed the highest inter-pattern correlation covariance. To ensure generalizability, we replicated our analysis across all clustering *k* (3-8) (Supplementary Material). Additionally, to account for a potential effect of the hemodynamic response, we re-analysed our data using the shifted versions of the connectivity matrices with time lags ranging from zero (four TRs before the probe) to two (two TRs pre-probe and two TRs post-probe matrices) (Supplementary Material). Across all analyses, model significance was corrected for the total number of clusters examined using the Bonferroni correction.

#### Coupling Between Connectivity Patterns and SW-like Activity

To understand how SW-like activity would correlate with connectivity patterns, we utilised Canonical Correlations Analysis (CCA). CCA is a multivariate statistical technique used to identify the relationships between two sets of variables ^101–103^. CCA reduces two sets of variables, X and Y, into lower-dimensional representations called canonical components. These components are created as linear combinations of the original variables. The method then selects the combinations in such a way that the canonical components from X and Y are maximally correlated with each other. To put it simply, CCA uncovers correlations in latent representations of sets of variables.

In our dataset, one set of variables refers to the SW markers per EEG channel (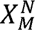, where *M* represents SW activity at every EEG channel, averaged over 10s preceding probes, and *N* represents the total number of experience-sampling reports across participants), and the other set refers to the distances of the connectivity matrix of the underlying fMRI patterns (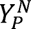 where *P* represents the average distance of a connectivity matrix from each of the *k* identified patterns and *N* represents the total number of experience-sampling reports across participants) (Figure 6a-b). The CCA dataset was comprised of 25 participants (mean age = 23.08 years; 15 females), as data from additional participants were excluded due to EEG signal corruption arising from the simultaneous EEG–fMRI acquisition. Overall, the CCA dataset included 931 observations.

A regularisation penalty cap L1 and cap L2 were added to each set of variables, respectively, to ensure numerical stability and avoid multicollinearity. These parameters were selected using 10-fold cross-validation, in which the data were repeatedly split into ten subsets, with the model trained on nine subsets and evaluated on the remaining subset. Performance was assessed based on the correlation between canonical variates estimated on held-out data. Given the exploratory nature of this analysis and the small number of subjects with uncorrupted EEG-fMRI data, we did not retain the nested probe structure. To assess the significance of each canonical component, a permutation approach was adopted ^103^. We permuted the EEG variable 5,000 times and refitted the CCA model, extracting an empirical null distribution for the correlations. The resulting empirical p-values were FDR-corrected for the number of centroids tested per model. Finally, to assess the contribution of each variable to the canonical component, we estimated the correlations between the canonical components and each variable.

### Statistical Analysis

Statistical analysis was performed using generalised mixed-effects models and non-parametric tests. For the mixed-effects models, the family used to model the dependent variable varied depending on the distribution of the dependent variable. For a binary variable, the data were modelled using a Binomial distribution with a logit link function. In case of positive variables, data were modelled using a Gamma distribution. Finally, the remaining tests were conducted using a Gaussian distribution with an identity function. We allowed intercepts and slopes to vary freely, using subject ID as a random effect. In the case of a singular fit, we first iteratively removed slopes and subsequently intercepts. Models containing main effects of mental states and vigilance levels, as well as their interactions, were contrasted with null models that included only random intercepts. Model comparison was conducted using a chi-square difference test, with model evidence further quantified via Bayes factors (relative evidence based on Bayesian Information Criterion). Pairwise multiple contrasts of significant models were corrected using FDR correction. Since confidence limits are not estimated using the FDR correction, we provide the Bonferroni-corrected confidence limits. For all analyses, descriptive statistics of all fixed parameters are included to ensure transparency and facilitate replication based on our own distribution parameters. All statistical analyses were conducted using two-tailed tests.

### Software

Analyses were conducted using the following programming languages: Python v3.6, Python v3.12, R 4.3.3, and MATLAB. fMRI preprocessing and denoising were performed using the ‘fMRIPrep’ pipeline ^84^, ‘Nipypè v1.9.2 ^85^ and custom Python functions. Global signal amplitude and time-varying functional connectivity extraction were conducted using custom Python functions and the ‘EMD’ library v0.8.1 ^104^. Clustering was conducted using the ‘pyclustering’ v0.10.1 library ^105^. Mixed-effects models were fit using the ‘lme4’ v1.1.35 package ^106^. Contrasts were estimated using the ‘emmeans’ v1.11 package ^107^. Model comparison was conducted using the ‘flexplot’ v0.7.4 package. Canonical correlations were estimated using the ‘ccà v1.2.2 package ^108^.

## DATA AVAILABILITY

All behavioural dataframes, ROI time courses, global signal dataframes, k-means models, connectome distance dataframes and canonical correlations models across all our analysis parameters can be found in https://doi.org/10.58119/ULG/4LEK9M.

## CODE AVAILABILITY

All analysis was conducted in Python, R and MATLAB. An archived version of the code at the time of submission can be found at https://gitlab.uliege.be/poc/mind-blanking-monash-collab.

## ACKNOWLEDGEMENTS

Analysis was conducted at the GIGA-CRC-Human Imaging platform of ULiège, Belgium. We would like to thank Dr. Sepehr Mortaheb and Nikolaos-Ioannis Simos for their insightful comments during analysis. The authors acknowledge the facilities and scientific and technical assistance of the National Imaging Facility (NIF), a National Collaborative Research Infrastructure Strategy (NCRIS) capability at Monash Biomedical Imaging (MBI), a Technology Research Platform at Monash University. The authors acknowledge the facilities and scientific and technical assistance of Monash Biomedical Imaging (MBI), a Technology Research Platform at Monash University.

At the time of the research, PAB was an FNRS Research Fellow. AD is an FNRS Research Associate. This work was supported by the Belgian Fund for Scientific Research (FRS-FNRS), the European Union’s Horizon 2020 Research and Innovation Marie Skłodowska-Curie RISE programme 15 NeuronsXnets (grant agreement 101007926), the European Cooperation in Science and Technology 16 COST Action (CA18106), the Léon Fredericq Foundation, and the University Hospital of Liège. TA, NT, and AK were supported by Australian Research Council (DP240102680) and National Health Medical Research Council (GNT1183280, GNT2037172). NT and AK were supported by Japan Society for the Promotion of Science Grant-in-Aid for Transformative Research Areas (A) (23H04829, 23H04830) and Japan Science and Technology (JST) Moonshot R&D Grant (JPMJMS2295-14). NT was supported by Theoretical Sciences Visiting Program, Okinawa Institute of Science and Technology. The funders had no role in study design, data collection and analysis, decision to publish, or preparation of the manuscript.

## AUTHOR CONTRIBUTIONS

<u>PAB</u>: Conceptualisation, Data Curation, Formal Analysis, Methodology, Project Administration, Software, Validation, Visualisation, Writing – original draft. <u>AK</u>: Data Curation, Formal Analysis, Methodology, Formal Analysis, Software, Resources, Writing -review & editing. <u>NT:</u> Methodology, Writing – review & editing. <u>TA</u>: Formal Analysis, Funding Acquisition, Methodology, Software, Supervision, Writing -review & editing. <u>AD</u>: Conceptualisation, Methodology, Supervision, Writing – original draft, Writing – review & editing.

## COMPETING INTERESTS

The authors declare no competing interests.

## Notes

https://doi.org/10.58119/ULG/4LEK9M

### Competing Interest Statement

The authors have declared no competing interest.

### Summary of Updates

We have made the following adjustments: 1. More clear definition and operationalization of mind-blanking. This includes control analysis for both MB and No Recall reports 2. Control analysis to validate the MB is not micro-sleep 3. Control analysis to account for gender 4. Included more nuanced discussion on the implications of our results 5. Included more nuanced discussion on the justification of our methodological choices

