## Supplementary Material for "Distinct cortical profiles underlie the common reportability of thought-free experiences"

Anikó Kusztor^3,4^

Naotsugu Tsuchiya^3,4,5,6,7^

Thomas Andrillon^8^

Athena Demertzi^1,2*^

^1^Physiology of Cognition Lab, GIGA CRC-Human Imaging, Allée du 6 Août 8 (B30), 4000, University of Liège, Belgium

^2^Psychology and Neuroscience of Cognition Research Unit, Place des Orateurs 3 (B33), 4000, University of Liège, Belgium

^3^School of Psychological Science, Monash University, Melbourne, Victoria 3168, Australia

^4^Turner Institute for Brain and Mental Health, Faculty of Medicine, Nursing and Health Science, Monash University, Victoria, Australia

^5^Center for Information and Neural Networks (CiNet), National Institute of Information and Communications Technology (NICT), Osaka, Japan

^6^Advanced Telecommunications Research Computational Neuroscience Laboratories, Kyoto, Japan.

^7^Theoretical Sciences Visiting Program (TSVP), Okinawa Institute of Science and Technology Graduate University, Onna, 904-0495, Japan

^8^Institut du Cerveau—Paris Brain Institute—ICM, Inserm, Sorbonne Université, CNRS, APHP, Hôpital de la Pitié Salpêtrière, 75013 Paris, France

These authors contributed equally: Paradeisios Alexandros Boulakis, Anikó Kusztor

These authors jointly supervised this work: Athena Demertzi, Thomas Andrillon

**Supplementary Table 1.** Descriptive statistics of mental state, alertness and mental state frequencies stratified by alertness.

| **Report** | **Mean** | **Median** | **Min** | **Max** | **SD** | **IQR1** | **IQR3** | **Skewness** | **Kurtosis** |
| --- | --- | --- | --- | --- | --- | --- | --- | --- | --- |
| ONTASK | 0.486 | 0.462 | 0 | 1.0 | 0.064 | 0.253 | 0.288 | 0.644 | 0.294 |
| MW | 0.389 | 0.404 | 0 | 0.8 | 0.047 | 0.217 | 0.229 | 0.534 | 0.082 |
| BLANK | 0.125 | 0.081 | 0 | 0.5 | 0.019 | 0.136 | 0.006 | 0.204 | 1.063 |
| SLEEPY | 0.428 | 0.425 | 0.000 | 0.925 | 0.089 | 0.298 | 0.192 | 0.677 | 0.074 |
| ALERT | 0.572 | 0.575 | 0.075 | 1.000 | 0.089 | 0.298 | 0.323 | 0.808 | -0.074 |
| SLEEPY: ONTASK | 0.105 | 0.076 | 0 | 0.514 | 0.014 | 0.119 | 0.000 | 0.135 | 1.666 |
| SLEEPY: OFFTASK | 0.221 | 0.197 | 0 | 0.667 | 0.039 | 0.197 | 0.057 | 0.353 | 0.654 |
| SLEEPY: MINDBLANK | 0.102 | 0.035 | 0 | 0.500 | 0.018 | 0.135 | 0.000 | 0.144 | 1.370 |
| ALERT: ONTASK | 0.381 | 0.372 | 0 | 1.000 | 0.061 | 0.247 | 0.181 | 0.551 | 0.613 |
| ALERT: OFFTASK | 0.168 | 0.136 | 0 | 0.700 | 0.023 | 0.153 | 0.042 | 0.250 | 1.215 |
| ALERT: MINDBLANK | 0.023 | 0.000 | 0 | 0.300 | 0.003 | 0.051 | 0.000 | 0.026 | 4.123 |
| SD = standard deviation; IQR1 = interquartile range – 25% percentile; IQR3 = interquartile range – 75% percentile | | | | | | | | | |

**Supplementary Table S2.** Descriptive statistics of mental state and alertness level report frequencies pre-aggregation.

| **Report** | **Mean** | **Median** | **Min** | **Max** | **SD** | **IQR1** | **IQR3** | **Skewness** | **Kurtosis** |
| --- | --- | --- | --- | --- | --- | --- | --- | --- | --- |
| ONTASK | 0.486 | 0.462 | 0 | 1.000 | 0.253 | 0.288 | 0.644 | 0.294 | -0.694 |
| MW | 0.389 | 0.404 | 0 | 0.800 | 0.217 | 0.229 | 0.534 | 0.082 | -0.781 |
| BLANK | 0.100 | 0.065 | 0 | 0.444 | 0.115 | 0.000 | 0.135 | 1.235 | 0.682 |
| NO RECALL | 0.025 | 0.000 | 0 | 0.176 | 0.043 | 0.000 | 0.032 | 2.281 | 5.1 |
| EXSLEEPY | 0.133 | 0.046 | 0.000 | 0.675 | 0.181 | 0.000 | 0.199 | 1.306 | 0.605 |
| SLEEPY | 0.295 | 0.275 | 0.000 | 0.639 | 0.186 | 0.175 | 0.425 | -0.051 | -1.152 |
| ALERT | 0.484 | 0.478 | 0.075 | 1.000 | 0.261 | 0.283 | 0.628 | 0.411 | -0.791 |
| EXALERT | 0.088 | 0.050 | 0.000 | 0.350 | 0.104 | 0.000 | 0.138 | 1.134 | 0.005 |
| SD = standard deviation; IQR1 = interquartile range – 25% percentile; IQR3 = interquartile range – 75% percentile | | | | | | | | | |

**Supplementary Table S3.** MB and Sleepiness were reported less frequently compared to other reports

| **Contrast** | **Statistic** | **pFDR** | **Rank-biserial r** | **CI** |
| --- | --- | --- | --- | --- |
| ONTASK-OFFTASK | -1.34 | 1.8e-01 | 0.21 | [-0.05, 0.44] |
| ONTASK-BLANK | -6.28 | 9.9e-10 | 0.8 | [0.69, 0.88] |
| OFFTASK-BLANK | -4.94 | 1.1e-06 | 0.69 | [0.53, 0.8] |
| SLEEPY-ALERT | 2.02 | 4.2e-02 | -0.27 | [-0.49, -0.02] |
| CI = confidence intervals 95%; FDR = false discovery rate | | | | |

**Supplementary Table S4.** MB and OFFTASK reports increase during sleepiness, while ONTASK reports increase during alertness

| **Contrast** | **Odds Ratio** | **SE** | **Z ratio** | **CL** | **pFDR** |
| --- | --- | --- | --- | --- | --- |
| ONTASK: SLEEPY / ALERT | 0.15 | 0.02 | -11.67 | [0.11, 0.21] | 5.7e-31 |
| OFFTASK: SLEEPY/ALERT | 2.64 | 0.4 | 6.5 | [1.7, 3.53] | 7.9e-11 |
| BLANK: SLEEPY / ALERT | 8.03 | 2.1 | 7.96 | [4.8, 13.4] | 1.7e-15 |
| SE = standard error; CL = confidence limits 95 %; FDR = false discovery rate | | | | | |

**Supplementary Table S5.** Descriptive statistics of mental state and alertness level reaction times to probes.

| **Report** | **Mean** | **Median** | **Min** | **Max** | **SD** | **IQR1** | **IQR3** | **Skewness** | **Kurtosis** |
| --- | --- | --- | --- | --- | --- | --- | --- | --- | --- |
| ONTASK | 1.649 | 1.143 | 0.272 | 7.690 | 1.397 | 0.687 | 2.053 | 1.732 | 2.759 |
| OFFTASK | 1.980 | 1.614 | 0.295 | 7.826 | 1.322 | 1.035 | 2.583 | 1.491 | 2.436 |
| BLANK | 2.793 | 2.330 | 0.325 | 7.886 | 1.760 | 1.501 | 3.567 | 1.087 | 0.676 |
| SLEEPY | 2.202 | 1.837 | 0.295 | 7.886 | 1.468 | 1.133 | 2.876 | 1.338 | 1.772 |
| ALERT | 1.713 | 1.213 | 0.272 | 7.690 | 1.424 | 0.723 | 2.201 | 1.762 | 3.039 |
| SD = standard deviation; IQR1 = interquartile range – 25% percentile; IQR3 = interquartile range – 75% percentile | | | | | | | | | |

**Supplementary Table S6**. Main effects and interaction contrasts describing the relationship between reaction times, mental states, and alertness levels.

| **Contrast** | **Estimate** | **SE** | **Z ratio** | **CL** | **p-value** |
| --- | --- | --- | --- | --- | --- |
| ONTASK - OFFTASK | 0.03 | 0.02 | 1.25 | [-0.03, 0.08] | 2.13e-01 |
| **ONTASK - BLANK** | **0.18** | **0.03** | **7.11** | **[0.12, 0.25]** | **3.55e-12** |
| **OFFTASK - BLANK** | **0.16** | **0.02** | **6.35** | **[0.10, 0.22]** | **3.21e-10** |
| SLEEPY - ALERT | 0.03 | 0.02 | 1.51 | [-0.01, 0.07] | 1.31e-01 |
| SLEEPY: ONTASK - OFFTASK | 0.00 | 0.03 | 0.03 | [-0.08, 0.08] | 9.79e-01 |
| **SLEEPY: ONTASK - BLANK** | **0.09** | **0.04** | **2.59** | **[0.01, 0.18]** | **1.45e-02** |
| **SLEEPY: OFFTASK - BLANK** | **0.09** | **0.03** | **3.27** | **[0.02, 0.16]** | **3.22e-03** |
| ALERT: ONTASK - OFFTASK | 0.05 | 0.03 | 1.92 | [-0.01, 0.12] | 5.47e-02 |
| **ALERT: ONTASK - BLANK** | **0.28** | **0.04** | **7.24** | **[0.18, 0.37]** | **1.38e-12** |
| **ALERT: OFFTASK - BLANK** | **0.22** | **0.04** | **5.44** | **[0.12, 0.32]** | **8.18e-08** |
| SE = standard error; CL = confidence limits 95 %; FDR = false discovery rate | | | | | |

**Supplementary Table S7.** Descriptive statistics of global signal amplitude across mental states and alertness levels.

| **Report** | **Mean** | **Median** | **Min** | **Max** | **SD** | **IQR1** | **IQR3** | **Skewness** | **Kurtosis** |
| --- | --- | --- | --- | --- | --- | --- | --- | --- | --- |
| ONTASK | 0.514 | 0.441 | 0.003 | 4.030 | 0.363 | 0.264 | 0.661 | 1.981 | 7.237 |
| OFFTASK | 0.538 | 0.465 | 0.005 | 3.468 | 0.370 | 0.278 | 0.709 | 1.888 | 6.988 |
| BLANK | 0.599 | 0.509 | 0.030 | 2.369 | 0.398 | 0.327 | 0.781 | 1.347 | 2.224 |
| SLEEPY | 0.597 | 0.512 | 0.005 | 3.468 | 0.421 | 0.299 | 0.781 | 1.695 | 4.734 |
| ALERT | 0.488 | 0.421 | 0.003 | 4.030 | 0.323 | 0.262 | 0.638 | 1.830 | 7.268 |
| SD = standard deviation; IQR1 = interquartile range – 25% percentile; IQR3 = interquartile range – 75% percentile | | | | | | | | | |

**Supplementary Table S8**. Main effect and interaction contrasts describing the relationship between global signal amplitude, mental states and alertness levels.

| **Contrast** | **Estimate** | **SE** | **Z ratio** | **CL** | **p-value** |
| --- | --- | --- | --- | --- | --- |
| ONTASK - OFFTASK | -0.00 | 0.02 | -0.17 | [-0.04, 0.04] | 8.68e-01 |
| **ONTASK - BLANK** | **-0.09** | **0.03** | **-3.02** | **[-0.15, -0.02]** | **4.43e-03** |
| **OFFTASK - BLANK** | **-0.08** | **0.03** | **-2.97** | **[-0.15, -0.02]** | **4.43e-03** |
| **SLEEPY - ALERT** | **0.12** | **0.02** | **5.71** | **[0.08, 0.17]** | **1.13e-08** |
| SLEEPY: ONTASK - OFFTASK | 0.04 | 0.03 | 1.66 | [-0.02, 0.11] | 1.46e-01 |
| SLEEPY: ONTASK - BLANK | -0.04 | 0.03 | -1.31 | [-0.12, 0.03] | 1.91e-01 |
| **SLEEPY: OFFTASK - BLANK** | **-0.09** | **0.03** | **-3.13** | **[-0.15, -0.02]** | **5.29e-03** |
| **ALERT: ONTASK - OFFTASK** | **-0.05** | **0.02** | **-2.40** | **[-0.10, 0.00]** | **2.49e-02** |
| **ALERT: ONTASK - BLANK** | **-0.13** | **0.05** | **-2.78** | **[-0.24, -0.02]** | **1.61e-02** |
| ALERT: OFFTASK - BLANK | -0.08 | 0.05 | -1.64 | [-0.19, 0.04] | 1.01e-01 |
| SE = standard error; CL = confidence limits 95 %; FDR = false discovery rate | | | | | |

**Supplementary Table S9.** Descriptive statistics of brain pattern occurrence frequency.

| **Brain Pattern** | **Mean** | **Median** | **Min** | **Max** | **SD** | **IQR1** | **IQR3** | **Skewness** | **Kurtosis** |
| --- | --- | --- | --- | --- | --- | --- | --- | --- | --- |
| Pattern 1 | 0.209 | 0.184 | 0.094 | 0.410 | 0.071 | 0.167 | 0.256 | 0.961 | 0.679 |
| Pattern 2 | 0.162 | 0.155 | 0.061 | 0.313 | 0.055 | 0.120 | 0.188 | 0.489 | 0.012 |
| Pattern 3 | 0.166 | 0.161 | 0.071 | 0.392 | 0.062 | 0.114 | 0.192 | 1.190 | 2.539 |
| Pattern 4 | 0.132 | 0.126 | 0.049 | 0.293 | 0.051 | 0.091 | 0.158 | 0.798 | 0.772 |
| Pattern 5 | 0.331 | 0.320 | 0.134 | 0.679 | 0.132 | 0.228 | 0.427 | 0.532 | -0.409 |
| SD = standard deviation; IQR1 = interquartile range – 25% percentile; IQR3 = interquartile range – 75% percentile | | | | | | | | | |

**Supplementary Table S10.** Contrast analysis of the differences in brain pattern occurrence frequencies.

| **Contrast** | **Estimate** | **SE** | **Z ratio** | **CL** | **p-value** |
| --- | --- | --- | --- | --- | --- |
| **Pattern 1 - Pattern 2** | **0.05** | **0.02** | **2.54** | **[-0.01, 0.10]** | **2.05e-02** |
| **Pattern 1 - Pattern 3** | **0.04** | **0.02** | **2.33** | **[-0.01, 0.09]** | **3.05e-02** |
| **Pattern 1 - Pattern 4** | **0.08** | **0.02** | **4.18** | **[0.02, 0.13]** | **9.96e-05** |
| **Pattern 1 - Pattern 5** | **-0.12** | **0.02** | **-6.71** | **[-0.17, -0.07]** | **9.69e-10** |
| Pattern 2 - Pattern 3 | -0.00 | 0.02 | -0.21 | [-0.06, 0.05] | 8.36e-01 |
| Pattern 2 - Pattern 4 | 0.03 | 0.02 | 1.64 | [-0.02, 0.08] | 1.14e-01 |
| **Pattern 2 - Pattern 5** | **-0.17** | **0.02** | **-9.24** | **[-0.22, -0.12]** | **1.20e-15** |
| Pattern 3 – Pattern 4 | 0.03 | 0.02 | 1.85 | [-0.02, 0.09] | 8.25e-02 |
| **Pattern 3 - Pattern 5** | **-0.17** | **0.02** | **-9.04** | **[-0.22, -0.11]** | **2.74e-15** |
| **Pattern 4 – Pattern 5** | **-0.20** | **0.02** | **-10.89** | **[-0.25, -0.15]** | **1.20e-19** |
| SE = standard error; CL = confidence limits 95 %; FDR = false discovery rate | | | | | |

**Supplementary Table S11.** Descriptive statistics of distances of time-varying FC during mental states and alertness levels from the brain patterns.

| **Pattern** | **Report** | **Mean** | **Median** | **Min** | **Max** | **SD** | **IQR1** | **IQR3** | **Skewness** | **Kurtosis** |
| --- | --- | --- | --- | --- | --- | --- | --- | --- | --- | --- |
| P1 | ONTASK | 4440.301 | 4430.541 | 3910.006 | 6244.807 | 190.954 | 4333.560 | 4519.712 | 1.568 | 7.870 |
| P1 | OFFTASK | 4432.874 | 4425.496 | 3960.780 | 5655.372 | 176.840 | 4335.004 | 4509.746 | 1.184 | 5.047 |
| P1 | BLANK | 4424.426 | 4417.706 | 3893.356 | 5190.178 | 176.026 | 4321.109 | 4516.312 | 0.492 | 1.428 |
| P1 | SLEEPY | 4438.984 | 4427.483 | 3981.135 | 5401.561 | 174.620 | 4337.988 | 4519.839 | 0.946 | 2.860 |
| P1 | ALERT | 4433.034 | 4426.717 | 3893.356 | 6244.807 | 190.201 | 4328.146 | 4512.897 | 1.543 | 8.131 |
| P2 | ONTASK | 4473.548 | 4453.604 | 3934.488 | 6072.208 | 174.012 | 4379.748 | 4536.416 | 1.949 | 9.536 |
| P2 | OFFTASK | 4463.692 | 4449.971 | 3959.624 | 5601.975 | 156.508 | 4376.045 | 4530.844 | 1.180 | 4.998 |
| P2 | BLANK | 4463.822 | 4445.065 | 4129.599 | 5156.260 | 139.840 | 4377.811 | 4533.040 | 0.866 | 1.870 |
| P2 | SLEEPY | 4464.524 | 4448.148 | 3959.624 | 5312.159 | 154.505 | 4373.013 | 4534.082 | 0.934 | 2.488 |
| P2 | ALERT | 4471.441 | 4453.587 | 3934.488 | 6072.208 | 169.743 | 4382.107 | 4534.557 | 1.991 | 10.330 |
| P3 | ONTASK | 4460.675 | 4435.260 | 4045.824 | 6133.537 | 178.734 | 4362.456 | 4520.920 | 2.075 | 9.415 |
| P3 | OFFTASK | 4456.744 | 4437.679 | 4033.137 | 5587.687 | 159.734 | 4365.919 | 4523.906 | 1.436 | 5.230 |
| P3 | BLANK | 4445.873 | 4429.600 | 4057.406 | 5195.973 | 155.860 | 4349.987 | 4522.042 | 0.890 | 2.293 |
| P3 | SLEEPY | 4464.324 | 4442.391 | 4052.211 | 5387.385 | 163.552 | 4368.209 | 4535.896 | 1.176 | 3.071 |
| P3 | ALERT | 4452.404 | 4430.101 | 4033.137 | 6133.537 | 172.568 | 4359.073 | 4512.238 | 2.141 | 10.519 |
| P4 | ONTASK | 4472.174 | 4441.831 | 4051.184 | 6232.415 | 169.202 | 4380.343 | 4528.024 | 2.470 | 12.697 |
| P4 | OFFTASK | 4461.935 | 4436.054 | 4077.725 | 5640.300 | 157.344 | 4371.709 | 4521.159 | 1.952 | 7.975 |
| P4 | BLANK | 4460.362 | 4438.273 | 4127.010 | 5278.435 | 149.941 | 4375.651 | 4527.183 | 1.130 | 3.347 |
| P4 | SLEEPY | 4460.565 | 4434.407 | 4051.184 | 5456.137 | 159.473 | 4369.433 | 4522.228 | 1.536 | 4.883 |
| P4 | ALERT | 4471.223 | 4443.381 | 4110.607 | 6232.415 | 164.525 | 4381.617 | 4527.515 | 2.585 | 13.847 |
| P5 | ONTASK | 4414.449 | 4462.178 | 3455.978 | 5676.460 | 243.877 | 4296.275 | 4566.336 | -0.610 | 1.626 |
| P5 | OFFTASK | 4416.752 | 4465.423 | 3529.340 | 5583.751 | 239.234 | 4292.084 | 4571.685 | -0.617 | 0.978 |
| P5 | BLANK | 4425.492 | 4468.797 | 3587.530 | 5212.208 | 226.461 | 4315.341 | 4565.511 | -0.680 | 1.397 |
| P5 | SLEEPY | 4407.826 | 4462.825 | 3455.978 | 5290.038 | 247.382 | 4277.344 | 4566.728 | -0.699 | 0.859 |
| P5 | ALERT | 4422.983 | 4465.291 | 3477.422 | 5676.460 | 234.479 | 4308.900 | 4570.062 | -0.549 | 1.769 |
| SD = standard deviation; IQR1 = interquartile range – 25% percentile; IQR3 = interquartile range – 75% percentile | | | | | | | | | | |

**Supplementary Table S12.** Main effect and interaction contrasts describing the relationship between patterns, mental states and alertness levels.

| Pattern | Contrast | Estimate | SE | D.F. | CL | T-statistic | P-value |
| --- | --- | --- | --- | --- | --- | --- | --- |
| P1 | ONTASK - OFFTASK | 7.24 | 5.1 | 8122.91 | [-4.97, 19.46] | 1.42 | 1.56e-01 |
| **P1** | **ONTASK - BLANK** | **48.1** | **8.31** | **8370.65** | **[28.19, 68.00]** | **5.79** | **2.24e-08** |
| **P1** | **OFFTASK - BLANK** | **40.85** | **8.17** | **8402.03** | **[21.30, 60.41]** | **5** | **8.70e-07** |
| **P1** | **SLEEPY - ALERT** | **15.38** | **6.37** | **7630.23** | **[2.89, 27.88]** | **2.41** | **1.58e-02** |
| P1 | SLEEPY: ONTASK - OFFTASK | 1.5 | 7.86 | 8337.07 | [-17.31, 20.31] | 0.19 | 8.48e-01 |
| P1 | SLEEPY: ONTASK - BLANK | 18.02 | 9.39 | 8258.42 | [-4.47, 40.50] | 1.92 | 8.26e-02 |
| P1 | SLEEPY: OFFTASK - BLANK | 16.51 | 8.02 | 8380.86 | [-2.68, 35.71] | 2.06 | 8.26e-02 |
| **P1** | **ALERT: ONTASK - OFFTASK** | **12.98** | **6.06** | **8332.72** | **[-1.53, 27.49]** | **2.14** | **3.22e-02** |
| **P1** | **ALERT: ONTASK - BLANK** | **78.18** | **13.64** | **8432.54** | **[45.53, 110.83]** | **5.73** | **3.05e-08** |
| **P1** | **ALERT: OFFTASK - BLANK** | **65.19** | **14.21** | **8412.95** | **[31.16, 99.23]** | **4.59** | **6.84e-06** |
| P2 | ONTASK - OFFTASK | 11.27 | 4.56 | 7474.49 | [0.35, 22.20] | 2.47 | 2.03e-02 |
| **P2** | **ONTASK - BLANK** | **24.1** | **7.45** | **8102.02** | **[6.27, 41.94]** | **3.24** | **3.65e-03** |
| P2 | OFFTASK - BLANK | 12.83 | 7.32 | 8190.08 | [-4.70, 30.36] | 1.75 | 7.97e-02 |
| P2 | SLEEPY - ALERT | 0.96 | 5.69 | 6474.61 | [-10.19, 12.11] | 0.17 | 8.66e-01 |
| **P2** | **SLEEPY: ONTASK - OFFTASK** | **18.69** | **7.04** | **8017.14** | **[1.84, 35.53]** | **2.66** | **2.38e-02** |
| P2 | SLEEPY: ONTASK - BLANK | 16.07 | 8.41 | 7810.65 | [-4.06, 36.20] | 1.91 | 8.39e-02 |
| P2 | SLEEPY: OFFTASK - BLANK | -2.61 | 7.18 | 8131.38 | [-19.81, 14.58] | -0.36 | 7.16e-01 |
| P2 | ALERT: ONTASK - OFFTASK | 3.86 | 5.43 | 7996.57 | [-9.14, 16.85] | 0.71 | 4.78e-01 |
| **P2** | **ALERT: ONTASK - BLANK** | **32.14** | **12.23** | **8286.69** | **[2.86, 61.41]** | **2.63** | **2.58e-02** |
| **P2** | **ALERT: OFFTASK - BLANK** | **28.28** | **12.74** | **8223.13** | **[-2.23, 58.79]** | **2.22** | **3.97e-02** |
| P3 | ONTASK - OFFTASK | 9.01 | 4.7 | 7900.14 | [-2.24, 20.27] | 1.92 | 5.53e-02 |
| **P3** | **ONTASK - BLANK** | **33.95** | **7.67** | **8284.66** | **[15.59, 52.30]** | **4.43** | **2.89e-05** |
| **P3** | **OFFTASK - BLANK** | **24.94** | **7.53** | **8335.84** | **[6.90, 42.97]** | **3.31** | **1.40e-03** |
| P3 | SLEEPY - ALERT | 11.33 | 5.87 | 7201.78 | [-0.17, 22.83] | 1.93 | 5.35e-02 |
| P3 | SLEEPY: ONTASK - OFFTASK | 2.9 | 7.24 | 8232.28 | [-14.44, 20.24] | 0.4 | 6.89e-01 |
| **P3** | **SLEEPY: ONTASK - BLANK** | **24.12** | **8.65** | **8108.38** | **[3.39, 44.84]** | **2.79** | **8.01e-03** |
| **P3** | **SLEEPY: OFFTASK - BLANK** | **21.22** | **7.39** | **8301.42** | **[3.52, 38.92]** | **2.87** | **8.01e-03** |
| **P3** | **ALERT: ONTASK - OFFTASK** | **15.12** | **5.59** | **8223.29** | **[1.74, 28.50]** | **2.71** | **1.02e-02** |
| **P3** | **ALERT: ONTASK - BLANK** | **43.78** | **12.58** | **8388.43** | **[13.66, 73.89]** | **3.48** | **1.51e-03** |
| **P3** | **ALERT: OFFTASK - BLANK** | **28.65** | **13.11** | **8354.27** | **[-2.73, 60.04]** | **2.19** | **2.89e-02** |
| P4 | ONTASK - OFFTASK | 7.87 | 4.52 | 8006.77 | [-2.95, 18.69] | 1.74 | 1.23e-01 |
| P4 | ONTASK - BLANK | 15.66 | 7.37 | 8326.79 | [-1.98, 33.31] | 2.13 | 1.01e-01 |
| P4 | OFFTASK - BLANK | 7.79 | 7.24 | 8368.55 | [-9.55, 25.13] | 1.08 | 2.82e-01 |
| **P4** | **SLEEPY - ALERT** | **-16.42** | **5.65** | **7401.84** | **[-27.48, -5.35]** | **-2.91** | **3.64e-03** |
| P4 | SLEEPY: ONTASK - OFFTASK | 7.05 | 6.96 | 8283.23 | [-9.62, 23.72] | 1.01 | 4.67e-01 |
| P4 | SLEEPY: ONTASK - BLANK | 9.81 | 8.32 | 8180.78 | [-10.11, 29.74] | 1.18 | 4.67e-01 |
| P4 | SLEEPY: OFFTASK - BLANK | 2.76 | 7.11 | 8340.42 | [-14.25, 19.78] | 0.39 | 6.97e-01 |
| P4 | ALERT: ONTASK - OFFTASK | 8.69 | 5.37 | 8276.6 | [-4.17, 21.55] | 1.62 | 1.59e-01 |
| P4 | ALERT: ONTASK - BLANK | 21.51 | 12.09 | 8410.49 | [-7.44, 50.46] | 1.78 | 1.59e-01 |
| P4 | ALERT: OFFTASK - BLANK | 12.82 | 12.6 | 8383.37 | [-17.35, 42.99] | 1.02 | 3.09e-01 |
| P5 | ONTASK - OFFTASK | 4.3 | 6.52 | 8430.99 | [-11.32, 19.91] | 0.66 | 5.10e-01 |
| P5 | ONTASK - BLANK | 20.57 | 10.61 | 8468.31 | [-4.84, 45.98] | 1.94 | 1.58e-01 |
| P5 | OFFTASK - BLANK | 16.28 | 10.42 | 8470.95 | [-8.68, 41.23] | 1.56 | 1.78e-01 |
| **P5** | **SLEEPY - ALERT** | **-18.17** | **8.16** | **8323.18** | **[-34.17, -2.17]** | **-2.23** | **2.60e-02** |
| P5 | SLEEPY: ONTASK - OFFTASK | 5.15 | 10.03 | 8464.31 | [-18.86, 29.17] | 0.51 | 6.50e-01 |
| P5 | SLEEPY: ONTASK - BLANK | -5.44 | 11.99 | 8453.41 | [-34.16, 23.28] | -0.45 | 6.50e-01 |
| P5 | SLEEPY: OFFTASK - BLANK | -10.59 | 10.23 | 8469.26 | [-35.09, 13.90] | -1.04 | 6.50e-01 |
| P5 | ALERT: ONTASK - OFFTASK | 3.44 | 7.74 | 8464.12 | [-15.08, 21.97] | 0.45 | 6.56e-01 |
| **P5** | **ALERT: ONTASK - BLANK** | **46.59** | **17.39** | **8472** | **[4.94, 88.23]** | **2.68** | **2.22e-02** |
| **P5** | **ALERT: OFFTASK - BLANK** | **43.14** | **18.13** | **8471.54** | **[-0.28, 86.56]** | **2.38** | **2.61e-02** |
| SE = standard error; CL = confidence limits 95 %; FDR = false discovery rate; D.F. = degrees of freedom | | | | | | | |

**Supplementary Table S13**. Control analysis of the relationship between global signal, alertness and mental states by splitting MB into Blank and No Recall.

| **Contrast** | **Estimate** | **SE** | **Z ratio** | **CL** | **p-value** |
| --- | --- | --- | --- | --- | --- |
| ONTASK - MINDWANDER | 0.00 | 0.02 | -0.17 | [-0.049, 0.043] | 8.7e-01 |
| **ONTASK - MINDBLANK** | **-0.08** | **0.03** | **-2.77** | **[-0.161, -0.004]** | **2.0e-02** |
| ONTASK - NORECALL | -0.12 | 0.08 | -1.51 | [-0.321, 0.087] | 2.1e-01 |
| **MINDWANDER - MINDBLANK** | **-0.08** | **0.03** | **-2.72** | **[-0.157, -0.002]** | **2.0e-02** |
| MINDWANDER - NORECALL | -0.11 | 0.08 | -1.48 | [-0.317, 0.089] | 2.1e-01 |
| MINDBLANK - NORECALL | -0.03 | 0.08 | -0.43 | [-0.247, 0.178] | 8.0e-01 |
| SLEEPY - ALERT | 0.10 | 0.04 | 2.41 | [0.019, 0.185] | 1.6e-02 |
| SLEEPY: ONTASK - MINDWANDER | 0.04 | 0.03 | 1.66 | [-0.026, 0.114] | 1.9e-01 |
| SLEEPY: ONTASK - MINDBLANK | -0.04 | 0.03 | -1.27 | [-0.133, 0.047] | 3.1e-01 |
| SLEEPY: ONTASK - NORECALL | -0.04 | 0.05 | -0.73 | [-0.175, 0.099] | 5.6e-01 |
| **SLEEPY: MINDWANDER - MINDBLANK** | **-0.09** | **0.03** | **-2.93** | **[-0.166, -0.009]** | **2.1e-02** |
| SLEEPY: MINDWANDER - NORECALL | -0.08 | 0.05 | -1.67 | [-0.212, 0.047] | 1.9e-01 |
| SLEEPY: MINDBLANK - NORECALL | 0.01 | 0.05 | 0.10 | [-0.133, 0.143] | 9.2e-01 |
| **ALERT: ONTASK - MINDWANDER** | **-0.05** | **0.02** | **-2.40** | **[-0.105, 0.005]** | **4.9e-02** |
| **ALERT: ONTASK - MINDBLANK** | **-0.12** | **0.05** | **-2.52** | **[-0.25, 0.006]** | **4.9e-02** |
| ALERT: ONTASK - NORECALL | -0.20 | 0.15 | -1.35 | [-0.58, 0.188] | 2.7e-01 |
| ALERT: MINDWANDER - MINDBLANK | -0.07 | 0.05 | -1.43 | [-0.204, 0.061] | 2.7e-01 |
| ALERT: MINDWANDER - NORECALL | -0.15 | 0.15 | -1.00 | [-0.531, 0.239] | 3.8e-01 |
| ALERT: MINDBLANK - NORECALL | -0.07 | 0.15 | -0.49 | [-0.475, 0.327] | 6.3e-01 |
| SE = standard error; CL = confidence limits 95 %; FDR = false discovery rate | | | | | |

**Supplementary Table S14.** Main effect and interaction contrasts describing the relationship between patterns, mental states and alertness levels.

| Pattern | Contrast | Estimate | SE | D.F. | CL | T-statistic | P-value |
| --- | --- | --- | --- | --- | --- | --- | --- |
| P1 | ONTASK - MINDWANDER | 7.12 | 5.10 | 8139.958 | [-6.34, 20.57] | 1.40 | 2.4e-01 |
| **P1** | **ONTASK - MINDBLANK** | **56.88** | **8.71** | **8391.604** | **[33.88, 79.87]** | **6.53** | **4.2e-10** |
| P1 | ONTASK - NORECALL | -21.57 | 22.75 | 8465.047 | [-81.59, 38.45] | -0.95 | 3.4e-01 |
| **P1** | **MINDWANDER - MINDBLANK** | **49.76** | **8.59** | **8410.595** | **[27.1, 72.43]** | **5.79** | **2.1e-08** |
| P1 | MINDWANDER - NORECALL | -28.69 | 22.66 | 8462.138 | [-88.5, 31.12] | -1.27 | 2.5e-01 |
| **P1** | **MINDBLANK - NORECALL** | **-78.45** | **23.59** | **8456.822** | **[-140.71, -16.19]** | **-3.33** | **1.8e-03** |
| P1 | SLEPPY - ALERT | 1.25 | 12.45 | 8423.966 | [-23.15, 25.66] | 0.10 | 9.2e-01 |
| P1 | SLEPPY: ONTASK - MINDWANDER | 1.51 | 7.85 | 8343.801 | [-19.21, 22.23] | 0.19 | 8.6e-01 |
| P1 | SLEPPY: ONTASK - MINDBLANK | 23.64 | 9.96 | 8323.924 | [-2.64, 49.93] | 2.37 | 5.3e-02 |
| P1 | SLEPPY: ONTASK - NORECALL | -2.62 | 15.25 | 8445.705 | [-42.87, 37.63] | -0.17 | 8.6e-01 |
| P1 | SLEPPY: MINDWANDER - MINDBLANK | 22.14 | 8.70 | 8409.886 | [-0.82, 45.09] | 2.54 | 5.3e-02 |
| P1 | SLEPPY: MINDWANDER - NORECALL | -4.12 | 14.40 | 8467.775 | [-42.13, 33.89] | -0.29 | 8.6e-01 |
| P1 | SLEPPY: MINDBLANK - NORECALL | -26.26 | 15.36 | 8467.311 | [-66.79, 14.26] | -1.71 | 1.7e-01 |
| P1 | ALERT: ONTASK - MINDWANDER | 12.72 | 6.06 | 8339.332 | [-3.26, 28.71] | 2.10 | 5.4e-02 |
| **P1** | **ALERT: ONTASK - MINDBLANK** | **90.11** | **14.21** | **8438.600** | **[52.62, 127.61]** | **6.34** | **1.4e-09** |
| P1 | ALERT: ONTASK - NORECALL | -40.52 | 42.77 | 8456.762 | [-153.38, 72.33] | -0.95 | 3.4e-01 |
| **P1** | **ALERT: MINDWANDER - MINDBLANK** | **77.39** | **14.78** | **8418.875** | **[38.39, 116.39]** | **5.24** | **5.0e-07** |
| P1 | ALERT: MINDWANDER - NORECALL | -53.25 | 42.90 | 8456.295 | [-166.45, 59.95] | -1.24 | 2.6e-01 |
| **P1** | **ALERT: MINDBLANK - NORECALL** | **-130.64** | **44.55** | **8452.284** | **[-248.21, -13.07]** | **-2.93** | **6.7e-03** |
| **P2** | **ONTASK - MINDWANDER** | **11.18** | **4.56** | **7499.181** | **[-0.86, 23.22]** | **2.45** | **4.3e-02** |
| **P2** | **ONTASK - MINDBLANK** | **28.69** | **7.81** | **8160.428** | **[8.07, 49.31]** | **3.67** | **1.5e-03** |
| P2 | ONTASK - NORECALL | -0.47 | 20.44 | 8469.985 | [-54.4, 53.46] | -0.02 | 9.8e-01 |
| **P2** | **MINDWANDER - MINDBLANK** | **17.51** | **7.70** | **8215.360** | **[-2.81, 37.84]** | **2.27** | **4.6e-02** |
| P2 | MINDWANDER - NORECALL | -11.64 | 20.37 | 8469.385 | [-65.39, 42.1] | -0.57 | 6.8e-01 |
| P2 | MINDBLANK - NORECALL | -29.16 | 21.20 | 8465.509 | [-85.11, 26.8] | -1.38 | 2.5e-01 |
| P2 | SLEPPY - ALERT | 3.14 | 11.17 | 8274.753 | [-18.75, 25.04] | 0.28 | 7.8e-01 |
| **P2** | **SLEPPY: ONTASK - MINDWANDER** | **18.61** | **7.04** | **8029.365** | **[0.04, 37.17]** | **2.64** | **3.5e-02** |
| **P2** | **SLEPPY: ONTASK - MINDBLANK** | **22.55** | **8.92** | **7974.942** | **[-1, 46.1]** | **2.53** | **3.5e-02** |
| P2 | SLEPPY: ONTASK - NORECALL | -7.27 | 13.69 | 8342.365 | [-43.38, 28.85] | -0.53 | 6.1e-01 |
| P2 | SLEPPY: MINDWANDER - MINDBLANK | 3.94 | 7.80 | 8216.932 | [-16.64, 24.53] | 0.51 | 6.1e-01 |
| P2 | SLEPPY: MINDWANDER - NORECALL | -25.87 | 12.93 | 8434.095 | [-60, 8.25] | -2.00 | 6.8e-02 |
| P2 | SLEPPY: MINDBLANK - NORECALL | -29.82 | 13.80 | 8468.990 | [-66.22, 6.59] | -2.16 | 6.1e-02 |
| P2 | ALERT: ONTASK - MINDWANDER | 3.75 | 5.43 | 8008.392 | [-10.58, 18.07] | 0.69 | 7.4e-01 |
| **P2** | **ALERT: ONTASK - MINDBLANK** | **34.83** | **12.75** | **8308.365** | **[1.19, 68.47]** | **2.73** | **3.8e-02** |
| P2 | ALERT: ONTASK - NORECALL | 6.33 | 38.44 | 8465.524 | [-95.1, 107.76] | 0.16 | 9.5e-01 |
| P2 | ALERT: MINDWANDER - MINDBLANK | 31.09 | 13.25 | 8241.094 | [-3.89, 66.06] | 2.35 | 5.7e-02 |
| P2 | ALERT: MINDWANDER - NORECALL | 2.58 | 38.55 | 8465.064 | [-99.16, 104.32] | 0.07 | 9.5e-01 |
| P2 | ALERT: MINDBLANK - NORECALL | -28.50 | 40.05 | 8460.764 | [-134.18, 77.17] | -0.71 | 7.4e-01 |
| P3 | ONTASK - MINDWANDER | 8.89 | 4.70 | 7931.434 | [-3.52, 21.29] | 1.89 | 1.2e-01 |
| **P3** | **ONTASK - MINDBLANK** | **38.46** | **8.04** | **8322.384** | **[17.25, 59.67]** | **4.79** | **1.0e-05** |
| P3 | ONTASK - NORECALL | 18.26 | 21.00 | 8468.168 | [-37.15, 73.67] | 0.87 | 4.6e-01 |
| **P3** | **MINDWANDER - MINDBLANK** | **29.57** | **7.92** | **8353.350** | **[8.67, 50.48]** | **3.73** | **5.7e-04** |
| P3 | MINDWANDER - NORECALL | 9.37 | 20.92 | 8465.774 | [-45.84, 64.59] | 0.45 | 6.5e-01 |
| P3 | MINDBLANK - NORECALL | -20.20 | 21.78 | 8460.572 | [-77.68, 37.28] | -0.93 | 4.6e-01 |
| P3 | SLEPPY - ALERT | 19.26 | 11.49 | 8379.395 | [-3.25, 41.78] | 1.68 | 9.4e-02 |
| P3 | **SLEPPY: ONTASK - MINDWANDER** | **2.76** | **7.24** | **8246.471** | **[-16.35, 21.87]** | **0.38** | **7.0e-01** |
| P3 | SLEPPY: ONTASK - MINDBLANK | 33.00 | 9.18 | 8215.041 | [8.76, 57.23] | 3.59 | 9.9e-04 |
| P3 | SLEPPY: ONTASK - NORECALL | -7.76 | 14.07 | 8416.266 | [-44.9, 29.38] | -0.55 | 7.0e-01 |
| P3 | **SLEPPY: MINDWANDER - MINDBLANK** | **30.24** | **8.02** | **8352.907** | **[9.07, 51.41]** | **3.77** | **9.9e-04** |
| P3 | SLEPPY: MINDWANDER - NORECALL | -10.52 | 13.29 | 8459.917 | [-45.6, 24.56] | -0.79 | 6.4e-01 |
| P3 | **SLEPPY: MINDBLANK - NORECALL** | **-40.76** | **14.18** | **8469.575** | **[-78.17, -3.35]** | **-2.88** | **8.1e-03** |
| P3 | **ALERT: ONTASK - MINDWANDER** | **15.01** | **5.59** | **8237.526** | **[0.27, 29.75]** | **2.69** | **2.2e-02** |
| P3 | **ALERT: ONTASK - MINDBLANK** | **43.92** | **13.11** | **8401.855** | **[9.33, 78.51]** | **3.35** | **4.9e-03** |
| P3 | ALERT: ONTASK - NORECALL | 44.28 | 39.48 | 8460.539 | [-59.91, 148.47] | 1.12 | 3.9e-01 |
| P3 | ALERT: MINDWANDER - MINDBLANK | 28.91 | 13.63 | 8367.278 | [-7.07, 64.89] | 2.12 | 6.8e-02 |
| P3 | ALERT: MINDWANDER - NORECALL | 29.27 | 39.60 | 8460.047 | [-75.25, 133.78] | 0.74 | 5.5e-01 |
| P3 | ALERT: MINDBLANK - NORECALL | 0.36 | 41.13 | 8455.734 | [-108.19, 108.9] | 0.01 | 9.9e-01 |
| P4 | ONTASK - MINDWANDER | 7.72 | 4.52 | 8036.332 | [-4.2, 19.64] | 1.71 | 1.9e-01 |
| P4 | **ONTASK - MINDBLANK** | **20.51** | **7.72** | **8358.083** | **[0.13, 40.9]** | **2.66** | **4.8e-02** |
| P4 | ONTASK - NORECALL | 5.61 | 20.17 | 8466.864 | [-47.61, 58.84] | 0.28 | 9.2e-01 |
| P4 | MINDWANDER - MINDBLANK | 12.79 | 7.61 | 8383.067 | [-7.3, 32.88] | 1.68 | 1.9e-01 |
| P4 | MINDWANDER - NORECALL | -2.11 | 20.10 | 8464.173 | [-55.14, 50.93] | -0.10 | 9.2e-01 |
| P4 | MINDBLANK - NORECALL | -14.90 | 20.92 | 8458.854 | [-70.11, 40.31] | -0.71 | 7.1e-01 |
| P4 | SLEPPY - ALERT | -2.57 | 11.04 | 8402.438 | [-24.2, 19.07] | -0.23 | 8.2e-01 |
| P4 | SLEPPY: ONTASK - MINDWANDER | 6.87 | 6.96 | 8296.160 | [-11.49, 25.24] | 0.99 | 3.2e-01 |
| P4 | **SLEPPY: ONTASK - MINDBLANK** | **21.38** | **8.83** | **8270.494** | **[-1.91, 44.67]** | **2.42** | **2.9e-02** |
| P4 | **SLEPPY: ONTASK - NORECALL** | **-31.58** | **13.52** | **8431.730** | **[-67.27, 4.1]** | **-2.34** | **2.9e-02** |
| P4 | **SLEPPY: MINDWANDER - MINDBLANK** | **14.50** | **7.71** | **8382.428** | **[-5.84, 34.85]** | **1.88** | **7.2e-02** |
| P4 | **SLEPPY: MINDWANDER - NORECALL** | **-38.46** | **12.77** | **8464.377** | **[-72.16, -4.76]** | **-3.01** | **7.8e-03** |
| P4 | **SLEPPY: MINDBLANK - NORECALL** | **-52.96** | **13.62** | **8468.737** | **[-88.9, -17.03]** | **-3.89** | **6.1e-04** |
| P4 | ALERT: ONTASK - MINDWANDER | 8.57 | 5.37 | 8289.589 | [-5.6, 22.74] | 1.60 | 3.6e-01 |
| P4 | ALERT: ONTASK - MINDBLANK | 19.65 | 12.60 | 8421.178 | [-13.59, 52.89] | 1.56 | 3.6e-01 |
| P4 | ALERT: ONTASK - NORECALL | 42.81 | 37.92 | 8458.808 | [-57.26, 142.89] | 1.13 | 4.8e-01 |
| P4 | ALERT: MINDWANDER - MINDBLANK | 11.08 | 13.10 | 8394.150 | [-23.49, 45.65] | 0.85 | 4.8e-01 |
| P4 | ALERT: MINDWANDER - NORECALL | 34.24 | 38.04 | 8458.324 | [-66.14, 134.63] | 0.90 | 4.8e-01 |
| P4 | ALERT: MINDBLANK - NORECALL | 23.16 | 39.51 | 8454.127 | [-81.1, 127.42] | 0.59 | 5.6e-01 |
| P4 | ONTASK - MINDWANDER | 7.72 | 4.52 | 8036.332 | [-4.2, 19.64] | 1.71 | 1.9e-01 |
| P5 | ONTASK - MINDWANDER | 4.24 | 6.52 | 8428.536 | [-12.96, 21.45] | 0.65 | 5.2e-01 |
| P5 | ONTASK - MINDBLANK | 20.19 | 11.13 | 8467.739 | [-9.17, 49.55] | 1.81 | 3.0e-01 |
| P5 | ONTASK - NORECALL | 41.32 | 28.99 | 8452.907 | [-35.18, 117.83] | 1.43 | 3.0e-01 |
| P5 | MINDWANDER - MINDBLANK | 15.95 | 10.96 | 8469.229 | [-12.99, 44.88] | 1.45 | 3.0e-01 |
| P5 | MINDWANDER - NORECALL | 37.08 | 28.88 | 8450.349 | [-39.14, 113.3] | 1.28 | 3.0e-01 |
| P5 | MINDBLANK - NORECALL | 21.13 | 30.06 | 8446.493 | [-58.2, 100.47] | 0.70 | 5.2e-01 |
| P5 | SLEPPY - ALERT | 7.13 | 15.89 | 8469.685 | [-24.02, 38.28] | 0.45 | 6.5e-01 |
| P5 | SLEPPY: ONTASK - MINDWANDER | 5.03 | 10.03 | 8462.177 | [-21.44, 31.49] | 0.50 | 7.5e-01 |
| P5 | SLEPPY: ONTASK - MINDBLANK | -0.46 | 12.72 | 8459.465 | [-34.03, 33.12] | -0.04 | 9.7e-01 |
| P5 | SLEPPY: ONTASK - NORECALL | -23.16 | 19.46 | 8469.795 | [-74.52, 28.2] | -1.19 | 4.9e-01 |
| P5 | SLEPPY: MINDWANDER - MINDBLANK | -5.49 | 11.10 | 8469.127 | [-34.79, 23.82] | -0.49 | 7.5e-01 |
| P5 | SLEPPY: MINDWANDER - NORECALL | -28.19 | 18.37 | 8465.006 | [-76.66, 20.29] | -1.53 | 4.9e-01 |
| P5 | SLEPPY: MINDBLANK - NORECALL | -22.70 | 19.58 | 8455.376 | [-74.36, 28.96] | -1.16 | 4.9e-01 |
| P5 | ALERT: ONTASK - MINDWANDER | 3.46 | 7.74 | 8461.929 | [-16.96, 23.88] | 0.45 | 6.5e-01 |
| P5 | ALERT: ONTASK - MINDBLANK | 40.84 | 18.13 | 8469.958 | [-7.01, 88.69] | 2.25 | 9.2e-02 |
| P5 | ALERT: ONTASK - NORECALL | 105.81 | 54.49 | 8446.408 | [-38, 249.61] | 1.94 | 9.2e-02 |
| P5 | ALERT: MINDWANDER - MINDBLANK | 37.38 | 18.86 | 8469.660 | [-12.4, 87.16] | 1.98 | 9.2e-02 |
| P5 | ALERT: MINDWANDER - NORECALL | 102.35 | 54.66 | 8446.106 | [-41.9, 246.6] | 1.87 | 9.2e-02 |
| P5 | ALERT: MINDBLANK - NORECALL | 64.97 | 56.77 | 8443.542 | [-84.83, 214.77] | 1.14 | 3.0e-01 |
| SE = standard error; CL = confidence limits 95 %; FDR = false discovery rate; D.F. = degrees of freedom | | | | | | | |

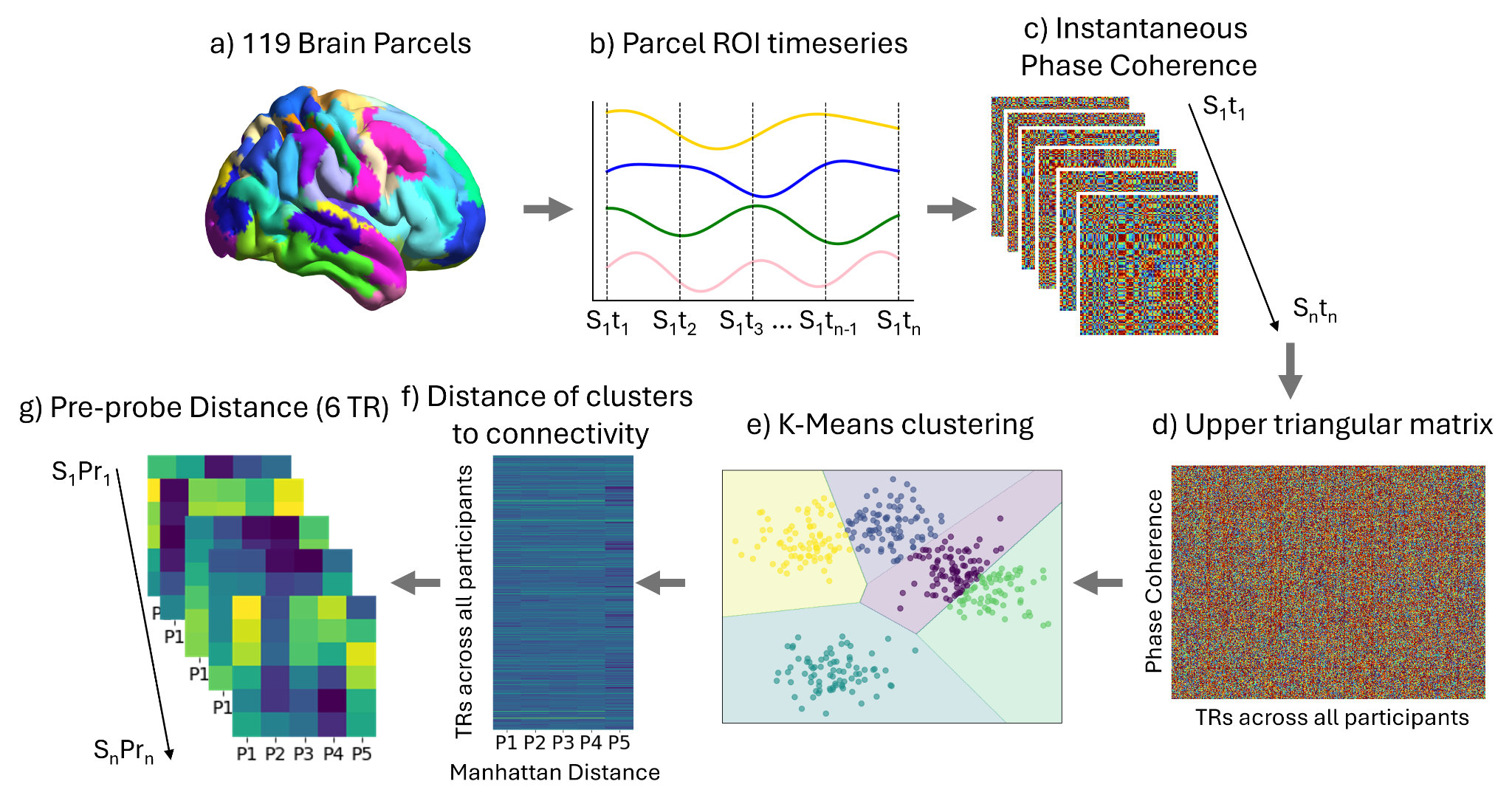

**Supplementary Figure S1. Schematic representation of time-varying functional connectivity clustering.** a) Nineteen subcortical parcels were appended to the Schaefer 100 cortical regions of interest (ROIs), yielding a combined cortical–subcortical parcellation. b) For each subject S, the mean BOLD timeseries was extracted from every ROI, resulting in an N_TR_ x 119 matrix, where each row corresponds to the mean BOLD signal per ROI. c) The Hilbert transform was applied to obtain an analytic representation of each ROI timeseries, from which instantaneous phase coherence was computed between all ROI pairs at every BOLD acquisition. d) Given that phase coherence matrices are symmetric, only the upper triangular elements were retained for each subject and timepoint. These matrices were concatenated across subjects and time to generate a comprehensive dataset representation of phase coherence. e) To identify representative connectivity patterns that capture the temporal organization of functional connectivity, we applied K-means clustering using the Manhattan distance metric. This procedure yielded five recurring connectivity patterns that describe the observed functional connectivity across time. f) For each timepoint, we quantified the similarity to each of the five connectivity patterns by computing the Manhattan distance. Effectively, this informed us as to how similar the brain looks to each of the connectivity patterns at every TR. g) Finally, to investigate probe-specific functional connectivity, we extracted the last six TRs (approximately 10 seconds) preceding each probe P, capturing their distances to the identified connectivity patterns. The y-axis of each heatmap represents distances from t_n-6_ up to t_n-1_ BOLD scans before each probe. Each participant had approximately 40 probes.

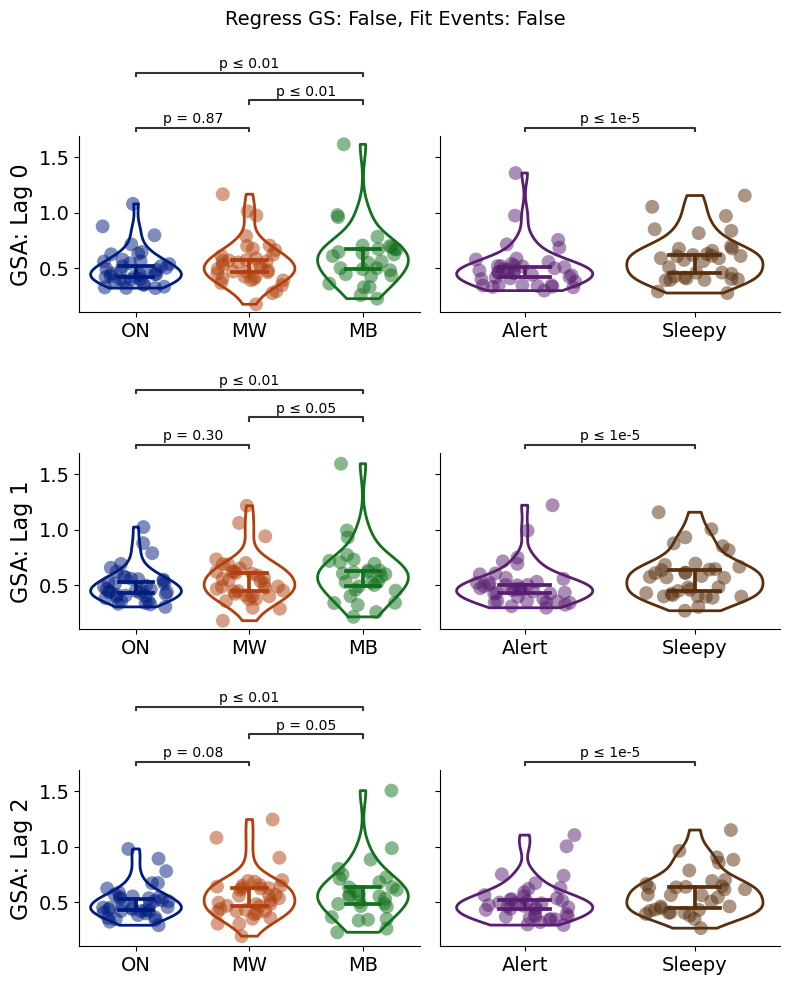

**Supplementary Figure S2.** Replication analysis of the relationship between global signal amplitude, mental states and alertness levels. Analysis parameters: task regression = False, Global signal regression = False. Across 3 different lags (shifting the analysis windows by 1 TR relative to the auditory probe to report content and alertness), participant consistently indicate higher global signal amplitude during MB and sleepiness. GSA = global signal amplitude

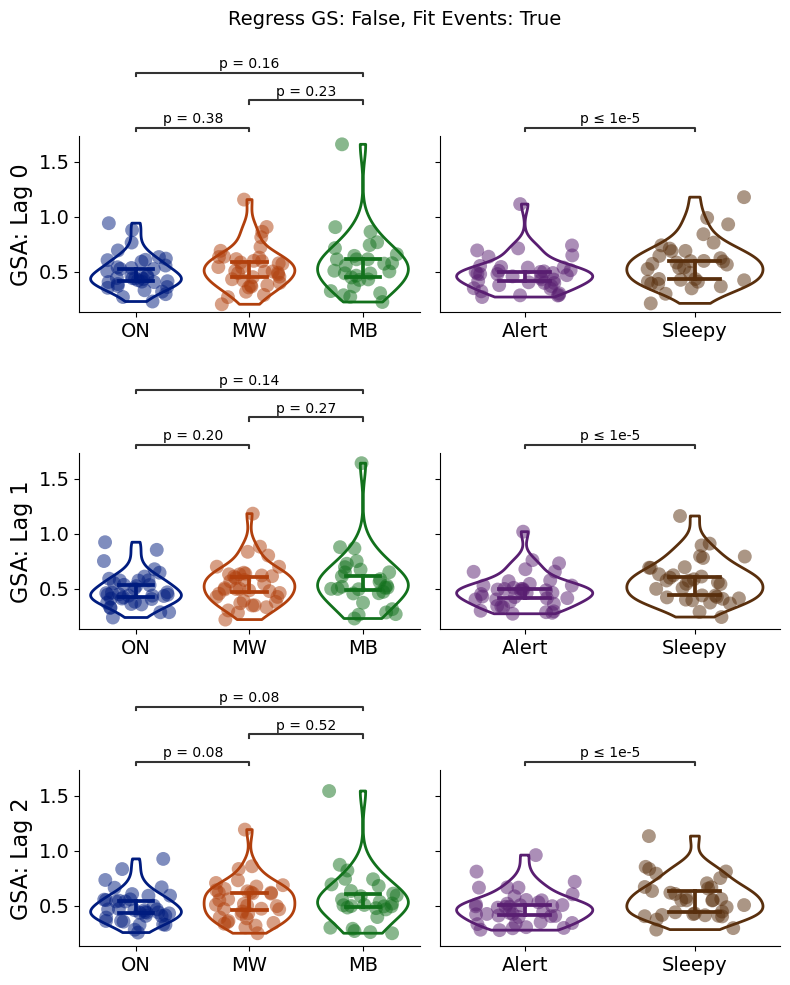

**Supplementary Figure S3.** Replication analysis of the relationship between global signal amplitude, mental states and alertness levels. Analysis parameters: task regression = True, Global signal regression = False. Across 3 different lags (shifting the analysis windows by 1 TR relative to the auditory probe to report content and alertness), participant consistently indicate higher global signal amplitude sleepiness, but not MB. GSA = global signal amplitude

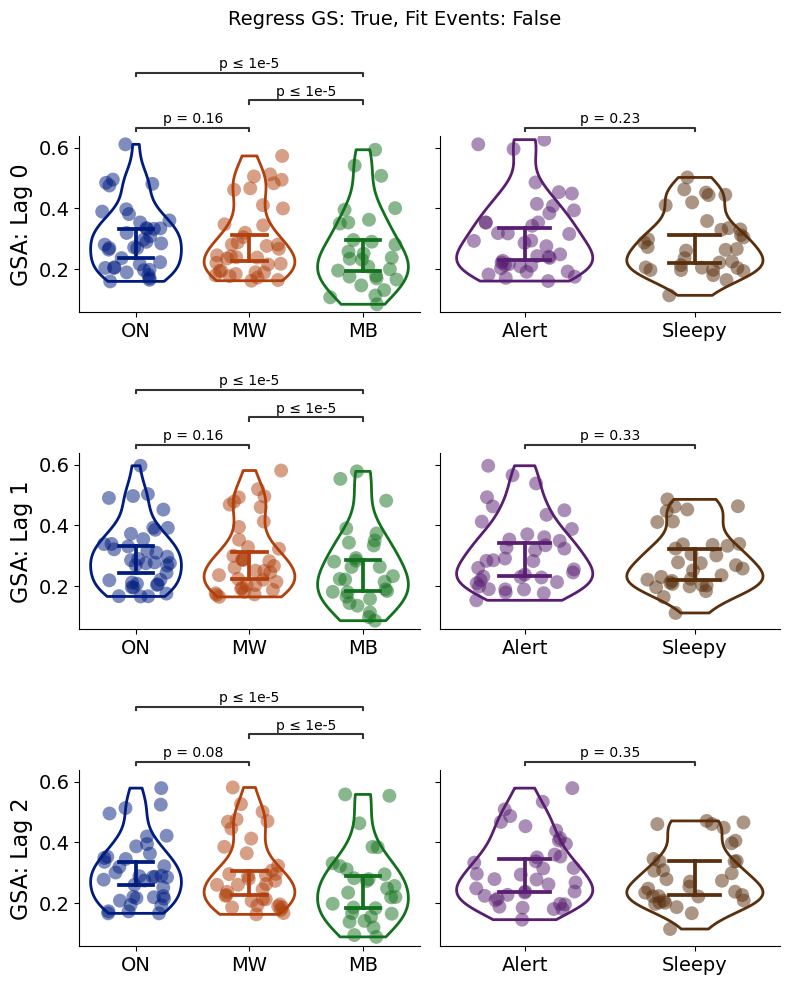

**Supplementary Figure S4.** Replication analysis of the relationship between global signal amplitude, mental states and alertness levels. Analysis parameters: task regression = False, Global signal regression = True, N ROI = 100. Global signal regression removed the relationship between MB and sleepiness and global signal amplitude. GSA = global signal amplitude

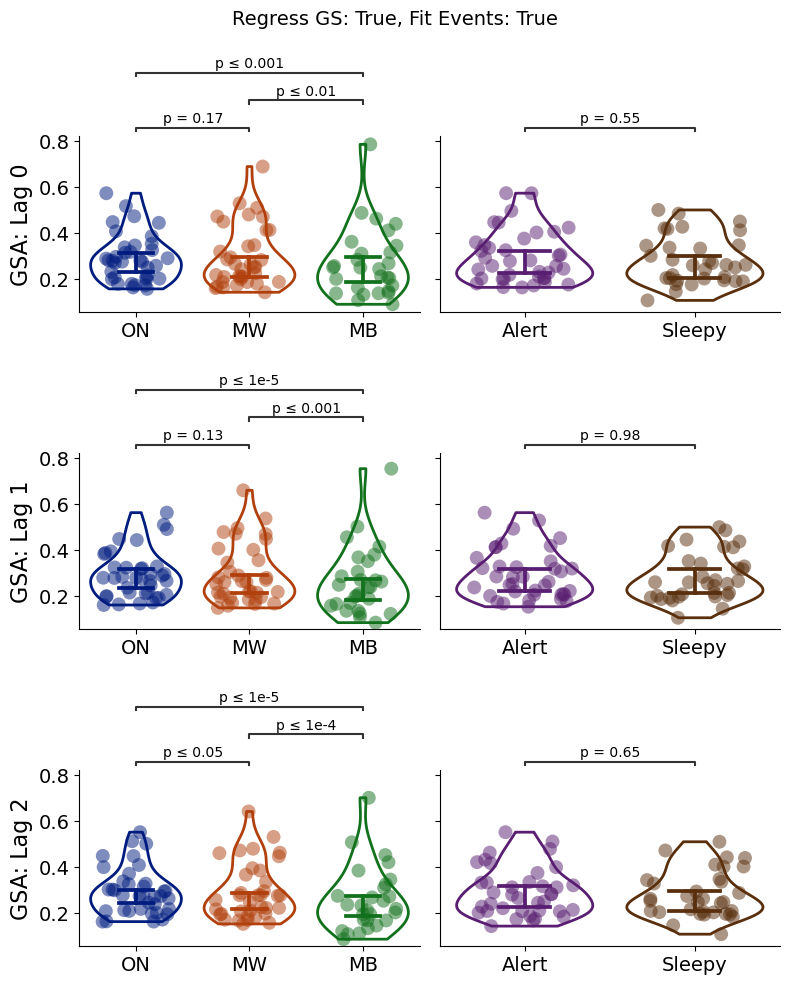

**Supplementary Figure S5.** Replication analysis of the relationship between global signal amplitude, mental states and alertness levels. Analysis parameters: task regression = True, Global signal regression = True, N ROI = 100. Global signal regression removed the relationship between MB and sleepiness and global signal amplitude. GSA = global signal amplitude

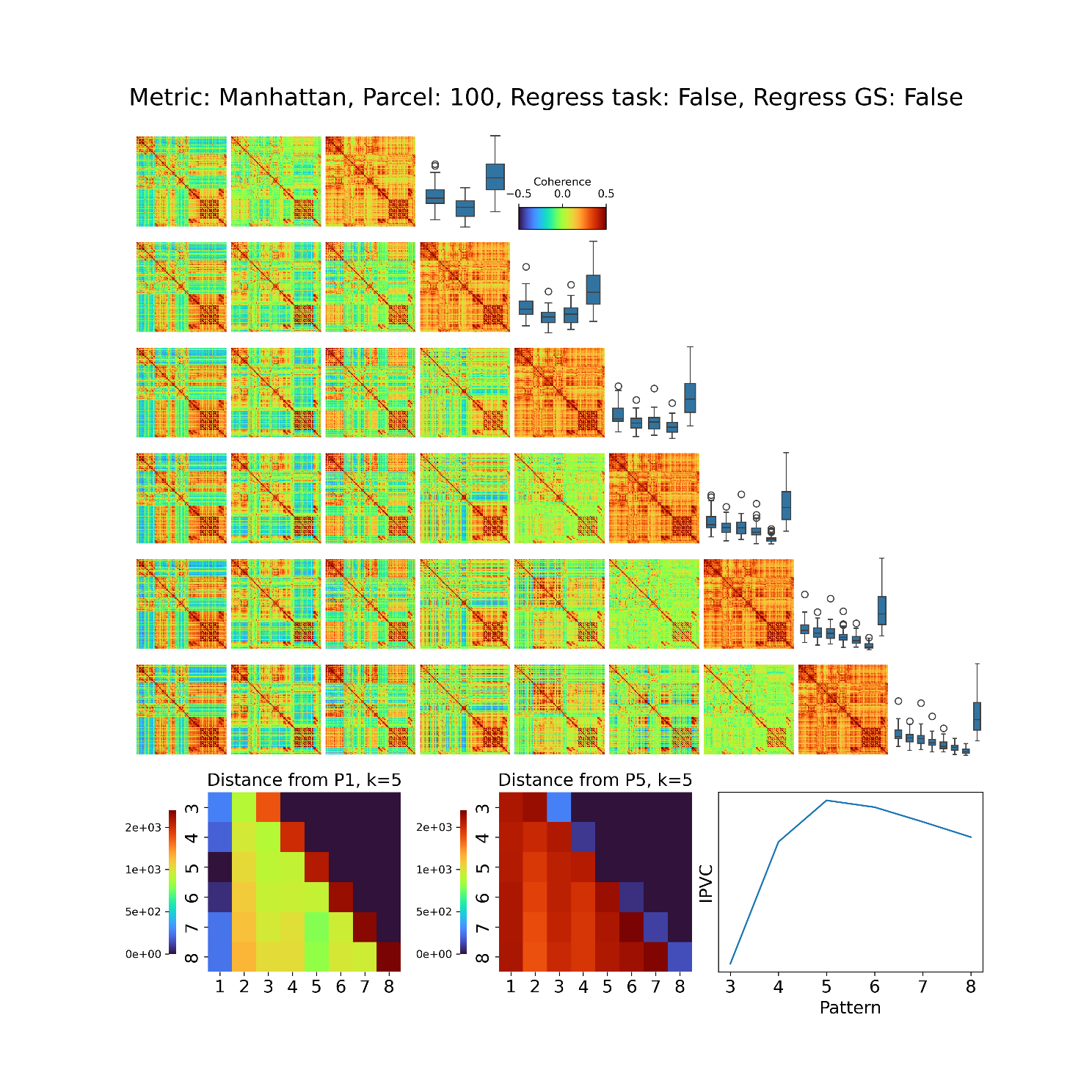

**Supplementary Figure S6.** Recurrent and consistent brain configurations emerge under different clustering dimensionalities during task engagement. Clustering parameters: Metric = Manhattan distance, Global Signal Regression: False, Event Regression: False. *Heatmaps:* For each value of *k*, patterns are ordered based on the standard deviation of their connectome, from the most variant (left) to the least (right). *Bottom row – Left*: To examine whether patterns produced across different values of *k* presented correspondence, we estimated the distance of pattern 1 for *k*= 5 with respect to all patterns obtained for *k* = 3 to 8. Similarly, we estimated the distance of pattern 5 for *k*= 5 with respect to all patterns obtained for *k* = 3 to 8. *Bottom row – Right*: For each selection of the number of clusters in the K-means algorithm, we computed the inter-pattern correlation variability (IPVC) between all the upper triangular parts of the resulting centroids**.** The variability of dynamic coordination patterns found by the clustering procedure is maximal with *k* = 5 clusters.

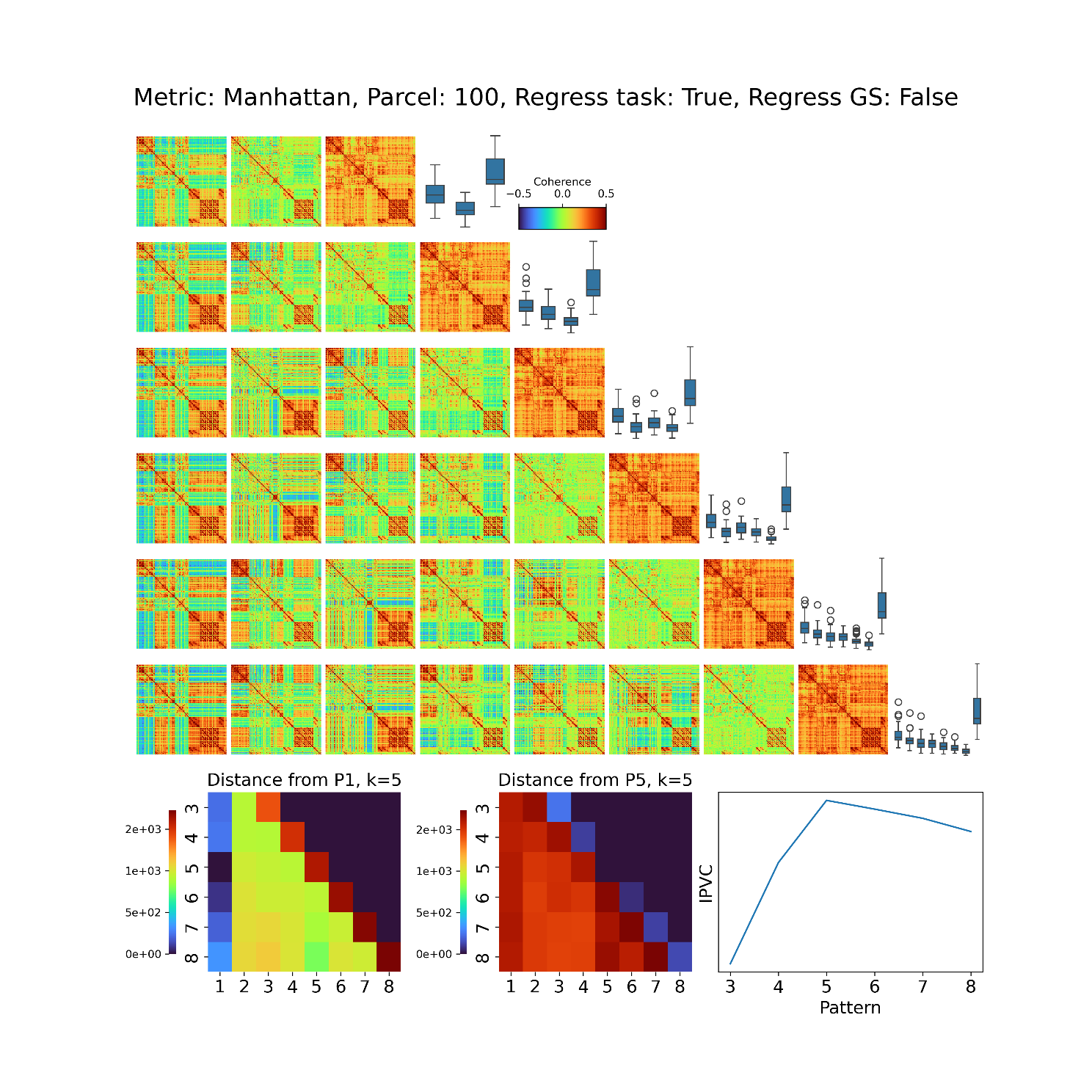

**Supplementary Figure S7.** Recurrent and consistent brain configurations emerge under different clustering dimensionalities during task engagement. Clustering parameters: Metric = Manhattan distance, Global Signal Regression: False, Event Regression: True. *Heatmaps:* For each value of *k*, patterns are ordered based on the standard deviation of their connectome, from the most variant (left) to the least (right). *Bottom row – Left*: To examine whether patterns produced across different values of *k* presented correspondence, we estimated the distance of pattern 1 for *k*= 5 with respect to all patterns obtained for *k* = 3 to 8. Similarly, we estimated the distance of pattern 5 for *k*= 5 with respect to all patterns obtained for *k* = 3 to 8. *Bottom row – Right*: The variability of dynamic coordination patterns found by the clustering procedure is maximal with *k* = 5 clusters. For each selection of the number of clusters in the K-means algorithm, we computed the inter-pattern correlation variability (IPVC) between all the upper triangular parts of the resulting centroids**.**

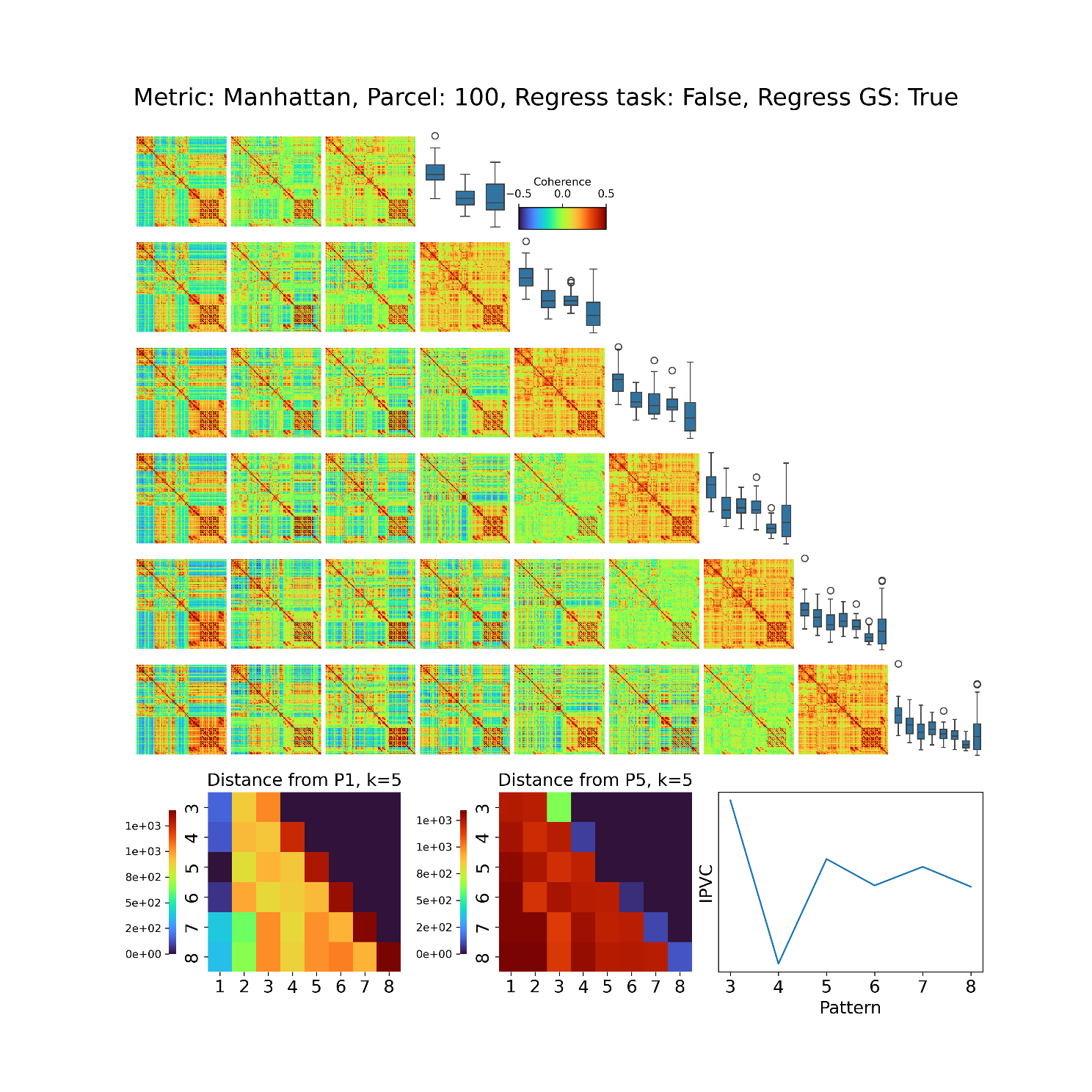

**Supplementary Figure S8.** Recurrent and consistent brain configurations emerge under different clustering dimensionalities during task engagement. Clustering parameters: Metric = Manhattan distance, Global Signal Regression: True, Event Regression: False. *Heatmaps:* For each value of *k*, patterns are ordered based on the standard deviation of their connectome, from the most variant (left) to the least (right). *Bottom row – Left*: To examine whether patterns produced across different values of *k* presented correspondence, we estimated the distance of pattern 1 for *k*= 5 with respect to all patterns obtained for *k* = 3 to 8. Similarly, we estimated the distance of pattern 5 for *k*= 5 with respect to all patterns obtained for *k* = 3 to 8. *Bottom row – Right*: The variability of dynamic coordination patterns found by the clustering procedure is maximal with *k* = 5 clusters. For each selection of the number of clusters in the K-means algorithm, we computed the inter-pattern correlation variability (IPVC) between all the upper triangular parts of the resulting centroids**.**

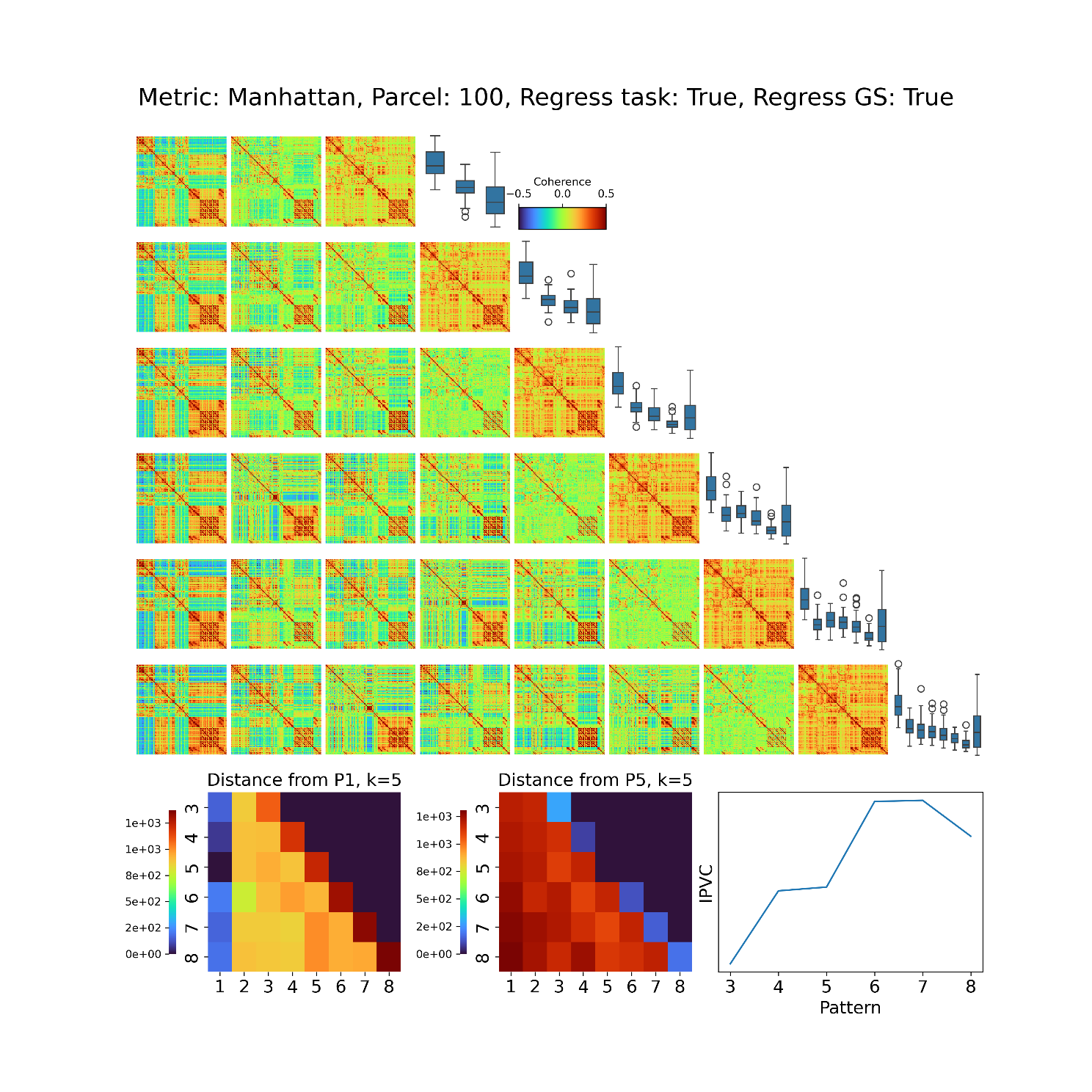

**Supplementary Figure S9.** Recurrent and consistent brain configurations emerge under different clustering dimensionalities during task engagement. Clustering parameters: Metric = Manhattan distance, Global Signal Regression: True, Event Regression: True. *Heatmaps:* For each value of *k*, patterns are ordered based on the standard deviation of their connectome, from the most variant (left) to the least (right). *Bottom row – Left*: To examine whether patterns produced across different values of *k* presented correspondence, we estimated the distance of pattern 1 for *k*= 5 with respect to all patterns obtained for *k* = 3 to 8. Similarly, we estimated the distance of pattern 5 for *k*= 5 with respect to all patterns obtained for *k* = 3 to 8. *Bottom row – Right*: The variability of dynamic coordination patterns found by the clustering procedure is maximal with *k* = 5 clusters. For each selection of the number of clusters in the K-means algorithm, we computed the inter-pattern correlation variability (IPVC) between all the upper triangular parts of the resulting centroids**.**

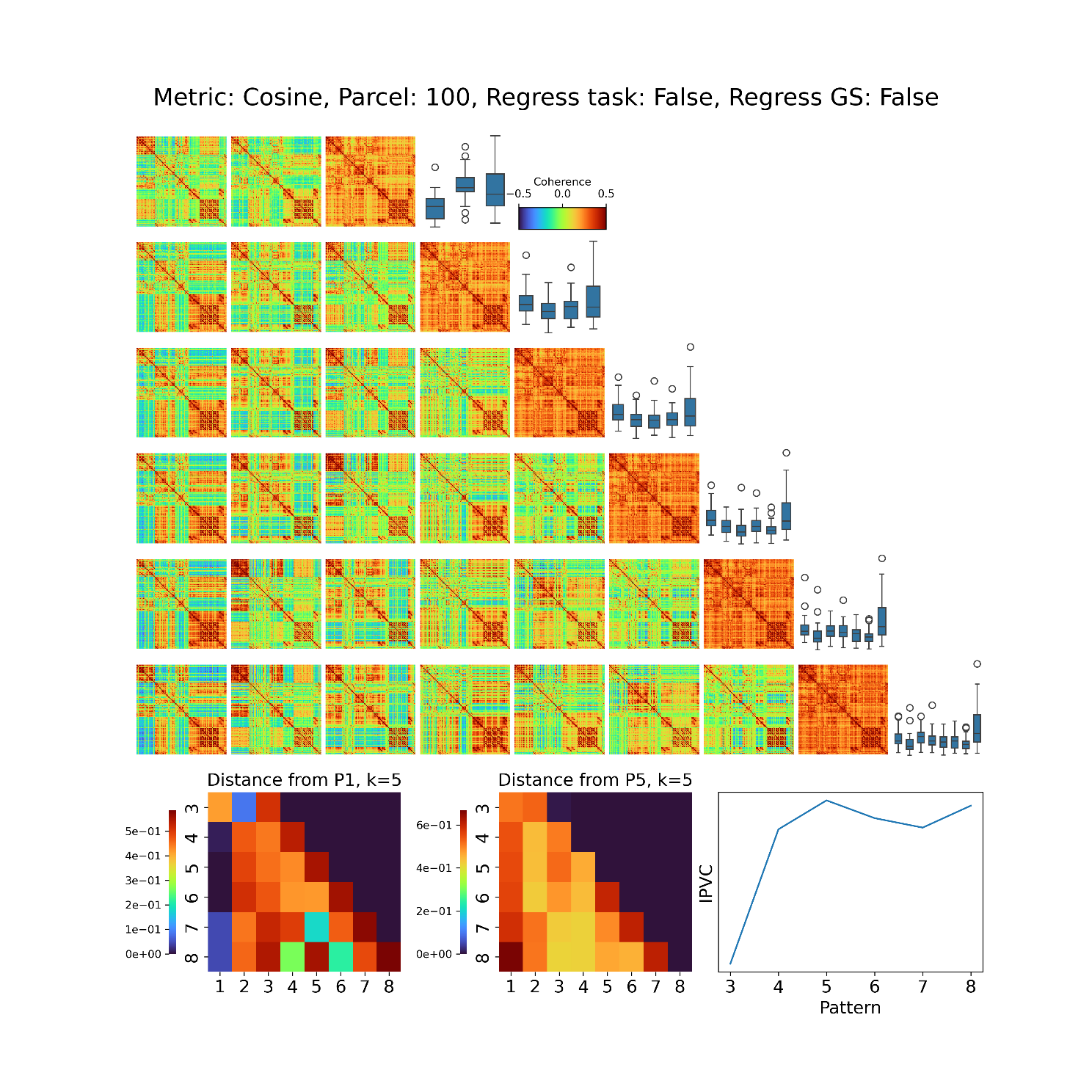

**Supplementary Figure S10.** Recurrent and consistent brain configurations emerge under different clustering dimensionalities during task engagement. Clustering parameters: Metric = **Cosine** distance, Global Signal Regression: False, Event Regression: False. *Heatmaps:* For each value of *k*, patterns are ordered based on the standard deviation of their connectome, from the most variant (left) to the least (right). *Bottom row – Left*: To examine whether patterns produced across different values of *k* presented correspondence, we estimated the distance of pattern 1 for *k*= 5 with respect to all patterns obtained for *k* = 3 to 8. Similarly, we estimated the distance of pattern 5 for *k*= 5 with respect to all patterns obtained for *k* = 3 to 8. *Bottom row – Right*: The variability of dynamic coordination patterns found by the clustering procedure is maximal with *k* = 5 clusters. For each selection of the number of clusters in the K-means algorithm, we computed the inter-pattern correlation variability (IPVC) between all the upper triangular parts of the resulting centroids**.**

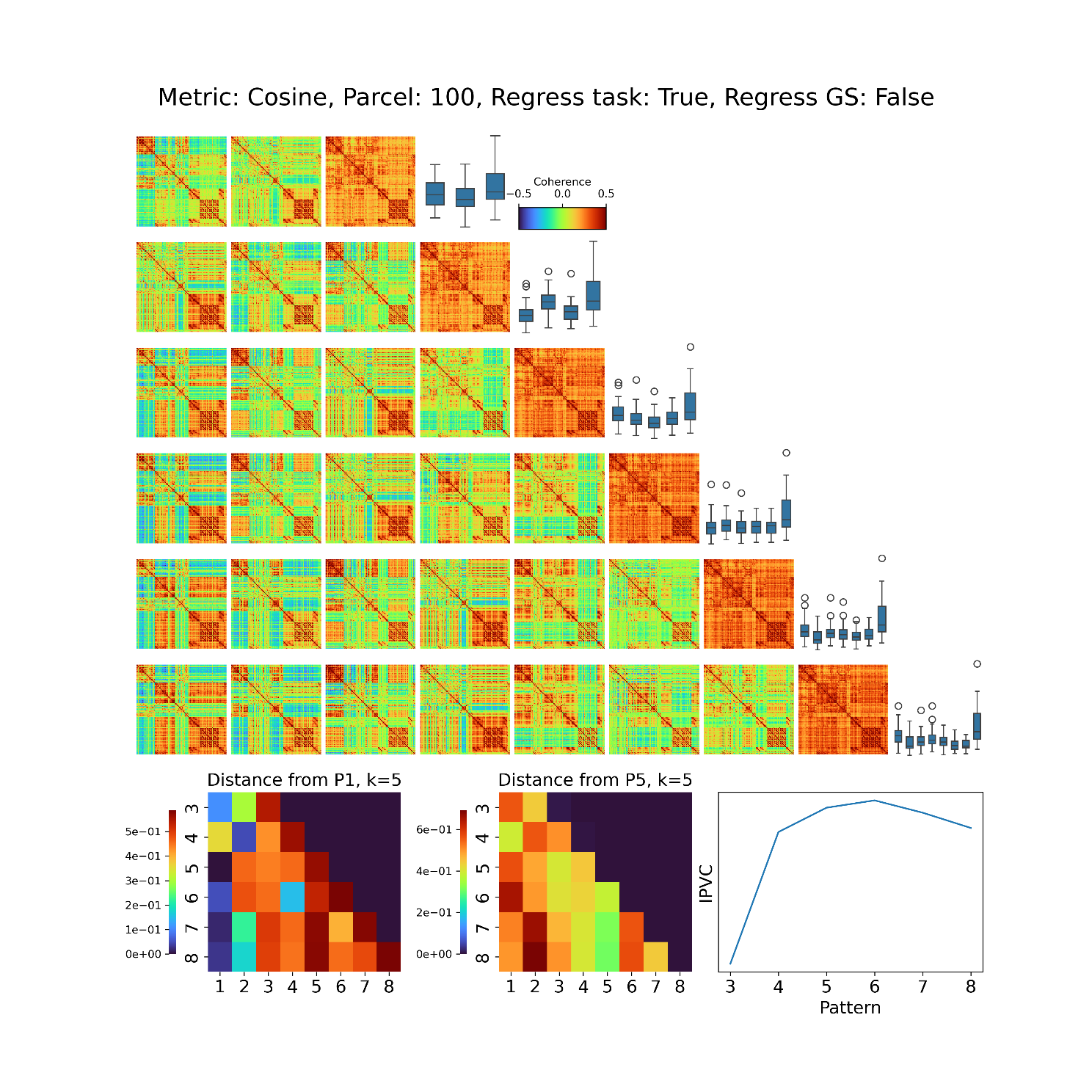

**Supplementary Figure S11.** Recurrent and consistent brain configurations emerge under different clustering dimensionalities during task engagement. Clustering parameters: Metric = **Cosine** distance, Global Signal Regression: False, Event Regression: True. *Heatmaps:* For each value of *k*, patterns are ordered based on the standard deviation of their connectome, from the most variant (left) to the least (right). *Bottom row – Left*: To examine whether patterns produced across different values of *k* presented correspondence, we estimated the distance of pattern 1 for *k*= 5 with respect to all patterns obtained for *k* = 3 to 8. Similarly, we estimated the distance of pattern 5 for *k*= 5 with respect to all patterns obtained for *k* = 3 to 8. *Bottom row – Right*: The variability of dynamic coordination patterns found by the clustering procedure is maximal with *k* = 5 clusters. For each selection of the number of clusters in the K-means algorithm, we computed the inter-pattern correlation variability (IPVC) between all the upper triangular parts of the resulting centroids**.**

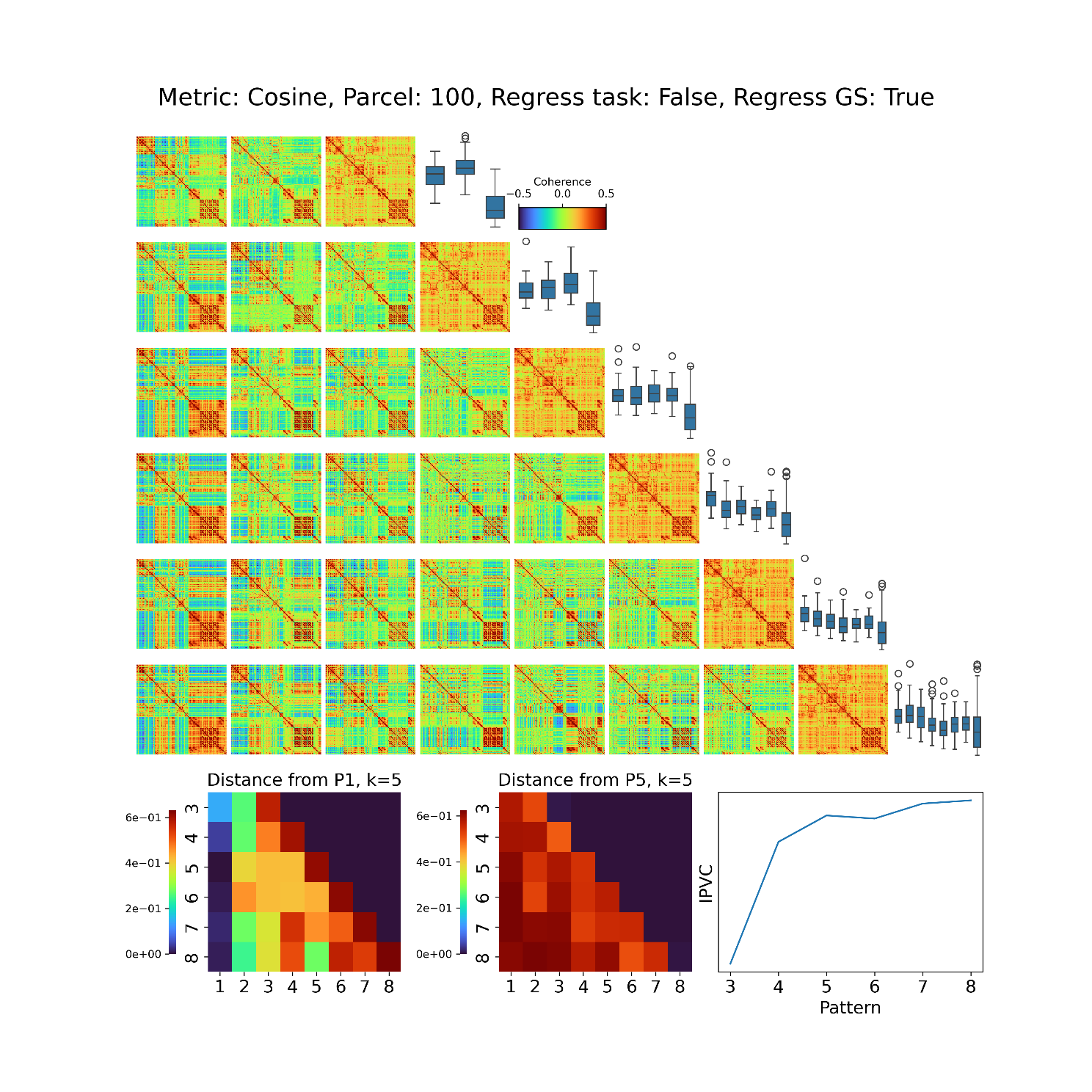

**Supplementary Figure S12.** Recurrent and consistent brain configurations emerge under different clustering dimensionalities during task engagement. Clustering parameters: Metric = **Cosine** distance, Global Signal Regression: True, Event Regression: False. *Heatmaps:* For each value of *k*, patterns are ordered based on the standard deviation of their connectome, from the most variant (left) to the least (right). *Bottom row – Left*: To examine whether patterns produced across different values of *k* presented correspondence, we estimated the distance of pattern 1 for *k*= 5 with respect to all patterns obtained for *k* = 3 to 8. Similarly, we estimated the distance of pattern 5 for *k*= 5 with respect to all patterns obtained for *k* = 3 to 8. *Bottom row – Right*: The variability of dynamic coordination patterns found by the clustering procedure is maximal with *k* = 5 clusters. For each selection of the number of clusters in the K-means algorithm, we computed the inter-pattern correlation variability (IPVC) between all the upper triangular parts of the resulting centroids**.**

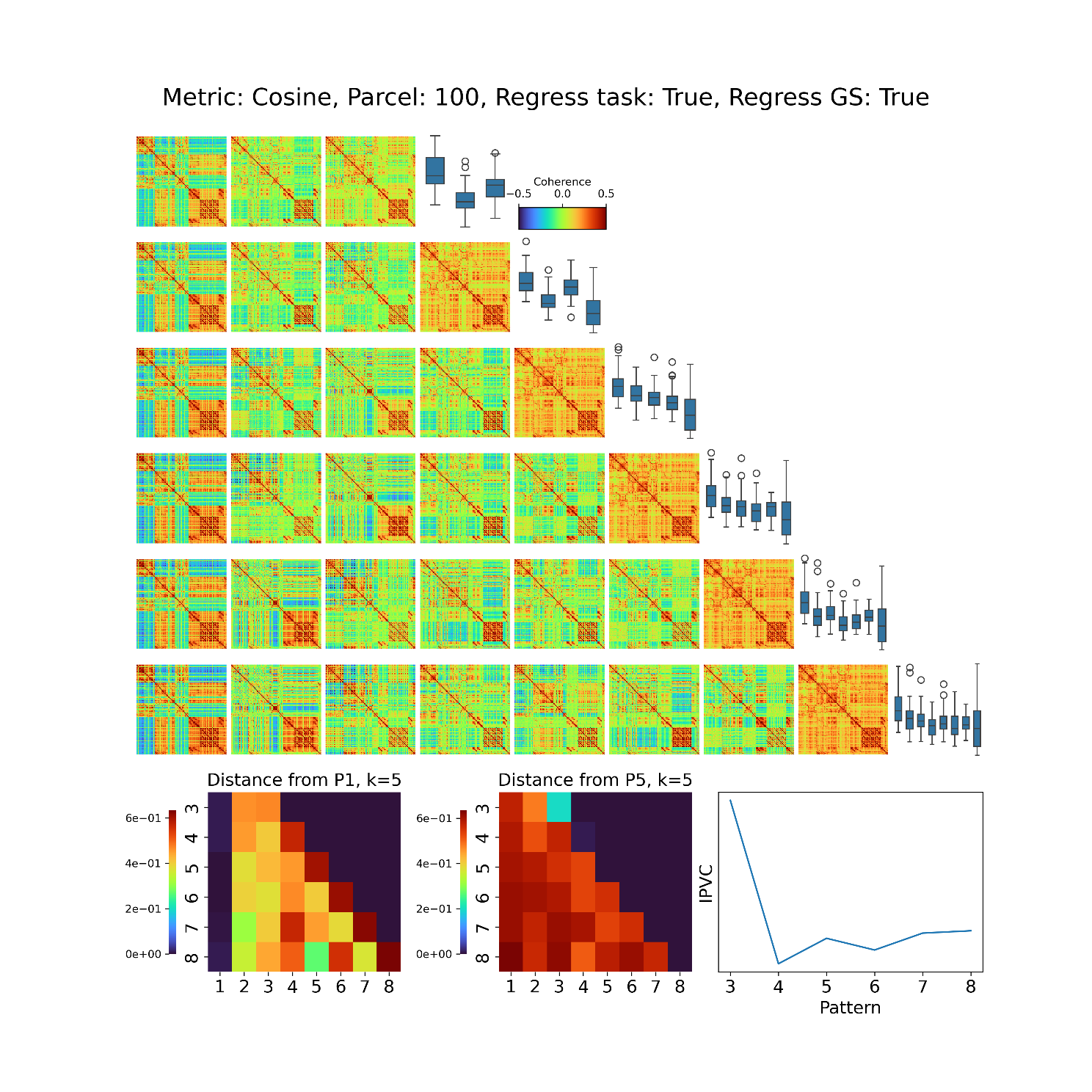

**Supplementary Figure S13.** Recurrent and consistent brain configurations emerge under different clustering dimensionalities during task engagement. Clustering parameters: Metric = **Cosine** distance, Global Signal Regression: True, Event Regression: True. *Heatmaps:* For each value of *k*, patterns are ordered based on the standard deviation of their connectome, from the most variant (left) to the least (right). *Bottom row – Left*: To examine whether patterns produced across different values of *k* presented correspondence, we estimated the distance of pattern 1 for *k*= 5 with respect to all patterns obtained for *k* = 3 to 8. Similarly, we estimated the distance of pattern 5 for *k*= 5 with respect to all patterns obtained for *k* = 3 to 8. *Bottom row – Right*: The variability of dynamic coordination patterns found by the clustering procedure is maximal with *k* = 5 clusters. For each selection of the number of clusters in the K-means algorithm, we computed the inter-pattern correlation variability (IPVC) between all the upper triangular parts of the resulting centroids**.**

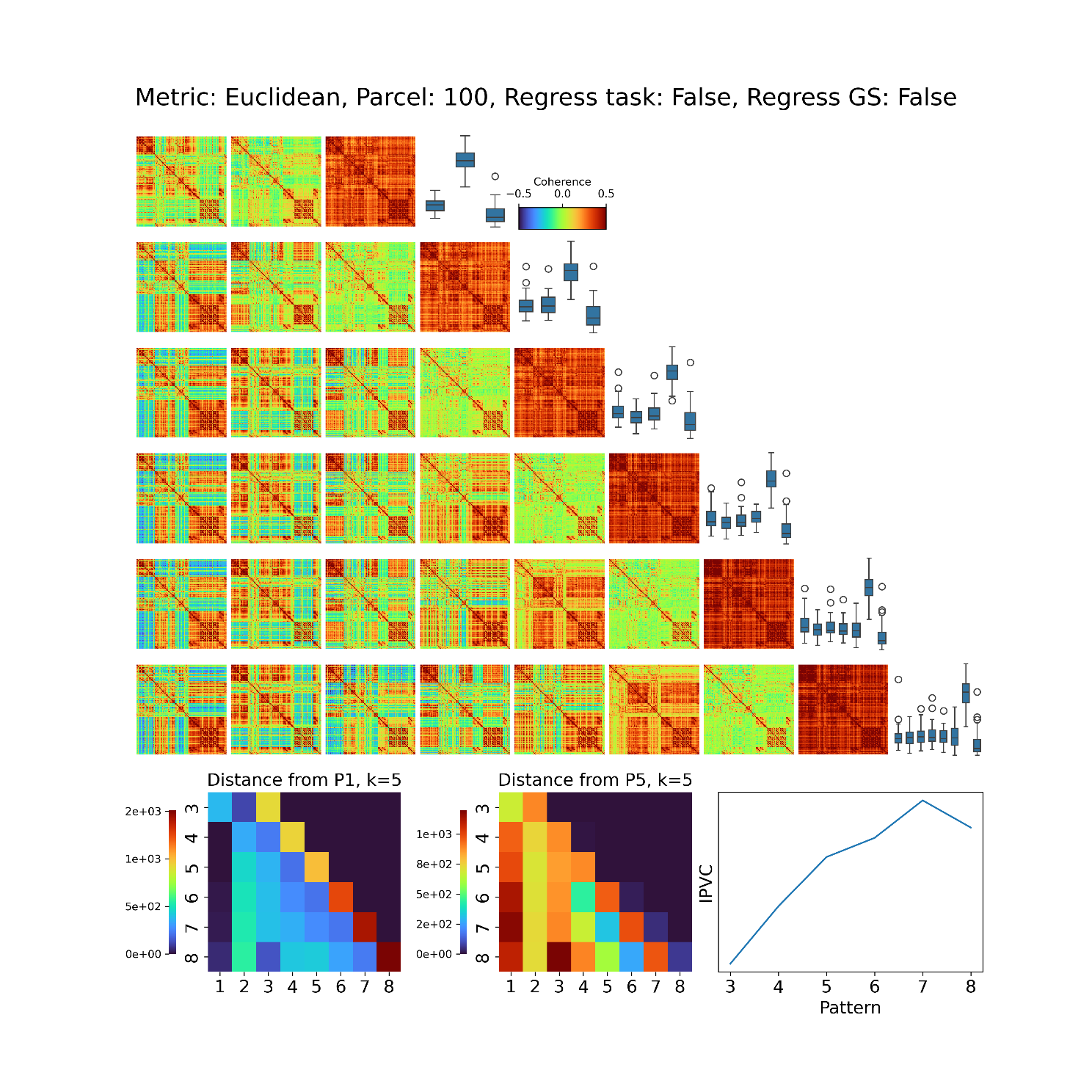

**Supplementary Figure S14.** Recurrent and consistent brain configurations emerge under different clustering dimensionalities during task engagement. Clustering parameters: Metric = **Euclidean** distance, Global Signal Regression: False, Event Regression: False. *Heatmaps:* For each value of *k*, patterns are ordered based on the standard deviation of their connectome, from the most variant (left) to the least (right). *Bottom row – Left*: To examine whether patterns produced across different values of *k* presented correspondence, we estimated the distance of pattern 1 for *k*= 5 with respect to all patterns obtained for *k* = 3 to 8. Similarly, we estimated the distance of pattern 5 for *k*= 5 with respect to all patterns obtained for *k* = 3 to 8. *Bottom row – Right*: The variability of dynamic coordination patterns found by the clustering procedure is maximal with *k* = 5 clusters. For each selection of the number of clusters in the K-means algorithm, we computed the inter-pattern correlation variability (IPVC) between all the upper triangular parts of the resulting centroids**.**

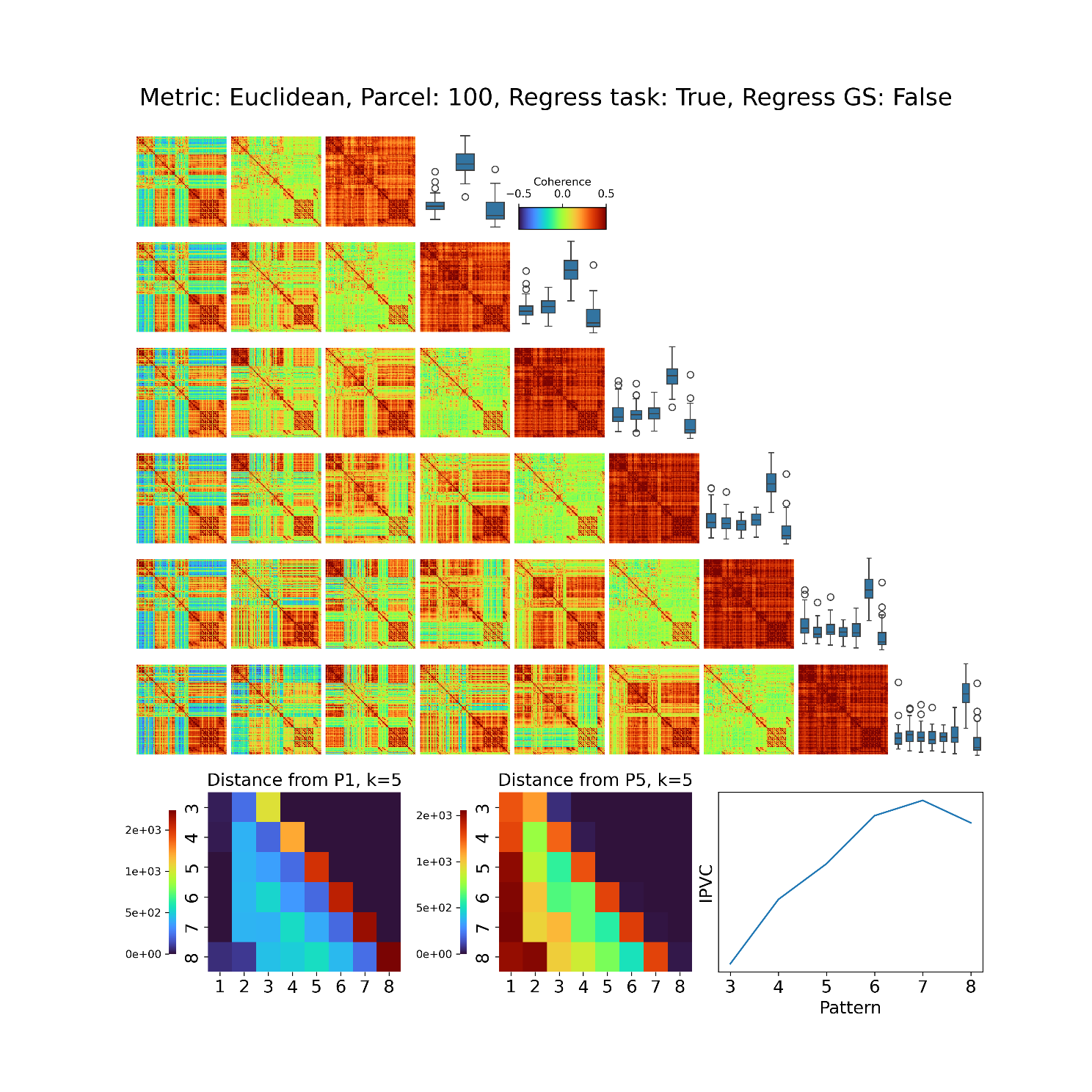

**Supplementary Figure S15.** Recurrent and consistent brain configurations emerge under different clustering dimensionalities during task engagement. Clustering parameters: Metric = **Euclidean** distance, Global Signal Regression: False, Event Regression: True. *Heatmaps:* For each value of *k*, patterns are ordered based on the standard deviation of their connectome, from the most variant (left) to the least (right). *Bottom row – Left*: To examine whether patterns produced across different values of *k* presented correspondence, we estimated the distance of pattern 1 for *k*= 5 with respect to all patterns obtained for *k* = 3 to 8. Similarly, we estimated the distance of pattern 5 for *k*= 5 with respect to all patterns obtained for *k* = 3 to 8. *Bottom row – Right*: The variability of dynamic coordination patterns found by the clustering procedure is maximal with *k* = 5 clusters. For each selection of the number of clusters in the K-means algorithm, we computed the inter-pattern correlation variability (IPVC) between all the upper triangular parts of the resulting centroids**.**

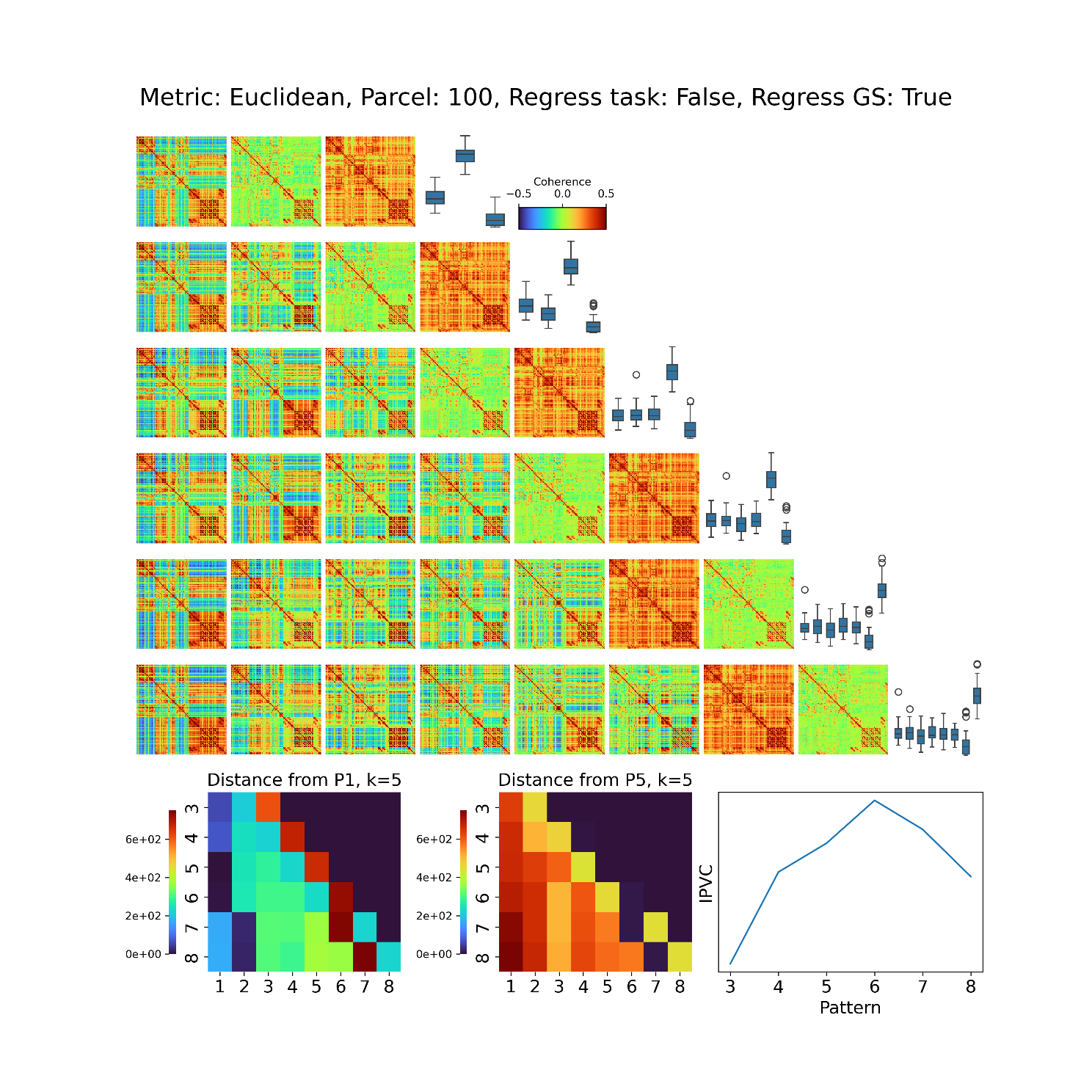

**Supplementary Figure S16.** Recurrent and consistent brain configurations emerge under different clustering dimensionalities during task engagement. Clustering parameters: Metric = **Euclidean** distance, Global Signal Regression: True, Event Regression: False. *Heatmaps:* For each value of *k*, patterns are ordered based on the standard deviation of their connectome, from the most variant (left) to the least (right). *Bottom row – Left*: To examine whether patterns produced across different values of *k* presented correspondence, we estimated the distance of pattern 1 for *k*= 5 with respect to all patterns obtained for *k* = 3 to 8. Similarly, we estimated the distance of pattern 5 for *k*= 5 with respect to all patterns obtained for *k* = 3 to 8. *Bottom row – Right*: The variability of dynamic coordination patterns found by the clustering procedure is maximal with *k* = 5 clusters. For each selection of the number of clusters in the K-means algorithm, we computed the inter-pattern correlation variability (IPVC) between all the upper triangular parts of the resulting centroids**.**

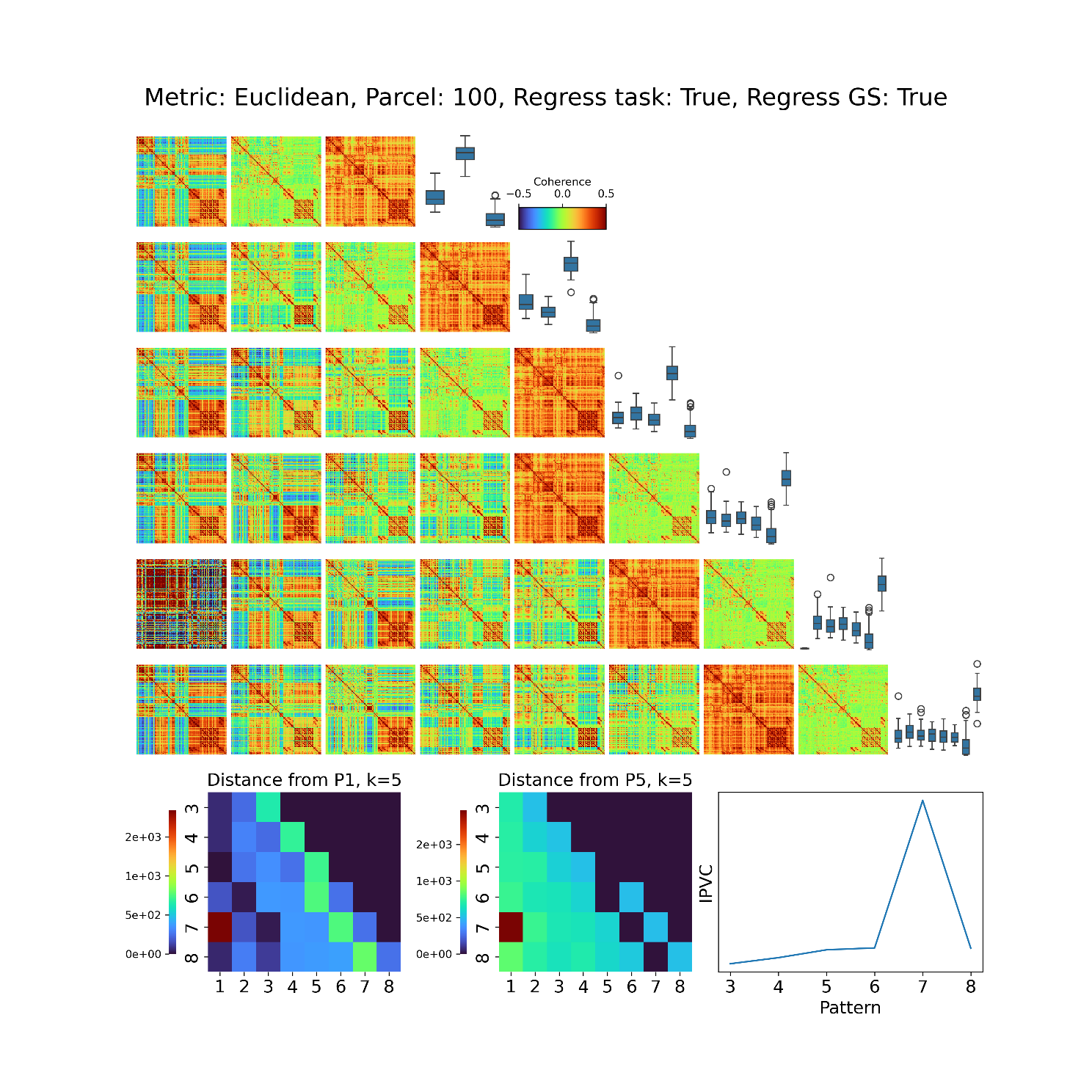

**Supplementary Figure S17.** Recurrent and consistent brain configurations emerge under different clustering dimensionalities during task engagement. Clustering parameters: Metric = **Euclidean** distance, Global Signal Regression: True, Event Regression: True. *Heatmaps:* For each value of *k*, patterns are ordered based on the standard deviation of their connectome, from the most variant (left) to the least (right). *Bottom row – Left*: To examine whether patterns produced across different values of *k* presented correspondence, we estimated the distance of pattern 1 for *k*= 5 with respect to all patterns obtained for *k* = 3 to 8. Similarly, we estimated the distance of pattern 5 for *k*= 5 with respect to all patterns obtained for *k* = 3 to 8. *Bottom row – Right*: The variability of dynamic coordination patterns found by the clustering procedure is maximal with *k* = 5 clusters. For each selection of the number of clusters in the K-means algorithm, we computed the inter-pattern correlation variability (IPVC) between all the upper triangular parts of the resulting centroids**.**

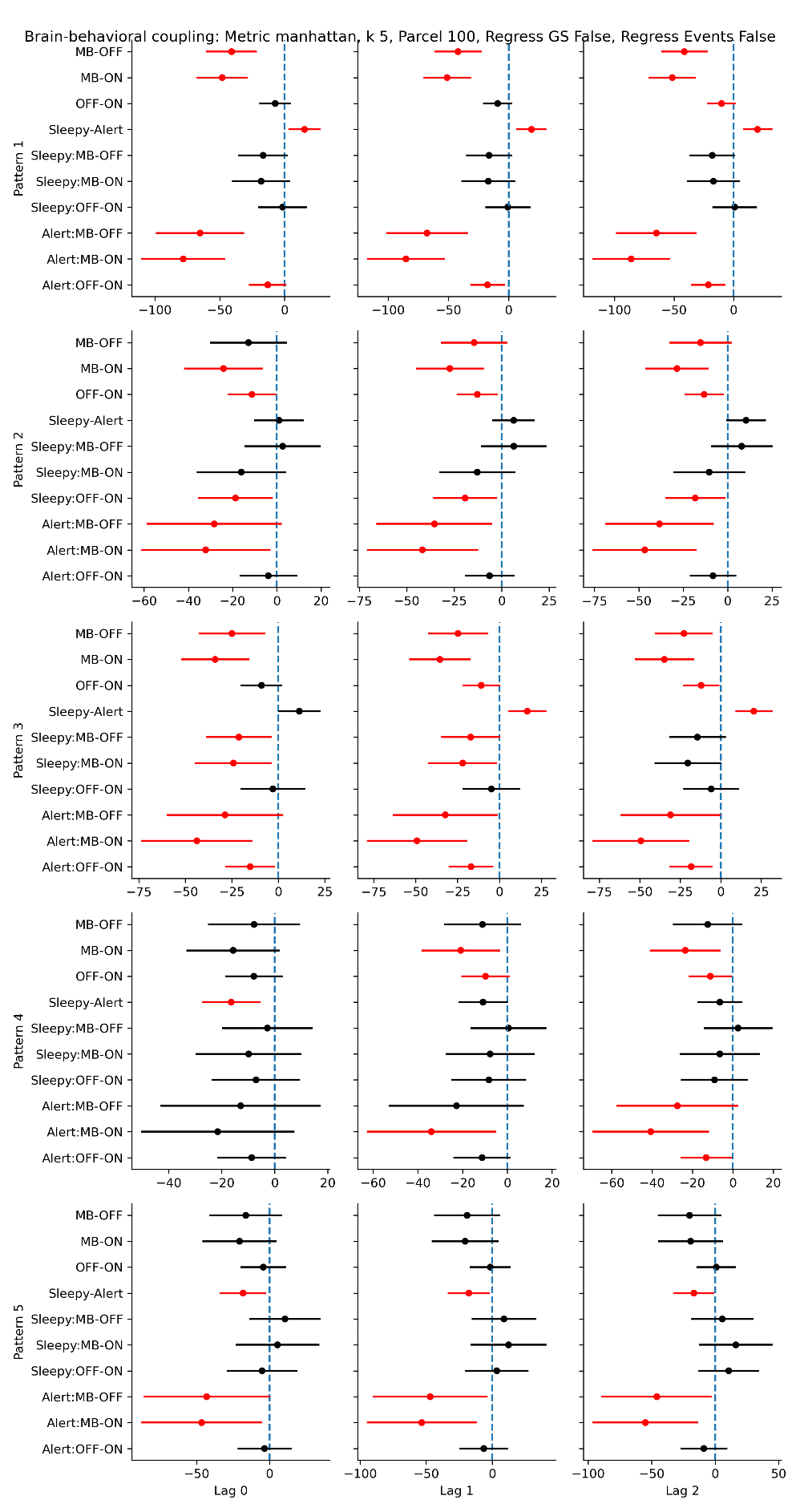

**Supplementary Figure S18.** Replication analysis of the relationship between brain patterns, mental states and alertness. Analysis parameters: distance=Manhattan, k clusters = 5, task regression = **False**, Global signal regression = **False**, N ROI = 100.

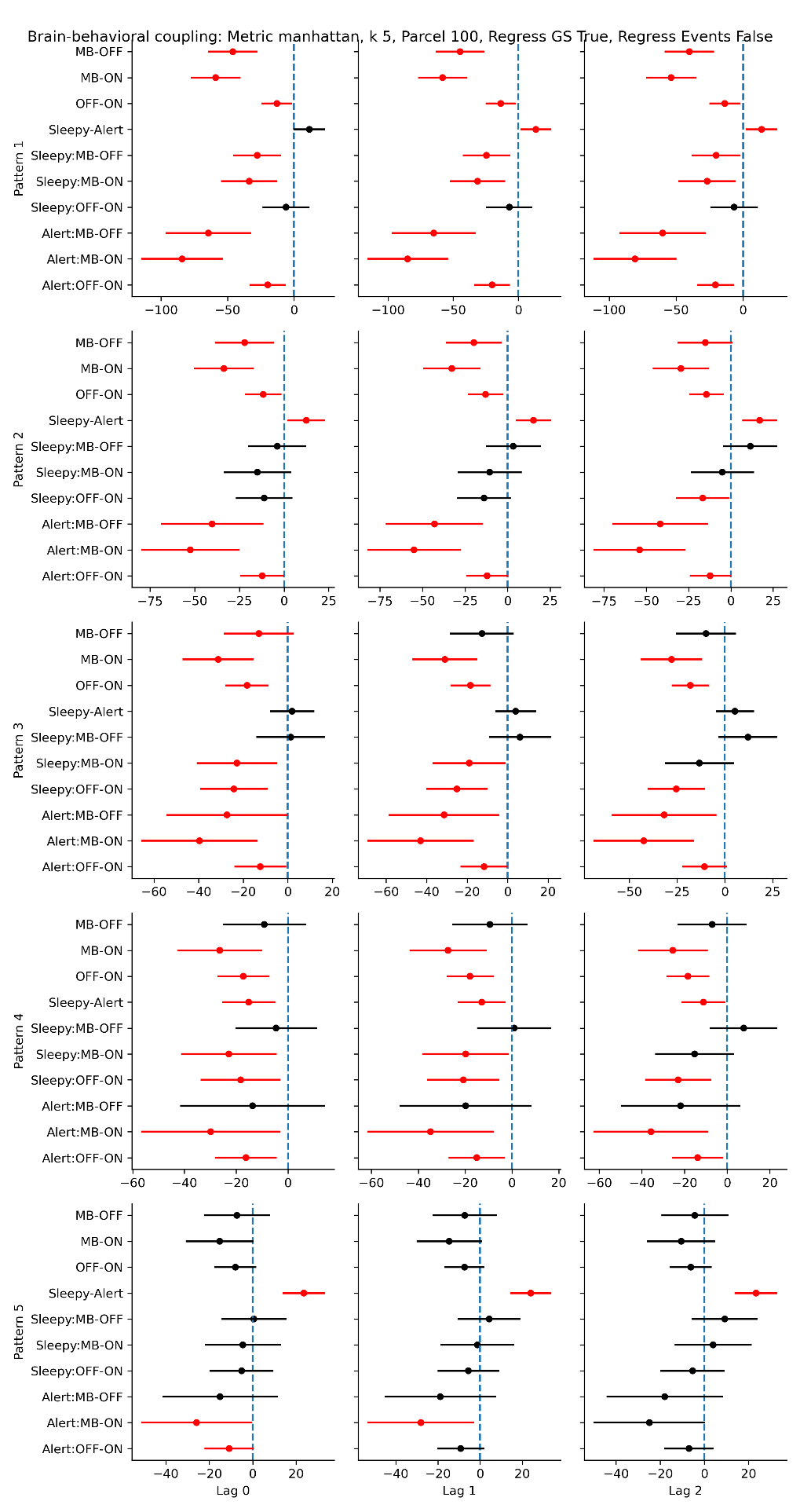

**Supplementary Figure S19.** Replication analysis of the relationship between brain patterns, mental states and alertness. Analysis parameters: distance=Manhattan, k clusters = 5, task regression = **False**, Global signal regression = **True**, N ROI = 100.

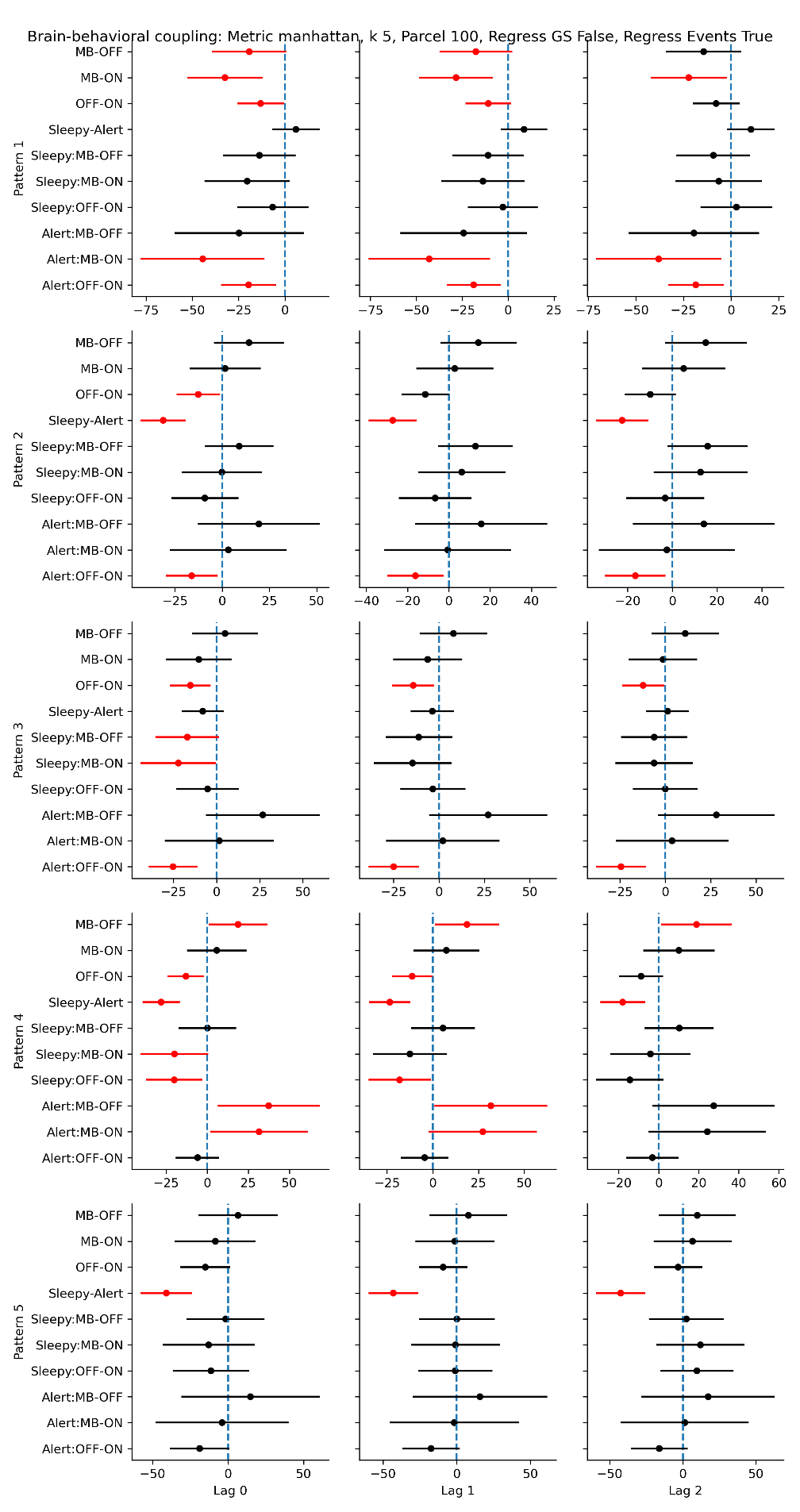

**Supplementary Figure S20.** Replication analysis of the relationship between brain patterns, mental states and alertness. Analysis parameters: distance=Manhattan, k clusters = 5, task regression = **True**, Global signal regression = **False**, N ROI = 100.

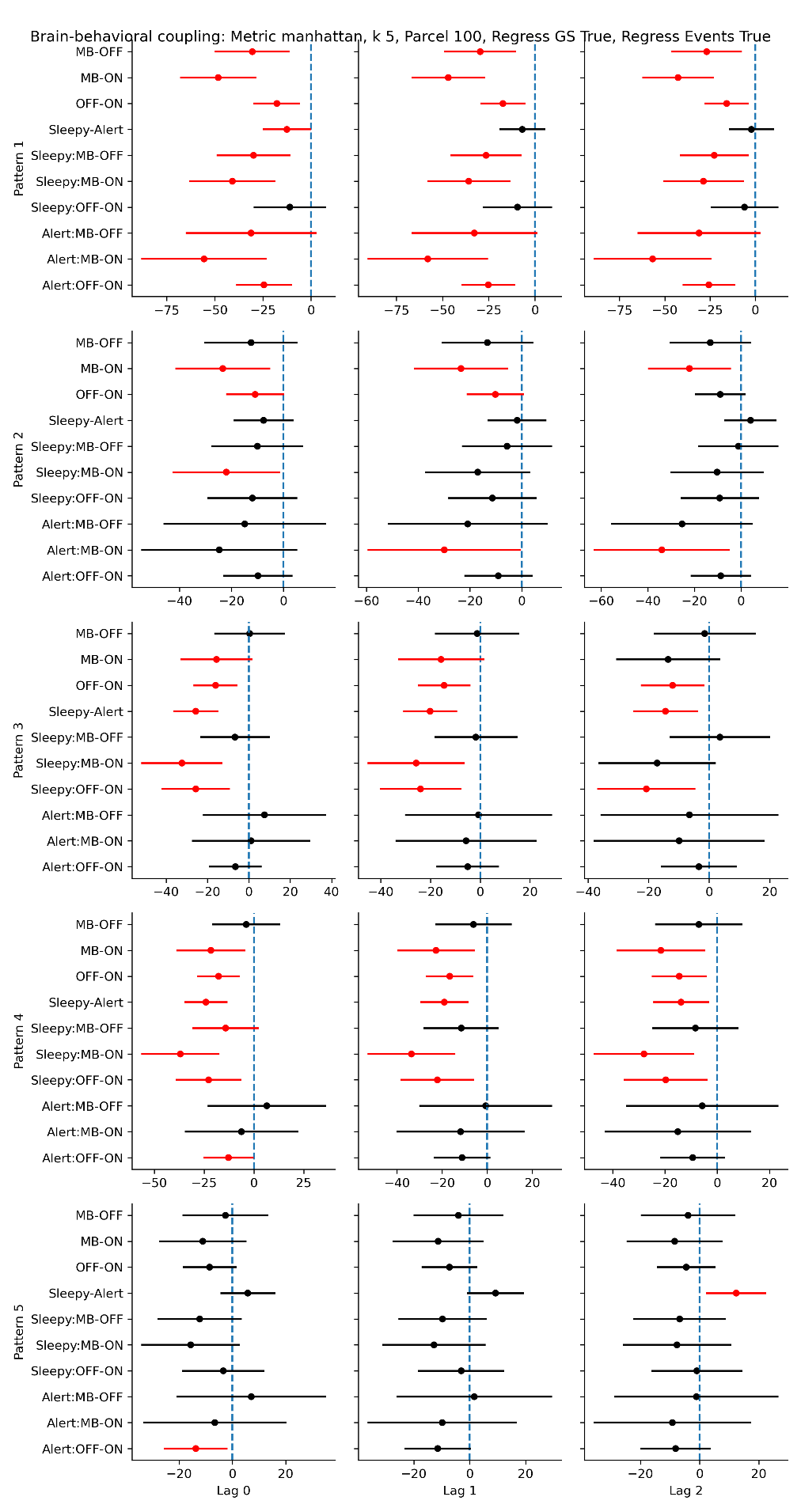

**Supplementary Figure S21.** Replication analysis of the relationship between brain patterns, mental states and alertness. Analysis parameters: distance=Manhattan, k clusters = 5, task regression = **True**, Global signal regression = **True**, N ROI = 100.

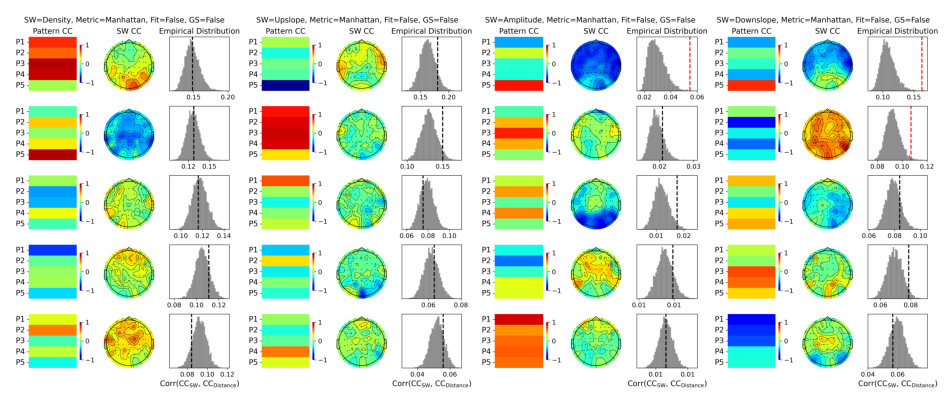

**Supplementary Figure S22.** Replication analysis of the relationship between brain patterns and SW-like activity. Analysis parameters: distance=Manhattan, k clusters = 5, task regression = **False**, Global signal regression = **False**, N ROI = 100.

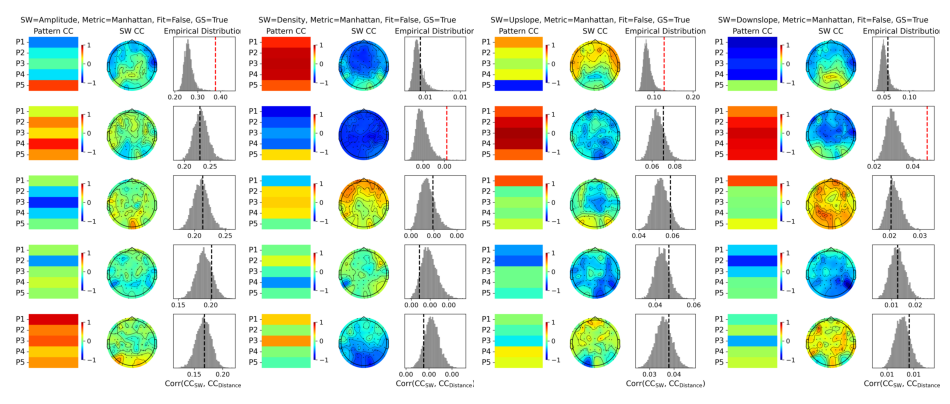

**Supplementary Figure S23.** Replication analysis of the relationship between brain patterns and SW-like activity. Analysis parameters: distance=Manhattan, k clusters = 5, task regression = **False**, Global signal regression = **True**, N ROI = 100.

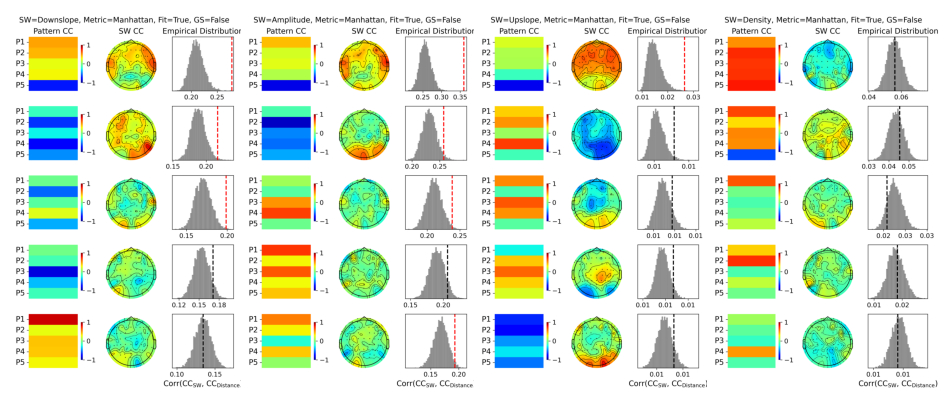

**Supplementary Figure S24.** Replication analysis of the relationship between brain patterns, mental states and alertness. Analysis parameters: distance=Manhattan, k clusters = 5, task regression = **True**, Global signal regression = **False**, N ROI = 100.

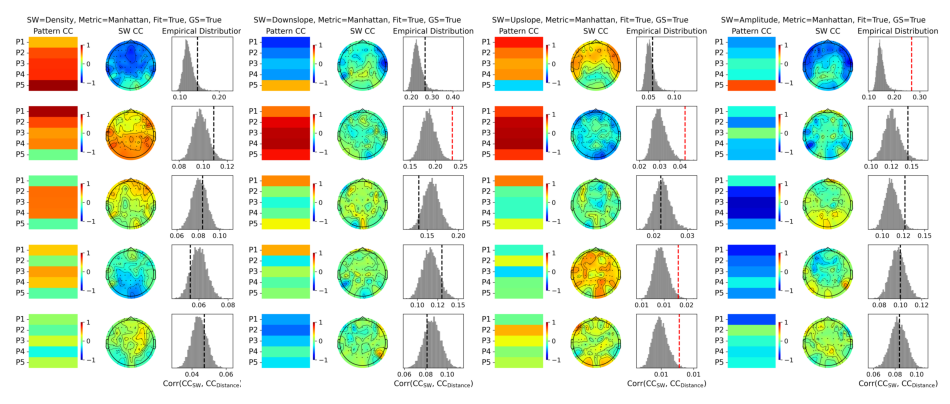

**Supplementary Figure S25.** Replication analysis of the relationship between brain patterns, mental states and alertness. Analysis parameters: distance=Manhattan, k clusters = 5, task regression = **True**, Global signal regression = **True**, N ROI = 100.
